# Persistent autism-relevant phenotype produced by *in utero* and lactational exposure of female mice to the commercial PBDE mixture, DE-71

**DOI:** 10.1101/2021.07.08.451690

**Authors:** Elena V. Kozlova, Matthew C. Valdez, Maximillian E. Denys, Anthony E. Bishay, Julia M. Krum, Kayhon M. Rabbani, Valeria Carrillo, Gwendolyn M. Gonzalez, Gregory Lampel, Jasmin D. Tran, Brigitte M. Vazquez, Laura M. Anchondo, Syed A. Uddin, Nicole M. Huffman, Eduardo Monarrez, Duraan S. Olomi, Bhuvaneswari D. Chinthirla, Richard E. Hartman, Prasada S. Rao Kodavanti, Gladys Chompre, Allison L. Phillips, Heather M. Stapleton, Bernhard Henkelmann, Karl-Werner Schramm, Margarita C. Curras-Collazo

## Abstract

Polybrominated diphenyl ethers (PBDEs) are ubiquitous persistent organic pollutants (POPs) that are known neuroendocrine disrupting chemicals with adverse neurodevelopmental effects. PBDEs may act as risk factors for autism spectrum disorders (ASD), characterized by abnormal psychosocial functioning, although direct evidence is currently lacking. Using a translational exposure model, we tested the hypothesis that maternal transfer of a commercial mixture of PBDEs, DE-71, produces ASD-relevant behavioral and neurochemical deficits in female offspring. C57Bl6/N mouse dams (F0) were exposed to DE-71 via oral administration of 0 (VEH/CON), 0.1 (L-DE-71) or 0.4 (H-DE-71) mg/kg bw/d from 3 wk prior to gestation through lactation. Mass spectrometry analysis indicated *in utero* and lactational transfer of PBDEs (ppb) to F1 female offspring brain tissue at postnatal day (PND) 15 which was reduced by PND 110. Neurobehavioral testing of social novelty preference (SNP) and social recognition memory (SRM) revealed that adult L-DE-71 F1 offspring display altered short- and long-term SRM, in the absence of reduced sociability, and increased repetitive behavior. These effects were concomitant with reduced olfactory discrimination of social odors. Additionally, L-DE-71 exposure also altered short-term novel object recognition memory but not anxiety or depressive-like behavior. Moreover, F1 L-DE-71 displayed downregulated mRNA transcripts for oxytocin (*Oxt*) in the bed nucleus of the stria terminalis (BNST) and supraoptic nucleus, vasopressin (*Avp*) in the BNST and upregulated *Avp1ar* in BNST, and *Oxtr* in the paraventricular nucleus. Our work demonstrates that developmental PBDE exposure produces ASD-relevant neurochemical, olfactory processing and behavioral phenotypes that may result from early neurodevelopmental reprogramming within central social and memory networks.

## Introduction

Autism spectrum disorder (ASD) is a group of neurodevelopmental conditions defined clinically by deficits in social reciprocity and communication, restricted interest and repetitive behaviors (American Psychiatric Association, DSM-V, 2013). Hallmarks of ASD, as classified by the NIH Research Domain Criteria (RDoC)^1^include disturbances in the social cognition (SC) domain such as facial recognition ability, empathy and evaluating emotion of others^2, 3^. The prevalence of ASD has increased dramatically over the past three decades. In the United States, the Centers for Disease Control (CDC) estimates that ASD affects 1 in 54 neurotypical children^4^, while the worldwide prevalence is estimated to be 1-2%^5^. While genetic heritability is an important factor in ASD etiology, the incremental incidence of autism over the last several decades, raises the possibility that environmental factors, such as xenobiotic chemicals, may contribute alongside genetic predisposition and to influence ASD risk^6, 7^. Although the incidence of autism is 4 times more greater in boys, girls and women with autism are often undiagnosed, misdiagnosed or receive a diagnosis of autism at later age^8^ suggesting underestimation in females. According to the female protective model, females may benefit from a higher threshold of genetic liability to manifest ASD phenotype^9, 10^ but may be more susceptible to xenobiotic chemicals^11^ that can potentially influence risk of neurodevelopmental disorders (NDDs). Indeed, we have found that female mice offspring exposed to PBDEs during prenatal and postnatal development exhibit endocrine and metabolic disruption, indicating that females may provide a more susceptible substrate for studying xenobiotic effects on neurodevelopment^12^.

Polybrominated diphenyl ethers (PBDEs) are a class of brominated flame retardants added to a wide range of products including consumer building material, electronics, textiles, plastics and foams including infant products^13^ since the 1970s^14^. Three commercial formulations of PBDEs were prevalent in commerce, including penta-BDE, octa-BDE and deca-BDE. Two commercial PBDE mixtures, penta- and octa-BDEs, were banned in Europe in 2003 and all PBDEs were voluntarily phased out in the US by 2013, leading to a slow, but measurable, decrease in environmental levels as well as in human sera and breastmilk concentrations of some PBDE congeners^15, 16^. Notwithstanding a commitment to voluntary phase out of deca-BDE by 2013, PBDE contamination is predicted to remain an ongoing problem through the next several decades due to their long half-lives, persistence in e-waste^17^, recycling into consumer products and inadvertent reappearance into environment^18^. In an unprecedented action, the U.S. EPA formally banned the production, import and distribution of deca-BDE in February 2021. Nevertheless, PBDEs are still being detected in various tissue samples worldwide, including human breastmilk^19,20,21,22,23^.

Compared to adults, infants and toddlers are at greater risk of the adverse health effects resulting from PBDE exposure since they disproportionately accumulate 3-to-9-fold greater body burdens^24^. Circulating levels of PBDEs in US children are 10-to-1000-fold higher than similar age populations in Mexico and Europe^25^. Elevated exposures in infants are due to the maternal transfer of PBDEs via cord blood and breastmilk^26^. After weaning in early childhood, an additional route of exposure is dust ingestion and inhalation associated with children’s mouthing and crawling behaviors^27, 28^. Therefore, high PBDE exposure poses significant health risks during critical periods of development.

Major health effects associated with PBDE exposures are endocrine disruption, reproductive and developmental toxicity and neurotoxicity^29,30,31,32,33^. Epidemiological studies examining an association between PBDE exposure and ASD show inconsistent findings. PBDE exposure (e.g.., BDE-153 and -47) during both pre- and post-natal development has been linked to adverse neurological outcomes such as impairments in executive function, poor attention and behavioral regulation, reduced social scores, and lower IQ. Early-life l exposure to PBDEs (BDE-47, -99 and/or -100) has been associated with externalizing behaviors such as hyperactivity and impulsivity^34,35,36,37,38^. With regard to the association of PBDEs with social behavior deficits and ASD, preschool-aged children with greater ΣPBDE exposures were rated as less assertive by their teachers^39^ or showed greater anxious behavior^40^. In the HOME prospective cohort study, PBDEs were associated with greater (BDE-28) or fewer (BDE-85) autistic behaviors^41^. Similarly, significantly higher risk of poor social competence symptoms was shown as a consequence of postnatal BDE-47 exposure^42^. Although the possibility that environmental toxicants serve as risk factors for social neurodevelopmental disorders (NDDs) has not been established^43^, PBDEs may have deleterious effects on children’s social development relevant to ASD^35, 41–43, 44^. Studies in experimental animals demonstrate that certain PBDE congeners produce adverse effects on behavior, learning, and memory in exposed offspring^29, 31, 45^ but information about the negative impact of PBDEs on psycho-social behavior is limited^46, 47^. We hypothesized that developmental PBDE exposure produces ASD-relevant behavioral and neurochemical phenotypes in a mouse toxicant model.

Social recognition, or the ability to distinguish between familiar and novel conspecifics, is a fundamental process across species required for forming long-term attachments, hierarchies, and other complex social strategies that enhances survival^48^. Disturbances in this capacity are present in individuals with ASD who have difficulties identifying faces of novel conspecifics from those previously encountered^2, 3^. Rodents, because of their highly social nature, are used as proxies for studying autism-relevant social competence^49^. Mouse social behavior paradigms rely on the natural propensity for investigation of social novelty compared to previously encountered individuals when given the choice^50^. This preference for social novelty has been shown to be absent in mono-genetic, idiopathic and environmental models of ASD^51,52,53^. In the current study, we used a toxicant exposure mouse model to characterize social recognition ability, repetitive behaviors and concomitant autism comorbidities such as anxiety, memory impairment and altered olfactory processing.

While the behavioral deficits in typical ASD rodent models are well established, the underlying neural mechanisms are not well understood. The neuropeptides oxytocin and vasopressin are considered major neurotransmitters implicated in social information processing and social cognition that have shown to bedisrupted in ASD patients^54^. Rodent studies have shown that these neuropeptidergic systems are involved in several social cognition domains such as social memory, social/emotional recognition and social reward^55,56,57,58^. Work by us and our collaborators has provided evidence that PBDEs (and the structural analogues, polychlorinated biphenyls (PCBs)) disrupt the magnocellular neuroendocrine system responsible for vasopressin production involved in osmoregulation, cardiovascular function and social behavior^31, 59,60,61,62,63,64^. We have shown that DE-71 exposure *in utero* and during lactation via maternal transfer can nearly abolish vasopressin immunoreactivity in the activated supraoptic (SON) and paraventricular nucleus (PVN) of the hypothalamus^63^. Therefore, we also tested the hypothesis that PBDEs disrupt gene expression of prosocial neuropeptides such as vasopressin, oxytocin, PACAP and their receptors in regions of the social brain network, which may underlie deficient social behavior^65,66,67^.

To lend insight to whether early-life exposure to PBDEs can produce ASD-relevant phenotypes, we exposed mouse dams to a commercial mixture of PBDEs, DE-71, at low doses to mimic chronic, low-level exposure to BDE congeners and doses encountered by infants and toddlers. We demonstrate that perinatal exposure to DE-71 produces dose-responsive deficient social recognition memory and general memory, altered olfactory function and altered neuromolecular phenotypes in brain regions that coordinate complex social behaviors. To the best of our knowledge, this study is the first to show a comprehensive profile of autistic-relevant behavior and comorbidities in female offspring impacted by maternal transfer of PBDEs. Concomitant characterization of ASD-relevant behavioral and neurochemical phenotypes exhibited by offspring developmentally exposed to and reprogrammed by DE-71, provides an integrative framework for exploring environmental risk factors that may contribute to the increasing incidence of ASD. A portion of our findings has been published in preliminary form^68^.

## Materials and Methods

### Animal Housing and Care

C57Bl/6N mice were generated using breeders obtained from Charles River Labs (West Sacramento, CA). Mice were housed 2-4 per cage in standard polycarbonate plastic cages with corn-cob bedding in a non-specific pathogen free vivarium and kept on a 12:12-h light:dark cycle in a controlled temperature (21.1–22.8°C) and humidity (20-70%) environment. Mice were provided rodent chow and water *ad libitum*. Care and treatment of animals was performed in compliance with guidelines from and approved by the University of California Riverside Institutional Animal Care and Use Committee (AUP#00170026 and 20200018).

### Experimental Design and DE-71 Exposure

DE-71 (technical pentabromodiphenyl oxide; Lot no. 1550OI18A), was obtained from Great Lakes Chemical Corporation (West Lafayette, IN). DE-71 dosing solutions were prepared in corn oil vehicle (VEH/CON) to yield two doses: 0.1 mg/kg/day (L-DE-71) and 0.4 mg/kg/d (H-DE-71) in 2 ml/kg body weight. The DE-71 doses were selected to contain the same molar concentrations of BDE-47 used in mouse studies^46, 69^. BDE-47 is the primary congener in human breast milk^16, 70^.

Offspring were exposed to DE-71 via maternal transfer using a 10-week dosing regimen as described previously (**Fig.1a**)^71^. Mice were randomly assigned to one of three exposure groups: corn oil vehicle control (VEH/CON), 0.1 mg/kg/d DE-71 (L-DE-71) or 0.4 mg/kg/d DE-71 (H-DE-71). This exposure paradigm was chosen to model chronic, low-level exposure to the mother and PBDE transfer to infant during gestation (2nd and 3rd trimester) and lactation as shown in humans^26, 72, 73^. After 3 weeks of pre-dosing, virgin females were paired with an untreated male using harem-style breeding. The presence of a vaginal plug was designated as gestational day (GD) 0. Females that failed to conceive within 10 d were removed from the study. Litters were not culled as justified previously^74^. F1 offspring were weaned and PND21 and housed in same-sex cages (2-4/cage). Dams (F0) and their adult female offspring (F1) were subjected to behavioral testing and later sacrificed by exsanguination via cardiac puncture under terminal isoflurane anesthesia (5%). To reduce cross-over effects, behavioral tests were distributed across three different cohorts. Mice were run through a battery of behavioral tests in the following order for Cohort 1 (mean age): Suok (PND 46); social novelty preference test (PND 71); 3 chamber social novelty (PND 87); elevated plus maze (PND 72). The brains of Cohort 1 were collected at sacrifice on PND 108 used in qRT-PCR. The following tests were performed on Cohort 2: Marble burying (PND 81); olfactory habituation/dishabituation (PND 79); olfactory preference (PND 102); forced swim test (PND 74). Cohort 3 was subjected to social memory recognition (PND 30); juvenile OFT (PND 31); juvenile MB (PND 35); novel object recognition (PND 111) tests. F1 and F0 were tested similarly, except that F0 did not get tested on the Social Recognition Memory Test (SMRT). Analytical characterization by mass spectrometry was performed on brains from Cohort 1 (PND 110) and a subset of Cohort 3 (PND 15). Enzyme-linked immunosorbent assays (ELISA) were performed on plasma from Cohorts 1-3. Whenever possible, the dam was used as the statistical unit of analysis for F1; unless otherwise indicated. In addition, results were replicated in a minimum of 3 independent experiments.

### Nest scoring

To test for possible effects of DE-71 on maternal parameters, nests of single-housed dams built from pressed cotton squares (5 x 5 cm; Nestlets; were evaluated at PND 0-1 using a modified scoring system^75^. Scores were assigned according to the height and closure of the walls surrounding the nest cavity. Scores were assigned if the nest contained a center (1) and a 50% border, (2) 75% border (3) or 100% border (4). A score of 5 was given if the nest resembled a dome (**Supplementary Figure 1**). Nest scores were boosted by 0.5 if the nest was elevated. For interrater reliability the Bland-Altman method was used to calculate bias as the mean of the differences (0 means two judges are not producing different results) and precision as 95% limits of agreement (standard deviation of mean bias +/- 1.96) (**Supplementary Figure 1**).

### Congener Analysis in Adult Offspring Brain

Using gas chromatography coupled with electron capture negative ion mass spectrometry (GC/ECNI-MS; Agilent 5975N MS), PBDE concentrations were measured in PND110 whole brain homogenate extracts by the Stapleton laboratory at Duke University as described previously^71^. Briefly, approximately 0.2-0.5 grams of tissue were first ground with clean sodium sulfate, spiked with two isotopically labeled standards (F-BDE-69 and 13C BDE-209) and then extracted using 50:50 DCM:hexane. Extracts were concentrated, measured for lipid content using a gravimetric analysis, and then purified using acidified silica before analysis for 26 different PBDE congeners ranging from BDE-30 to BDE 209. Laboratory processing blanks (clean sodium sulfate only) were analyzed alongside samples to monitor background contamination. Recoveries of F-BDE-69, and 13C BDE 209, averaged 91 (+/- 6.9%) and 106 (+/- 19.9%), respectively in all samples. All samples were blank-corrected on a congener-specific basis using the average concentrations measured in the laboratory processing blanks. Method detection limits (MDLs) were estimated using either a signal to noise ratio of 10, or, if analytes were detected in laboratory blanks, by calculating three times the standard deviation of the laboratory blanks). MDLs differed by congener and ranged from 0.8 (BDE-47) to 6.6 ng/g (BDE-206).

### PBDE Congener Analysis in Perinatal Offspring Brain

Due to force majeure, i.e. SARS-CoV-2 pandemic, we were unable to carry out planned analytical characterization of PND 15 tissues in collaboration with the Stapleton lab, therefore, the MS system in the Schramm lab was used. The performance of both methods were comparable, especially with regard to the limit of quantification. Using High Resolution Gas Chromatography–High Resolution Mass Spectrometry (HRGC/HRMS), PBDE concentrations were measured in P15 whole brain homogenates (0.1-0.2 g) as described^76^. PBDE analytes included 37 PBDE congeners (BDE-7, 10, 15, 17, 28, 30, 47, 49, 66, 71, 77, 85, 99, 100, 119, 126, 138, 139, 140, 153, 154, 156, 176, 180, 183, 184, 191, 196, 197, 201, 203, 204, 205, 206, 207, 208, 209). Samples were ground and homogenized to a fine powder under liquid nitrogen. Each sample (100-200 mg) was mixed with CHEM TUBE-Hydromatrix (Agilent Technologies) and spiked with ^13^C-labelled PBDE standard mix (BFR-LCS, Wellington Laboratories). For pressurized liquid extraction (Dionex ASE 200) n-hexane/acetone (3:1, v/v) was used at 120°C and 12 MPa. The volume of the extract was reduced to ∼5 mL using a vacuum rotary evaporator. Samples were purified using an automated system (DEXTech, LCTech, Germany), where the sample was passed and fractionated over an acidic silica, alumina and carbon column. Concentrated extracts were spiked with the recovery standard (BFR-SCS, Wellington Laboratories) and analyzed by HRGC/HRMS (Agilent 6890/Thermo MAT95 XL) using electron impact ionization (EI), in the selected ion monitoring mode. The instrumental parameters are listed in **Supplementary Table 1**. Average recovery for ^13^C-labelled PBDE standards ranged between 40 and 120%. All samples were blank-corrected on a congener-specific basis using the average of three procedural blank samples. Analytes with concentrations after blank correction that were lower than three times the standard deviation of the blank values or were not detected before blank correction were considered as not detectable (n.d.). The limit of quantification (LOQ) of the instrumental methodology was considered as a signal/noise ratio of 9:1 (**Supplementary Table 2**). Congener concentrations that were below detection limit were assigned a randomly generated value of LOQ/2. The accuracy of our method was confirmed by successful participation in interlaboratory comparison studies.

### Comparison of MS Methods

The GC/ECNI-MS method used the ECNI ionization mode to improve sensitivity. The latter provides equal sensitivity to HRGC/HRMS that uses electron impact.

### Neurobehavioral Testing Paradigms

At least 30 min prior to testing, mice were moved to a designated behavior room. Ethanol (70%) was used to remove debris and odors between individual mouse trials. Unless stated otherwise, mouse behavior was scored using automated video-tracking software (Ethovision XT 15, Noldus) or manual scoring software (BORIS^77^ or JWatcher), performed blind to treatment by trained observers. Mice were tested between 10am and 4 pm during the light phase under bright light conditions, unless otherwise stated.

### Social Novelty Preference

Social novelty preference (SNP) was conducted and analyzed according to methods adopted from published protocols^50^. Briefly, mice were habituated for 30 min to a polycarbonate cage identical to their home cage (27 x 15 cm), followed by 30 min to two wire interaction corrals (11 x 10 x 10 cm) placed on each side of the cage. During a 5-min training trial, a stimulus mouse was placed into one corral while the empty corral was removed. After a 30 min retention period, social recognition was assessed in the following 5 min test, during which the test mouse explored the same stimulus mouse (now familiar mouse) versus a novel stimulus mouse. Prior to testing days, sex- and age-matched conspecific stimulus mice were trained to stay in corrals for 15 min x 3/d for 7-14 d. Stimulus mice were single-housed in order to preserve their unique scent. Investigation by test mouse was measured as time spent sniffing (snout within 2 cm of stimulus). Test robustness was measured using an Investigation Index calculated as the ratio of time spent investigating the novel mouse to total investigation time during training period. (**Supplementary Figure 2**). Social recognition is represented as time spent investigating novel stimulus as percent of total investigation time in the test period. To evaluate between group differences, a Discrimination Index was calculated as the ratio of time spent investigating Novel - Familiar/total investigation time in test period.

### Three-Chamber Sociability Task

Sociability was assessed as described^78^. In brief, during the first habituation phase, test mice were habituated for 10 min to the center chamber of a Plexiglass three-chambered apparatus (22 x 40 x 23 cm). Next, the retractable doors partitioning the chambers were opened to permit exploration of all three chambers (second habituation phase). Sociability was tested in the following 10 min session, during which the test mouse was permitted to explore an empty 9 x 27 cm corral (novel object) versus a mouse placed inside another corral (novel social object). Inherent side preference during the second habituation phase was evaluated as Right Chamber time - Left Chamber time / Right Chamber time + Left Chamber time x 100. Mean values for test mice meeting the exclusion criterion (0 +/- 15%) are shown in **Supplementary Figure 2**. Sociability was analyzed during the subsequent testing phase both as time spent in chamber and time spent sniffing within 2 cm of stimulus.

### Marble burying and Nestlet Shredding Tests

The Marble Burying (MB) and Nestlet Shredding Tests were utilized for analysis of elicited repetitive behavior in rodents that is considered analogous to those observed in autistic individuals^51^. During the marble burying test, the test mouse was placed in the corner of a polycarbonate cage (19 x 29 x 13 cm) containing 5 cm of bedding^79^ allowed to interact for 30 min with an array of equidistantly aligned marbles (8 x 4 for adults or 6 x 4 for juvenile). A minimum ⅔ of the marble was defined as being buried in the 32-marble array and ½ buried in the 20-marble array. Images of the cage were scored by 2-3 investigators who were blind to treatment and a mean score obtained. For interrater reliability on marble burying the Bland-Altman method was used to calculate bias as the mean of the differences (0 means two judges are not producing different results) and precision as 95% limits of agreement (standard deviation of mean bias +/- 1.96) (**Supplementary Figure 3**). After a 5 min rest period, the test mouse was placed into another cage of the same size with 0.5 cm of bedding containing a pre-weighted square of cotton fiber (Nestlet). After 30 min, the remaining Nestlet was weighed and percent shredding calculated.

### Innate Olfactory Preference Test

To test the ability to detect attractive or aversive odorants, the innate Olfactory Preference Test (OPT) was performed and analyzed according to methods described^80^. Mice were habituated to the experimental conditions by being placed individually into an empty test cage (19 x 29 x 13 cm) and sequentially transferred to three other cages every 15 min. After the final habituation, mice were transferred into the test cage containing a filter paper (2 x 2 cm) infiltrated with 500 uL of a fresh sample of test odorants: 10% peanut butter, 1% vanilla, 1% butyric acid, or deionized water. The four test odorants were presented to the test mouse in a randomized order. Time spent sniffing the filter paper during the 3-min odorant trials was video-recorded and later measured. Cages were cleaned with 70% ethanol after each mouse was tested.

### Olfactory Habituation Test

The ability of mice to detect and differentiate social and non-social odorants was examined using the Olfactory Habituation/Dishabituation test (OHT)^51^. OHT involves presenting a test animal with various odorants, typically: (1) water, (2) two non-social odorants (almond and banana) and (3) two social odorants (obtained from cage bedding). Mice were acclimated for 45 min to an empty cage with a cotton-tipped applicator inserted through the water bottle hole in order to reduce the novelty of the applicator during test sessions. Non-social odors were prepared from extracts immediately before testing. They included: (1) deionized water; (2) almond (1:100 dilution); (3) banana (1:100) (McCormick). Two social odors were obtained the morning of test day by swiping applicator across the bottom of stimulus cages containing soiled bedding from sex-matched conspecifics. Cages housed 3-4 mice and bedding was at least 3 d old. Stimuli were presented in 2-min trials in the following order: water x 3, almond x 3, banana x 3, social odor 1 x 3, social odor 2 x 3. Time spent sniffing the applicator was recorded with a stopwatch. Parameters measured were habituation, defined as a decrement in olfactory investigation of the same odor after repeated presentations and dishabituation, defined as a reinstatement of olfactory investigation upon presentation of a new odorant.

### Social Recognition Memory Test

A two-trial social recognition memory test (SMRT) was performed as previously described^66^ to test assess long-term social recognition memory. Test mice (PND 28-40) were exposed to a juvenile sex-matched conspecific stimulus mouse (PND 15-32) during two 3 min trials following an intertrial delay of 24 h. For each experiment, test mice were individually placed into polycarbonate cages (27 x 16 x 12 cm) and allowed to habituate for 1 h under dim conditions. A juvenile sex-matched conspecific was then placed into the cage, and the mice were allowed to interact for 3 min (Trial 1). In Trial 2, performed 24 h later, the same test mouse was exposed to either the familiar (stimulus from Trial 1) or a novel stimulus. Each stimulus was not used more than 4 times. The tests were digitally recorded and scored for social investigation behavior. To evaluate the differences in ability to form a long-term social memory a Recognition Index was calculated as the ratio of the duration of investigations on Day 2 and Day 1.

### Novel Object Recognition Test

The novel object recognition test (NORT) was used to assess non-social recognition memory. We adapted a two-day protocol with a short- and long-term retention time. On Day 1, the test mouse was habituated to an empty square Plexiglas open field arena (39 x 39 x 38 cm) for 15 min as described^81^, followed by a 20 min rest in home cage. During the acquisition phase, the test mouse was placed in the open field containing two identical objects (F vs F’) and allowed to freely explore the environment and objects. During the short-term memory (30 min retention) testing session, the test mouse was again placed in the apparatus and allowed to explore a familiar and novel object (F vs N). After a 24 h retention time (Day 2), long term memory was assessed by placing mice into the open field containing both the familiar and a new novel object (F vs N’). All test/train sessions lasted 5 min. Preference for the novel object was expressed as the ratio of time exploring the novel relative to the total exploration time. To evaluate the differences in ability to form NOR memory, the Discrimination Index was calculated as the ratio of the difference in exploration time between novel and familiar objects relative to total exploration time, where 0 indicates equal preference. Test objects were first validated for intrinsic preference. After analysis of the data using Ethovision, the exclusion criteria was the lack of travelling a distance within one standard deviation of the group mean for trials 1 and 2 and/or any animal not visiting the familiar or novel target zones at least 6 times.

### Suok

Suok is an elevated platform behavioral paradigm used to analyze anxiety, anxiety-induced motor impairments and motor-vestibular anomalies in mice. The apparatus consists of a smooth aluminum rod (2 m long, 3 cm diameter) elevated to 20 cm and fixed to two clear acrylic walls as described^82^. Bilateral to a central segment (38 cm) of the aluminum rod are 10 cm segments labeled by line markings. After acclimation to the dimly lit testing room, several behaviors are scored over a 5 min trial: (1) horizontal and locomotor (normalized) activity, assessed by number of segments traveled, (2) sensorimotor coordination -measured by the number of hind leg slips and falls from the rod, (3) exploratory behavior like side looks and head dips, (4) anxiogenic behaviors like increased latency to leave the central zone and unprotected stretch attend postures (SAP), in which the mouse stretches forward and retracts without moving its feet-considered a non-social form of ambivalence, (5) vegetative responses (combined number of urinations and defecation boli), and (6) autogrooming behaviors. Hyperactivity, loss of sensorimotor coordination, increased anxiety and displacement behavior are represented by elevated values for #1, 2, 4 and #5, and 6. Measures were recorded manually by stopwatch. Locomotor activity was calculated as total test time minus time spent immobile in center.

### Open Field Test

The open field test allows rapid assessment of rodent locomotion, anxiety and habituation without a training requirement^83^. The open field apparatus, a Plexiglas square arena of 39 x 39 x 37.8 cm was designed as a large, brightly lit, open and aversive environment. Locomotor and other activity over a 1 h period was digitally recorded and scored using Ethovision for distance traveled, velocity and total time in periphery (10 cm adjacent to wall) and center.

### RNA Extraction From Brain Micropunches

At sacrifice under isoflurane anesthesia whole brains were rapidly dissected and snap frozen in 2-Methylbutane over dry ice. Brains were cryosectioned (0.3 mm thick) coronally and sections mounted on sterile glass slides and stored at -80^0^C. Five regions of interest were punched out bilaterally from tissue sections under a stereomicroscope using a microdissecting needle (16-gauge) adapted from the Palkovits micropunch technique^84^. The anatomical precision was determined based on the atlas of Paxinos and Franklin and cresyl violet stained sections of reference mouse brains. Tissue punches were immediately homogenized in TRIzol Reagent (Thermo Fisher Scientific, USA) using a hand-held homogenizer. Total RNA was prepared via a modified partial phenol-methanol extraction protocol (RNeasy Micro Kit, Qiagen, USA). Purity and quality of RNA were assessed by determining the optical density (OD) photometrically at 280 nm and 260 nm (NanoDrop ND-2000, Thermo-Fisher Scientific Inc., Waltham, MA, USA). RNA integrity was assessed using an Agilent 2100 Bioanalyzer (Agilent Technologies Inc. Santa Clara, CA, USA) (**Supplementary data 2**).

### Quantitative polymerase chain reaction (qRT-PCR)

RT-qPCR was used to quantitate mRNA transcripts for pro-social peptides, AVP, OXT, PACAP and their receptors. Oligonucleotide PCR primers were custom designed and synthesized or ordered as predesigned assays from Integrated DNA Technologies. Primers were designed to meet several criteria using NCBI Primer Blast and then optimized by testing against complementary DNA generated using RT-PCR and gel electrophoresis. Only primers that gave single-band amplicons in the presence of RT and that matched the base length of the predicted target and which yielded 90% to 110% efficiency were selected (**Table 1**). *Oxtr* and the reference gene, *ActB*, were multiplexed using hydrolysis probes with double-quenchers. For all other primers, intercalating dye chemistry was used. RT-qPCR was performed on a CFX Connect thermocycler with the Luna Universal or Probe one-step qPCR Master Mixes (New England Biolabs, Ispwich, MA). RNA (1-4 ng) was used per reaction run in triplicate. In each experiment, no-template controls (NTCs) without mRNA were run to rule out extraneous nucleic acid contamination and primer dimer formation. Negative RT controls, which contained the complete RNA synthesis reaction components without the addition of the enzyme reverse transcriptase (RT) were used to rule out presence of genomic DNA (gDNA). Fold-change gene expression was measured relative to the reference gene, *ActB*, and differential gene expression was determined compared to null group (VEH/CON) using the Pfaffl method. Molecular work was carried out in adherence to MIQE guidelines^85^ (**Supplementary data 2**).

**Table 1.**
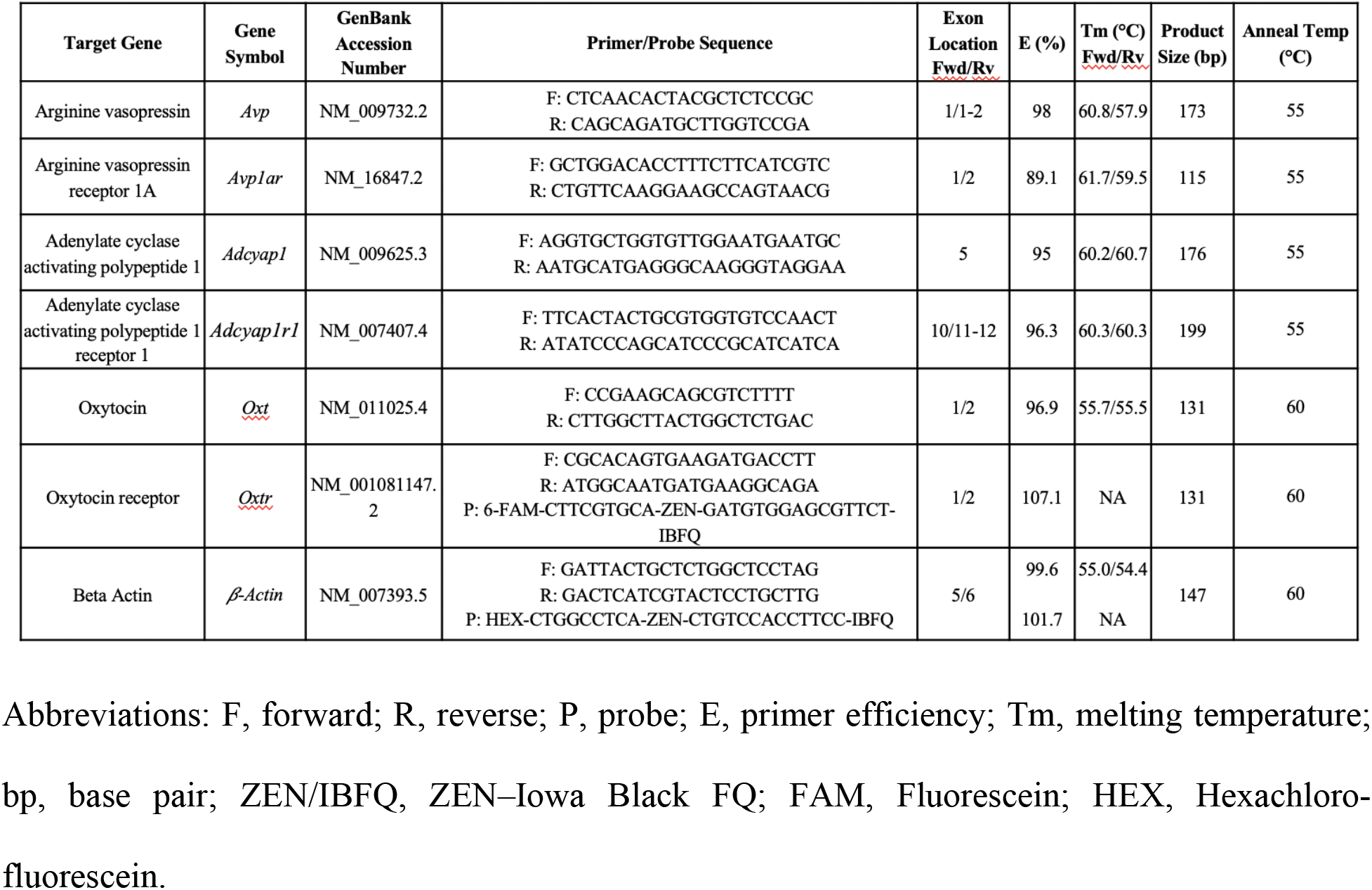
RT-qPCR Primers and Target Genes.

### Enzyme Immunoassays

Blood was collected by cardiac puncture and the plasma separated at 2000 x g centrifugation for 20 min at 4℃. Plasma levels of the neuropeptides OXT and arginine8- vasopressin (Arg^8^) were quantified using commercially available ELISA kits from Arbor Assays (Ann Arbor, MI USA, OXT, K048-H1, Arg^8^, K049-C1) and Enzo Life Sciences (OXT, ADI901153A0001, Arg^8^, ADI-900-017) following the manufacturer’s instructions. For the Arbor Assay kits, in order to reduce the non-specific binding, samples were first treated using the acetone-based extraction solution followed by vacuum lyophilization of the resulting supernatant. For oxytocin, the colorimetric reaction product was read as optical density at 450 nm on a plate reader (SpectraMax 190, Molecular Devices). The kit has a sensitivity of 1.7 pg/sample in a dynamic range of 16.38-10,000 pg/mL. ARG^8^ was detected using a luminescence plate reader (Victor3, Perkin Elmer). The ARG^8^ kit has a sensitivity of 0.9 pg/mL in a dynamic range of 1.638-1,000 pg/mL. For the Enzo Life Sciences kits, samples underwent solid phase extraction using 200 mg C18 Sep-pak columns as previously described^86^. Plasma oxytocin and arginine vasopressin were quantified by interpolating absorbance or luminosity values, respectively, using a 4-parameter-logarithmic standard curve (MyAssays).

### Statistical Analyses

Statistical analysis was performed using GraphPad Prism (version 8.4.3 San Diego, CA, USA). Within groups comparison was performed using paired Student’s t test or one-way ANOVA if more than two groups were compared. Between groups comparisons were accomplished using One-way, Two-way or Mixed model ANOVA with or without a repeated measures design. Non-parametric statistical tests (i.e., Kruskal-Wallis H test) were used when normality and/or equal variances assumptions were not met as measured using the Shapiro-Wilk and F-tests. If an equal variance assumption was not met, a Brown-Forsythe ANOVA or Welch’s correction was used. Post-hoc comparisons were performed using appropriate tests. Technical outliers were excluded when animals were unable to perform behavioral tests. Type 1 error rate (α) was set at 0.05; F and *P* values are presented in the figure legends or Supplemental statistical information. The data are expressed as the mean ± s.e.m, as mean with individual values as ‘before-after’ bars or as median and inter quartile range representing minimum and maximum values in whisker plots.

## Results

### DE-71 Dosing Paradigm and Maternal Parameters

C57Bl/6 mice dams were exposed to DE-71 and later investigated along with the F1 female offspring (**Fig. 1**). Using this dosing paradigm, we have previously reported no differences in litter size at birth, secondary sex ratio, nor gestational maternal parameters^71^. Dams exposed to DE-71 did not build inferior nests and F1 litters at PND 46 had normal body mass relative to VEH/CON (**Supplementary Figure 1**). In combination, these data indicate that perinatal DE-71 exposure does not interfere with pup health, maternal nest quality nor related behaviors shown to be affected by exposure to PCBs, a structural/functional analogue class of PBDEs^87^.

**Fig. 1.**
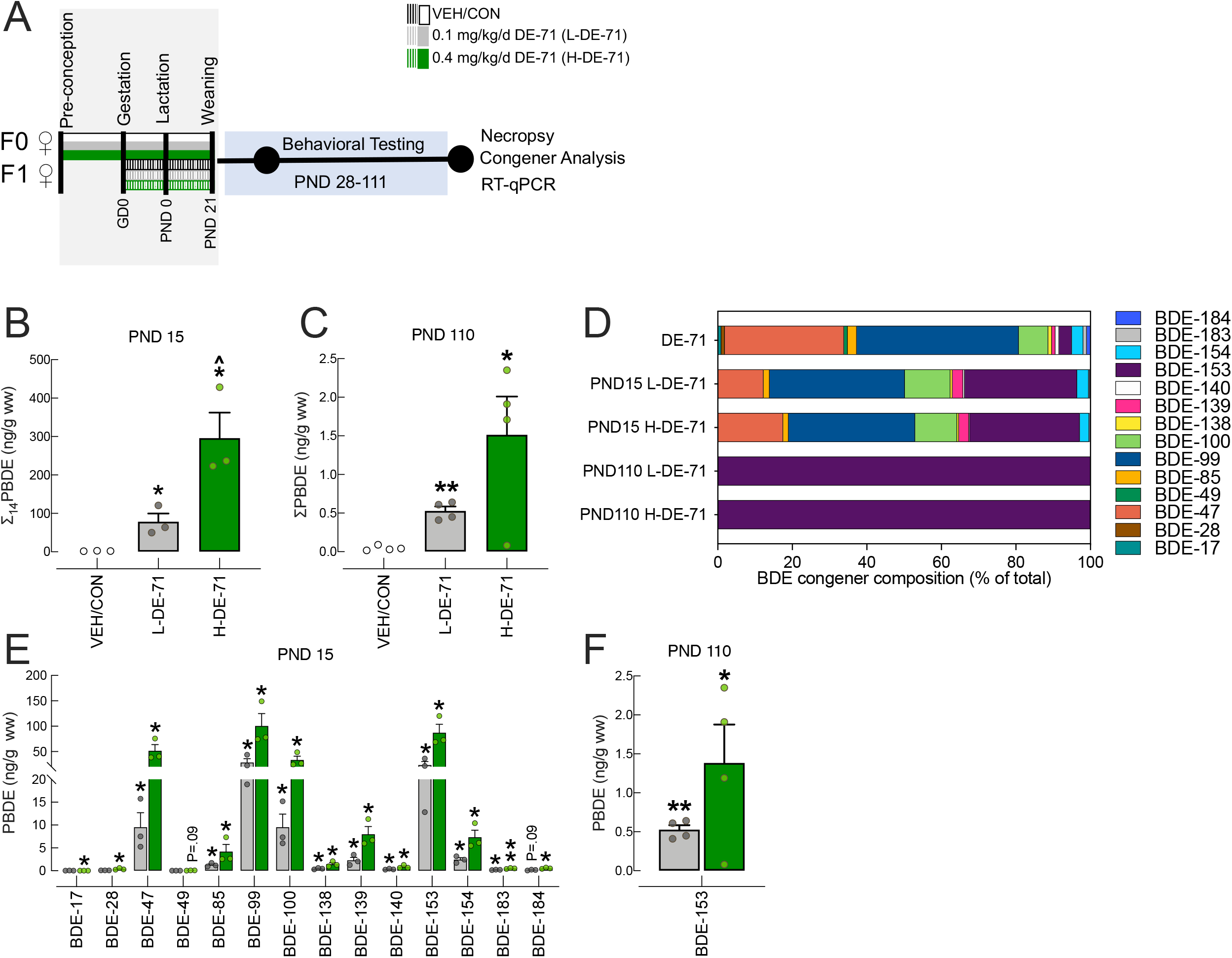
Maternal dosing paradigm for DE-71 produces BDE congener penetration in female F1 offspring brain. **a** Dosing and testing paradigm used for perinatal and adult exposure to DE-71. Direct exposure to DE-71 in adult dams (F0♀; solid shading), began ∼3-4 weeks pre-conception and continued until pup weaning at PND 21. Indirect exposure in female offspring (F1♀; hatched shading) occurred perinatally (GD 0 to PND 21). **b** The ng/g wet wt (ww) sum concentrations of the 14 PBDE congeners (∑_14_PBDE) detected at PND 15. **c** The ng/g wet wt sum concentrations of the 1 PBDE congener, BDE, 153, detected at PND 110. **d** BDE composition (% total) in DE-71 and in brains of exposed female offspring obtained at PND 15 and PND 110. The 7 congeners that comprise <1% of DE-71 were displayed as 1%. **e,f** Absolute congener concentrations at PND 15 and PND 110 for L- and H-DE-71. All values for VEH/CON were <MDL (not shown). \**P*<.05, \*\**P*<.01 compared to VEH/CON; ^*P*<.05 compared to L-DE-71. *n*=3-4/group. GD, gestational day; PND, postnatal day

### PBDE congener analysis in offspring brain

PBDE congener content was determined using HRGC/HRMS or GC/ECNI-MS in F1 female brain from offspring during the lactational period (PND 15) or adults (PND 110), respectively. Raw values are listed by exposure group in **Supplementary Table 3, 4, 5**. **Figure 1 b,c** shows a significant increase in ∑PBDEs in L-DE-71 and H-DE-71 relative to VEH/CON (*P*<.05), confirming that the dosing regime led to maternal transfer of PBDEs. Accumulation of PBDEs in PND 15 (but not PND 110) was dose-dependent (*P*<.05). Mean ∑_14_PBDE values in exposed F1 at PND 15 were 78 and 296 ng/g w.w. for L-DE-71 and H-DE-71, respectively. At PND 110 the corresponding mean total PBDEs (of which only BDE-153 was above detection limits) were 0.53 and 1.5 ng/g w.w. and 113-169 ng/g when normalized to lipid weight (l.w.). For PND 15 the range of BDE concentrations in L-DE-71 and H-DE-71 were as follows, respectively: BDE-17 (0.021, 0.006%), BDE-28 (0.088, 0.126%), BDE-47 (12.2, 17.4%), BDE-49 (0.014, 0.017%), BDE-85 (1.63, 1.41%), BDE-99 (36.3, 34.0%), BDE-100 (12.2+, 11.3%), BDE-138 (0.572, 0.488%) BDE-139 (2.90, 2.70%), BDE-140 (0.408, 0.288), BDE-153 (30.2, 29.5%), BDE-154 (3.12, 2.48), BDE-183 (0.244, 0.168), BDE-184 (0.185, 0.182%) (**Fig. 1d**). Collectively, 7 congeners (BDE-47, -85, -99, -100, -139, -153, -154) in L-DE-71 and H-DE-71 accounted for 98.5, 98.7%of all PBDEs penetrating the brain during lactation, respectively. These same 7 congeners comprise 97.1% of the DE-71 mixture. The remaining 7 of 14 congeners detected in our samples, made up the remaining 1.5, 1.3%, respectively: BDE-17, 28, 49, 138, 140, 183 and 184. **Figure 1e** shows that, with the exception of BDE-17, 28, 49 and 184 in L-DE-71 and BDE-49 in H-DE-71, all 14 congeners detected showed significantly elevated concentrations in DE-71 exposed PND 15 offspring relative to VEH/CON (*P*<.05 -.01). Of note, BDE-153 was ∼10-fold enriched and BDE-47 was slightly depleted (∼2-fold) relative to the DE-71 mixture as reported previously^32^.

By PND 110, the BDE composition in F1 brain was limited to BDE-153 (**Fig. 1f**), which was significantly elevated in L-DE-71 and H-DE-71 relative to VEH/CON (*P*<.01 and *P*<.05). BDE-153 at ppb (and an additional 6 congeners) is detectable in *postmortem* brain samples from 4-71 year-old neurotypical controls and autistic humans born in 1940 to 2000^88^.

### Early-life exposure to DE-71 produces deficits relevant to core symptoms of autism

#### Social novelty preference

Testing mice on a social novelty preference (SNP) test has been suggested to be ethologically relevant to symptoms observed in autistic behavior^50^. On this test, all F1 exposure groups except the L-DE-71 F1 group (*P*<.05) showed a preference for the novel over familiar stimulus (**Fig. 2a**), and this was also represented in the recognition index vs VEH/CON (**Fig. 2b**, *P*<.05). In contrast, there was no effect of exposure in F0 and all groups showed a preference for novel stimulus (**Fig. 2c**, *P*<.0001) and no differences were observed in recognition index (**Fig. 2d**). The investigation index for F1 and F0 approached 1 (**Supplementary Figure 2**) indicating that the reduced exploration of novel over familiar shown by L-DE-71 F1 was not due to a decrease in total investigation time indicating no lack of participation.

**Fig. 2.**
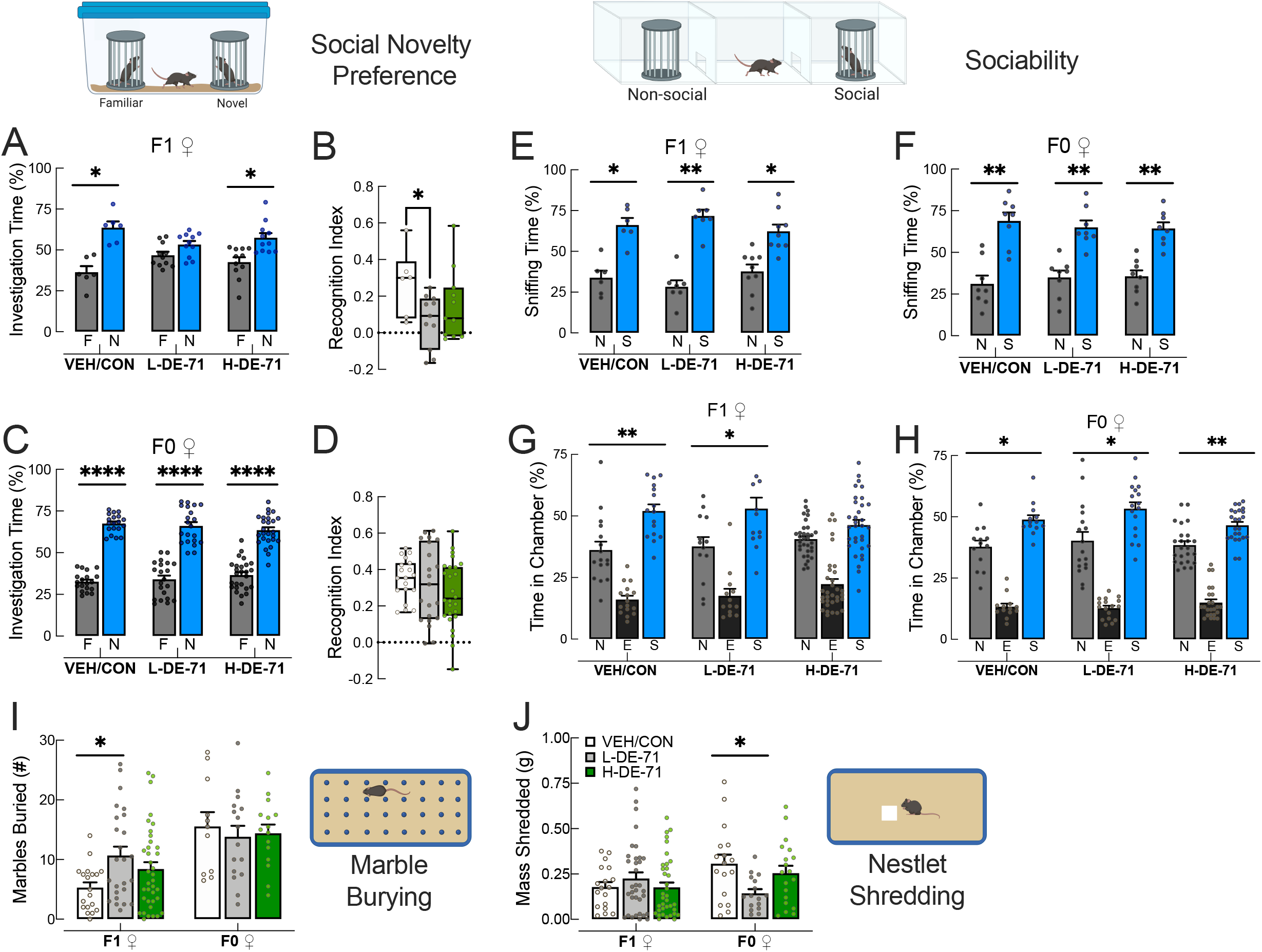
Early-life exposure to DE-71 produces deficits relevant to core symptoms of ASD. **a, c** Social Novelty Preference scores for dams and female offspring: unlike the F1 VEH/CON and F1 H-DE-71, F1 L-DE-71 females failed to spend more time with a novel relative to a familiar conspecific stimulus. F0 dams exposed to DE-71 did not show abnormal social recognition relative to VEH/CON. **b, d** Recognition Index scores show decreased preference for novel stimulus in L-DE-71 F1 relative to VEH/CON but not in F0. **e, f** Time spent sniffing in Sociability test. All exposure groups spent significantly more time sniffing social stimulus indicating normal sociability. **g, h** Chamber time scores in Sociability. All groups show significantly greater time spent in social chamber relative to non-social except for F1 H-DE-71. **i** Marble Burying scores showed offspring L-DE-71 buried a greater amount of marbles as compared to VEH/CON and H-DE-71, but not in dams. **j** Nestlet Shredding was not affected in exposed F1 but was reduced in L-DE-71 F0 relative to corresponding VEH/CON. \**P*<.05, \*\**P*<.01; \*\*\*\**P*<.0001 compared to VEH/CON (b,d,i,j), familiar (a,c) or non-social chamber (e,f,g,h). *n*=6-11 litters/group (a-b), 19-26 subjects/group (c-d), 6-9 litters/group (e), 8 litters/group (f), 13-33 litters/group (g), 13-24 subjects/group (h), 19-37 subjects/group for F1 and 11-16 subjects/group for F0 (i), 18-36 subjects/group for F1 and 16-19 subjects/group for F0 (j). F, familiar, N, novel; N, non-social, S, social; N, non-social, E, empty, S, social

#### Sociability

To determine social interest, an independent social cognition domain, we examined mouse behavior on a 3-chamber sociability test. All F1 groups (VEH/CON, L-DE-71, H-DE-71) showed preference for a novel social stimulus relative to a non-social novel stimulus as measured by sniffing time (**Fig. 2e**, *P*<.05, .01, .05), respectively, indicating normal sociability. Using chamber time VEH/CON and L-DE-71, but not H-DE-71 F1, showed a preference for social stimulus (**Fig. 2g**, *P*<.05, *P*<.01, ns). Sniffing time has been suggested to have superior validity over chamber time scores since active behaviors that are most directly related to social investigation are captured^89^ since the physical proximity allows for transmission of volatile and nonvolatile oderants^48, 89, 90^. For F0, chamber **(Fig. 2f**, *P*<.01) and sniffing time scores were congruent and no effect of exposure was found (Fig. **2h** *P*<.05 for VEH/CON and L-DE-71, and *P*<.01 for H-DE-71). As a measure of test robustness there was no indication of side preference during training for F1 and F0 (**Supplementary Figure 2**).

#### Repetitive Behavior

On the marble burying test, which measures repetitive and perseverative behavior in rodents^91^, L-DE-71 (but not H-DE-71) adult F1 buried a significantly greater number of marbles relative to VEH/CON (**Fig. 2i**, *P*<.05). A subgroup of F1 was tested at PND 30, but no group differences were measured, possibly indicating age-related physical hypoactivity, reduced habituation to test arena or a latently-emerging impact of PBDEs (**Supplementary Figure 3**). In contrast, no group differences were seen in F0 (**Fig. 2i**). Mean values for nestlet shredding were not affected by DE-71 exposure in F1. However, in F0, the L-DE-71 group showed a mean reduction in nestlet shredding relative to VEH/CON (**Fig. 2j**, *P*<.05). Less nestlet shredding did not translate into poorer maternal nest scores, however (**Supplementary Figure 1**).

### Exposure to L-DE-71 but not H-DE-71 reduces long-term social recognition memory in F1

We determined that SNP scores requiring a 30 min memory retention were abnormal in exposed F1 but not F0. To test the hypothesis that DE-71 compromises consolidation of *long-term* social recognition memory, we subjected F1 to a social recognition memory test (SMRT)^66^. On this test mice with intact memory exhibit less time investigating a familiar juvenile conspecific 24 h after a first exposure. **Figure 3a** shows that VEH/CON and H-DE-71 mice were able to form a social recognition memory of the stimulus by Day 2 since they spent significantly less time with a familiar stimulus mouse (*P*<.05 and *P*<.0001, respectively). We used a one-sample t-test to determine if the sample mean recognition index (RI) was statistically different from previously reported mean RI of 0.65 (Kogan et al, 2000; Tanimizu et al, 2017). Mean RI values for F1 were 0.71 for VEH/CON and significantly *lesser* for H-DE-71 (0.56, *P*<.05), suggesting enhanced recognition memory (**Fig. 3b**). In contrast, L-DE-71 F1 showed an apparently *greater* RI (mean RI, .85, *P*=.07), suggesting they had deficient long-term social recognition memory (**Fig. 3 a,b**). Next, we examined investigation time with a second novel stimulus mouse on Day 2 to determine whether the reduction of investigation time on Day 2 is specific to social memory formation and not due to disengagement. In **Figure 3c**, no significant reduction of investigation time was noted for VEH/CON and H-DE-71, suggesting that the reduction is specific to recognition memory formation (familiar mouse on Day 1). In contrast, L-DE-71 F1 exhibited a significant reduction in investigation time (*P*<.05), further supporting the results above. In this context, we used a one-sample t-test to determine if the sample mean RI is statistically different from a previously reported mean RI of 1. L-DE-71 showed a significantly lower RI (.89, P<.05). During test optimization using untreated controls the 3 but not 1 min of social exposure on Day 1 was sufficient to form a memory on Day 2 (*P*<.01) as reported^66^ (**Supplementary Figure 4**). In summary, these results indicate that developmental exposure to DE-71 at 0.1 mg/kg/d significantly reduces long-term social recognition memory in F1.

**Fig. 3.**
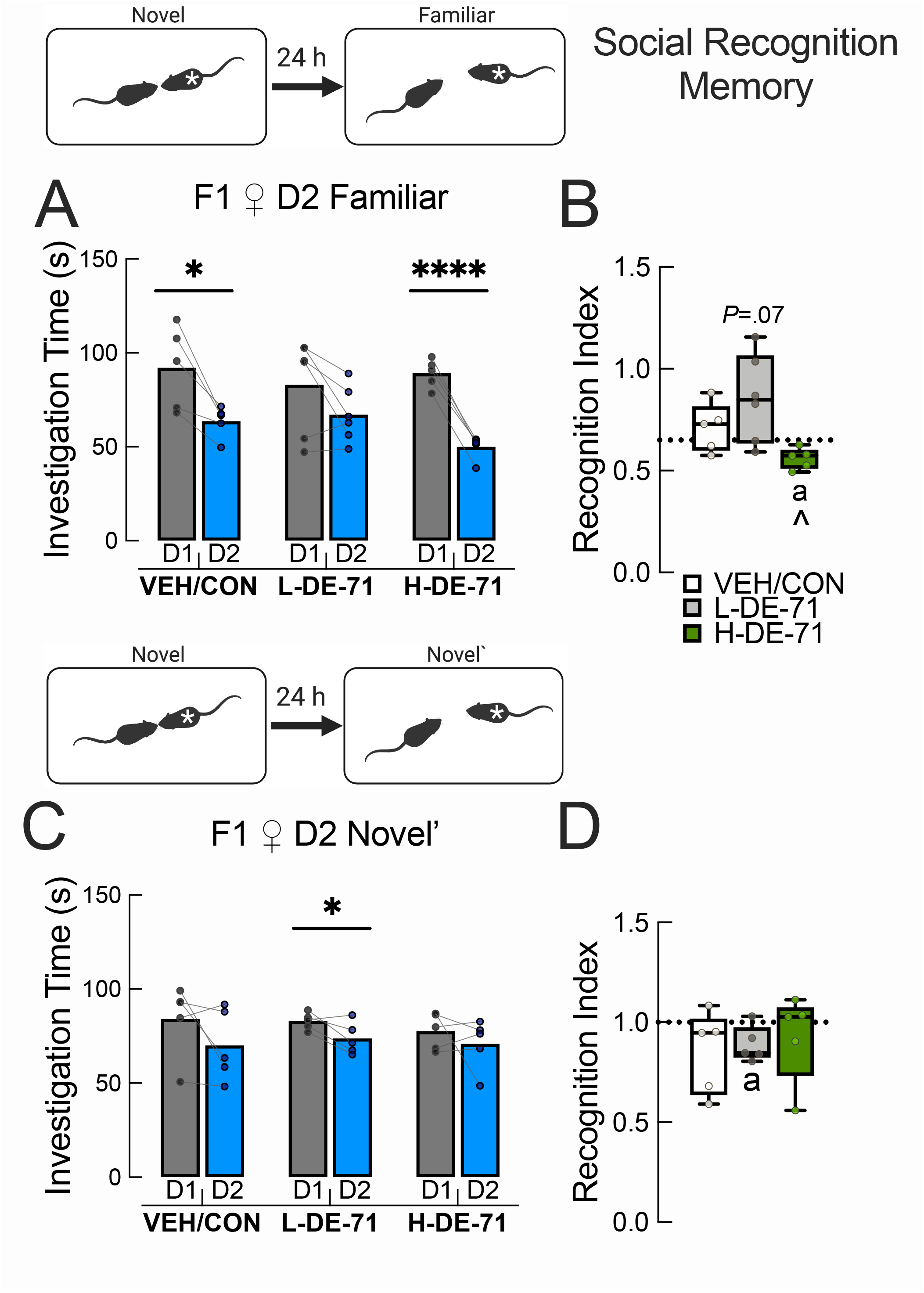
Exposure to L-DE-71 but not H-DE-71 reduces long-term social recognition memory in F1. **a** When using a familiar stimulus the VEH/CON and H-DE-71 F1 mice displayed a significant reduction in investigation time on Day 2 relative to Day 1, indicating normal SRM. In contrast, L-DE-71 showed deficient scores after the 24 h retention period. **b** Corresponding Recognition Index (RI), indicates a strong social recognition memory in VEH/CON and H-DE-71 groups. The mean RI value for L-DE-71 was significantly less. **c** When using a different novel mouse on Day 2 (Novel’) as a control, VEH/CON and H-DE-71 F1 mice did not show significantly reduced investigation time on Day 2 relative to Day 1, indicating reduction in investigation time is specific to familiar juveniles. **d** RI is near 1, indicating no significant change in investigation of Novel’ vs Novel stimulus. \**P*<.05, \*\*\*\**P*<.0001 compared to Day 1 (a, c), compared to VEH/CON (b, d). ^*P*<.05 compared to L-DE-71. ^a^*P*<.05 compared to .65, (a) and 1.0, (b). *n*=5-6 litters/group. ‘*’, stimulus mouse in insets. D, day

### Exposure to L-DE-71 compromises short-term novel object recognition memory in adult F1 and F0

Having found that DE-71 exposure produces significant impairment in the SNP and SMRT, we tested the hypothesis that DE-71 exposure also interferes with non-social recognition memory. Using a novel object recognition memory test (NORT), **Figure 4a** shows that L-DE-71 F1 did not display preferential exploration of the novel object during the Day 1 testing, as did the VEH/CON and H-DE-71 (*P*<.01) indicating that F1 exposed to 0.1 mg/kg DE-71 did not discriminate between objects presented 30 min earlier in the familiarization phase. This was corroborated using a discrimination index which showed that values for VEH/CON and H-DE-71 were >0, indicating memory for previously encountered objects (**Fig. 4b**, *P*<.05). In contrast, L-DE-71 group displayed a negative mean discrimination index (greater preference for familiar object, *P*<.01), which was also significantly reduced compared to VEH/CON (*P*<.001). Representative dwell time maps in the open field arena showed preference for novel object (right corner) for VEH/CON and H-DE-71 on Day 1 (**Fig. 4c**). In contrast, L-DE-71 showed less exploration of novel relative to familiar object. On Day 2 all exposure groups preferred novel over familiar object and showed similar discrimination index mean values and dwell times after a 24 hr retention time (**Fig. 4g, h, i, j**). There were no effects of exposure on locomotion (**Fig. 4e, f, k, l**) as indicated by raster plots (**Fig. 4d, j**) Both L-DE-71 and H-DE-71 exposed dams showed similar short-term memory deficits as L-DE-71 exposed F1 offspring (**Supplementary Figure 5**).

**Fig. 4.**
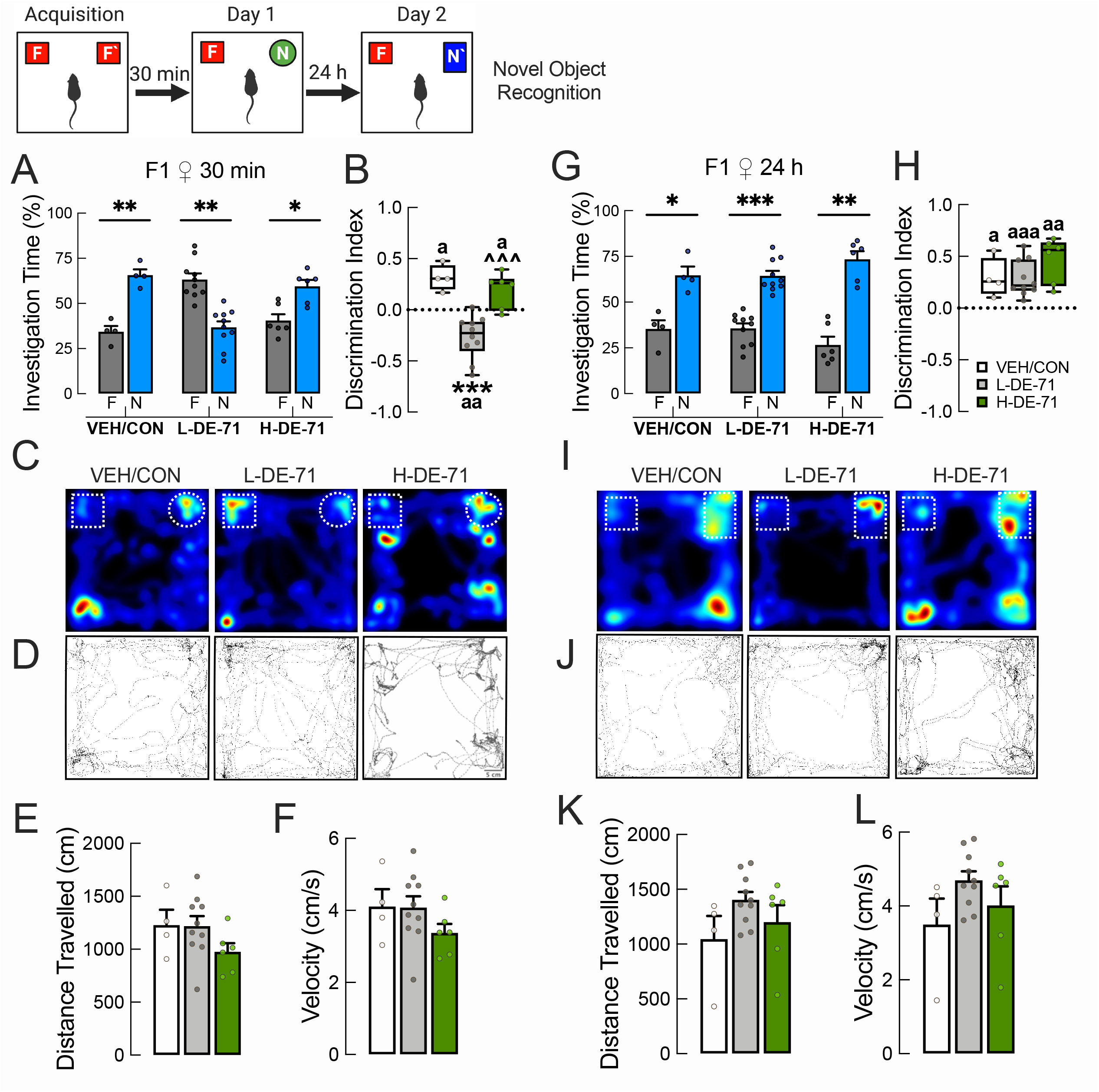
Perinatal exposure to L-DE-71 compromises short-term novel object recognition memory in F1**. a** Investigation time on novel object recognition test. F1 offspring in the VEH/CON and H-DE-71 but not L-DE-71 group show significantly greater time spent investigating the novel (circle, N) vs familiar (square, F). **b** L-DE-71 F1 shows a significant negative discrimination index indicating less time spent with novel object. **c** Representative dwell-time maps (double gradient, blue—minus; red—plus) of time spent exploring novel and familiar objects showed differences in dwell times for different exposure groups. **d-f** Representative raster plots indicate no significant effect of exposure on general locomotor activity quantified as cumulative distance travelled and velocity. **g-l** After a 24 h retention time there was no effect of exposure on investigation time of familiar and novel, discrimination index, dwell-time maps, raster plots, distance travelled, or velocity. \**P*<.05, \*\**P*<.01, \*\*\**P*<.001 compared to familiar object (a) or VEH/CON (b). ^^^*P*<.001 compared to L-DE-71 (b). ^a^*P*<.05, ^aa^*P*<.01, ^aaa^*P*<.001 compared to 0. *n*=4-10 subjects/group. F, familiar object; N and N’, novel object.

### Abnormal social behavior in F1 produced by DE-71 exposure is not due to deficits in general olfactory processing

In order to examine if DE-71-induced deficits observed in social recognition ability were due to insufficient olfactory ability, we subjected female offspring to an olfactory preference test. **Figure 5a** shows that all mice including those treated with DE-71 displayed increased odor sniffing duration for peanut butter over water (*P*<.05-.0001), butyric acid (*P*<.05-.0001), and vanilla (*P*<.05-.001). Similar results were obtained for dams (**Fig. 5b**). These results indicate that, like VEH/CON, DE-71 exposed offspring were able to process sensory signals from different non-social odors with enough sensitivity to show preference for peanut butter over others..

**Fig. 5.**
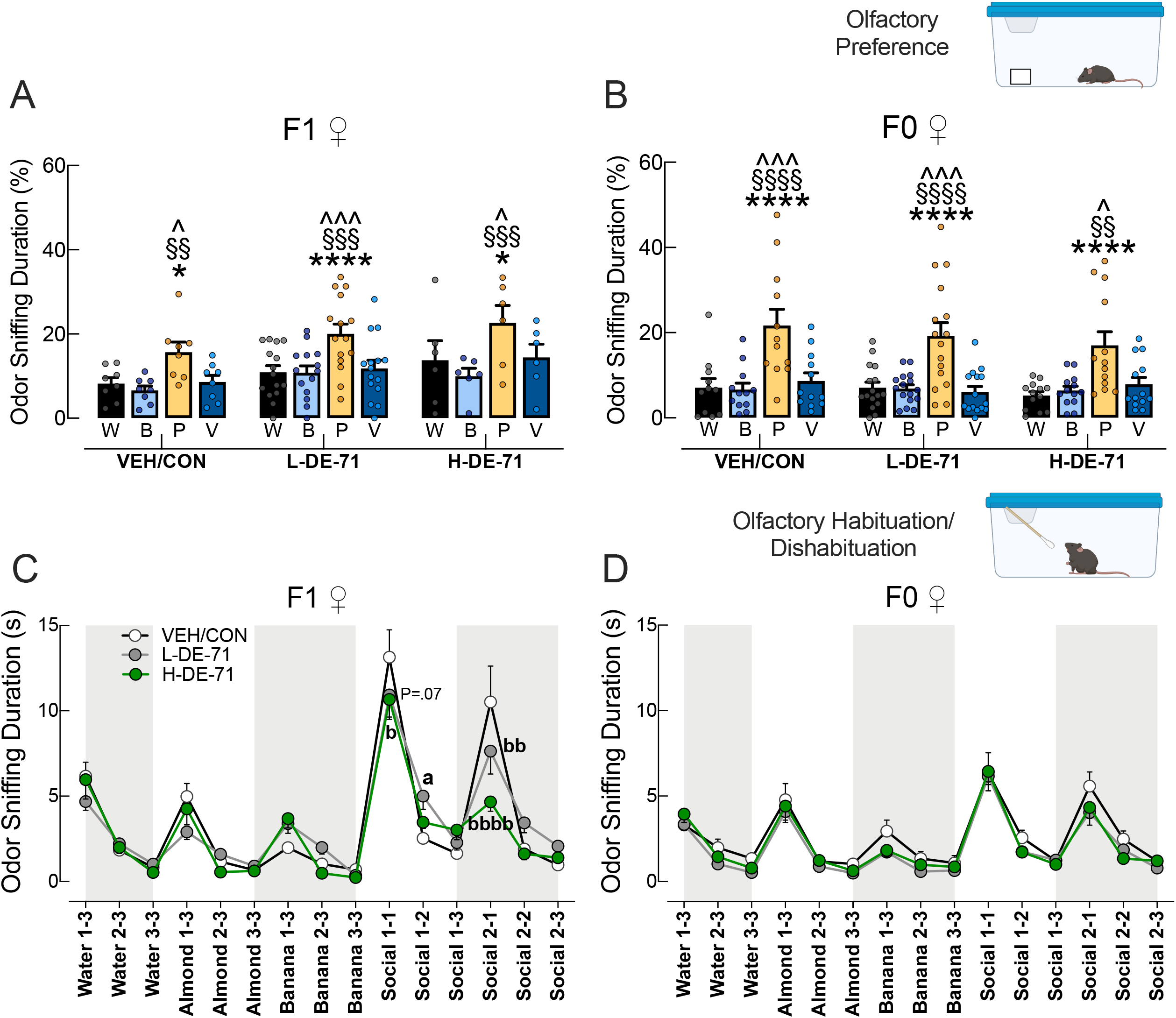
Perinatal exposure to DE-71 does not alter general olfaction function but disrupts discrimination of social odors. **a, b** Olfactory preference test on F1 and F0. Both groups showed normal olfactory preference for peanut butter odor. **c, d** Sniffing time on Olfactory habituation/dishabituation test showed that relative to VEH/CON, L-DE-71 F1 mice showed less habituation to social odor 1 and 2. Both L-DE-71 and H-DE-71 showed abnormally reduced dishabituation to social odor 2. H-DE-71 showed reduced dishabituation to social odor 1, an effect that was apparent in L-DE-71. No group differences were noted for F0. \**P*<.05, \*\*\*\**P*<.0001 compared to water. ^*P*<.05, ^^^*P*<.001 compared to vanilla; §§*P*<.01, §§§*P*<.001, §§§§P<.0001 compared to butyric acid. **^a^***P*<.05, **^aaa^***P*<.001 compared to VEH/CON during habituation. **^b^***P*<.05, **^bb^***P*<.01, **^bbbb^***P*<.0001 compared to VEH/CON during dishabituation. Additional statistical results are summarized in Table 2. *n*=6-15 litters/group (a), *n*=11-16 subjects/group (b), *n*=12-16 subjects/group (c), *n*=12-16 subjects/group (d). W, water, B, butyric acid, P, peanut butter, V, vanilla

**Table 2.** Statistical Results for the Olfactory Habituation/Dishabituation test.

| Group | Exposure | Habituation to water | Dishabituation water to non-social odor 1 | Habituation to non-social odor 1 | Dishabituation non-social odor 1 to non-social odor 2 | Habituation to non-social odor 2 | Dishabituation non-social odor 2 to social odor 1 | Habituation to social odor 1 (Trial 1 to 2) | Habituation to social odor 1 (Trial 1 to 3) | Dishabituation social odor 1 to social odor 2 | Habituation to social odor 2 (Trial 1 to 2) | Habituation to social odor 2 (Trial 1 to 3) |
| --- | --- | --- | --- | --- | --- | --- | --- | --- | --- | --- | --- | --- |
| F1 | VEH/CON<br>N=14-18 | P<.0001 | P<.01 | P<.001 | NS | NS | P<.0001 |  | P<.0001 | P<.0001 | P<.0001 | P<.0001 |
|  | L-DE-71<br>N=16-20 | P<.01 | NS | NS | NS | NS | P<.0001 |  | P<.0001 | P<.0001 | P<.001 | P<.0001 |
|  | H-DE-71<br>N=13-15 | P<.0001 | P<.05 | P<.05 | NS | P<.05 | P<.0001 |  | P<.0001 | NS | NS | NS (P=.09) |
|  | Exposure Group Difference | NS | NS | NS | NS | NS | - VEH/CON vs L-DE-71, P=.07<br>- VEH/CON vs H-DE-71, P<.05 (b) | -VEH/CON vs L-DE-71, P<.05 (a) | NS | - VEH/CON vs L-DE-71, P<.01 (bb)<br>- VEH/CON vs H-DE-71, P<.0001 (bbbb)<br>- L-DE-71 vs H-DE-71, <.01 | NS | NS |
| F0 | VEH/CON<br>N=16-18 | P<.0001 | P<.0001 | P<.0001 | NS | NS | P<.0001 |  | P<.0001 | P<.0001 | P<.0001 | P<.0001 |
|  | L-DE-71<br>N=18-22 | P<.0001 | P<.0001 | P<.0001 | NS | NS | P<.0001 |  | P<.0001 | P<.0001 | P<.01 | P<.0001 |
|  | H-DE-71<br>N=14-18 | P<.01 | P<.0001 | P<.0001 | P<.05 | P<.01 | P<.0001 |  | P<.0001 | P<.0001 | P<.0001 | P<.0001 |
|  | Exposure Group Difference | NS | NS | NS | NS | NS | NS |  | NS | NS | NS | NS |

### DE-71 exposure alters olfactory discrimination of social odors in F1

We used an olfactory habituation/dishabituation test to measure olfactory discrimination. Table 2 indicates the results of the habituation/dishabituation test. The F1 VEH/CON group displayed olfactory habituation to all non-social odors and social odors except non-social odor 2 (banana) as indicated by the decline in time spent sniffing by trial 3 **(Fig. 5c**). F1 VEH/CON displayed olfactory dishabituation when transitioning to a new odor except from non-social 2 (banana) to social 1 (*P*<.01, *P*<.0001). Both DE-71 groups display deficient habituation and/or dishabituation for more than 1 odor (**Table 2**). In particular, L-DE-71 showed reduced habituation to social odor 1 (from trial 1 to 2; *P*<.05; **Fig. 5c**). It appears that banana odor was problematic for most groups. **Figure 5c** shows that compared to VEH/CON, L-DE-71 and H-DE-71 showed less dishabituation from social odor 1 to 2 (*P*<.01, *P*<.0001), suggesting that DE-71 produces reduced olfactory discrimination (hyposmia) especially of social odors, which requires processing via MOE and VNO^92^. In addition, H-DE-71 also showed reduced dishabituation from non-social odor 2 to social odor 1 (*P*<.05). An apparently significant effect was also seen for L-DE-71 (*P*=.07). In combination with normal results on the olfactory preference test, these findings indicate altered social odor discrimination after perinatal exposure to DE-71 potentially associated with altered signaling through the VNO.

Olfactory discrimination of odors in F0 showed normal habituation/dishabituation profiles compared to VEH/CON (**Fig. 5d**, Table 2). There were no exposure group differences found for F0.

Mice were evaluated for anxiety using the EPM test and time spent in closed arms relative to open arms was significantly greater in all exposure groups (*P*<.0001). Similar results were obtained in F1 and F0. There was no effect of exposure on number of total arm entries for F1. In contrast, the H-DE-71 F0 group exhibited significantly fewer total entries relative to VEH/CON (**Supplementary Figure 6**). Using a forced swim test, depressive-like behavior was measured as time spent immobile and there was no significant effect of exposure on time spent immobile for F1 nor F0 (**Supplementary Figure 6**).

### Selective effects of DE-71 on Suok test

Using Suok, we measured the effects of DE-71 exposure on locomotion, exploratory behavior, sensorimotor coordination and anxiety. Relative to VEH/CON, H-DE-71 (but not L-DE-71) F1 showed decreased horizontal activity, as represented by segments crossed (**Fig. 6a**), decreased locomotion (**Fig. 6b**), decreased exploratory activity (**Fig. 6g**), increased SAP (**Fig. 6i**) and decreased grooming (**Fig. 6k**). Falls were significantly decreased in H-DE71, but not when normalized to segments crossed (**Fig. 6d**). In contrast to F1, F0 exposed to L-DE-71 showed decreased hind leg slips (**Fig. 6e**) and increased SAP relative to VEH/CON (**Fig. 6i**). There were no significant differences on the other measures.

**Fig. 6.**
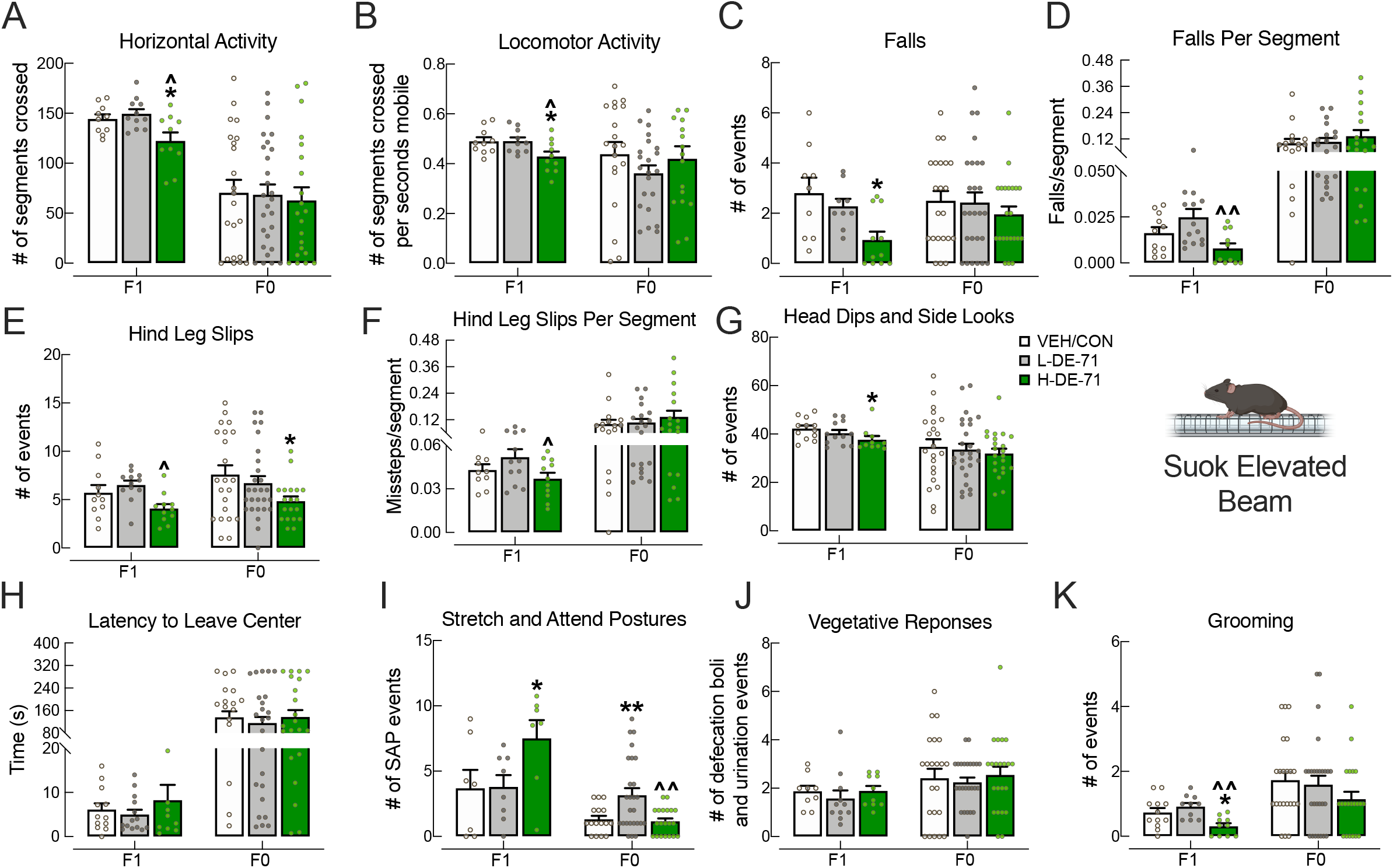
Selective effects of DE-71 exposure on Suok Test. Female offspring and dams were tested on SUOK for: **a, b** locomotion; **c-f** sensorimotor coordination; **g** exploratory activity; **h-j** anxiety behaviors; and **k** autogrooming. Only H-DE-71 F1 showed decreased mean values in a, b, g, k and increased i whereas F0 exposed to L-DE-71 showed increased mean value in i. \**P*<.05, \*\**P*<.01 compared to corresponding VEH/CON. ^*P*<.05, ^^ *P*<.01 compared to corresponding L-DE-71. *n* for F1 (litters/group): (a) 10-11; (b) 10; (c) 9-11; (d) 10-11; (e) 11-12; (f) 10-12; (g) 10-12; (h) 10-14; (i) 7-8; (j) 9-11; (k) 10-12. *n* for F0 (subjects/group): (a) 21-27 ; (b) 17-22; (c) 22-26; (d) 16-20; (e) 19-27; (f) 16-20; (g) 22-27; (h) 22-27; (i) 16-26; (j) 22-25; (k) 21-27.

### Early-life PBDE exposure does not alter locomotion on the open field test

The open field test informs about locomotion, habituation to novelty and anxiety. All F1 exposure groups showed similar reduced exploratory activity over time (habituation), measured as reduced distance traveled and velocity over the 1 h test (**Fig. 7 a,b,** *P*<.0001). Between-group comparisons showed no effect of exposure for F1. These results helped us rule out concerns of hyper- or hypo-mobility in DE-71 exposed female offspring relative to VEH/CON as reported after acute exposure to 0.8 mg/kg BDE-99 at PND 10^29, 93^. Other studies using chronic exposure of mouse dams to low doses of BDE-47 (0.1 mg/kg) or -99 (0.6 mg/kg) from gestation through 3rd week of lactation have shown inconsistent results with both hypoactivity and no effect reported on the open field test in female offspring^94,95,96^.

**Fig. 7.**
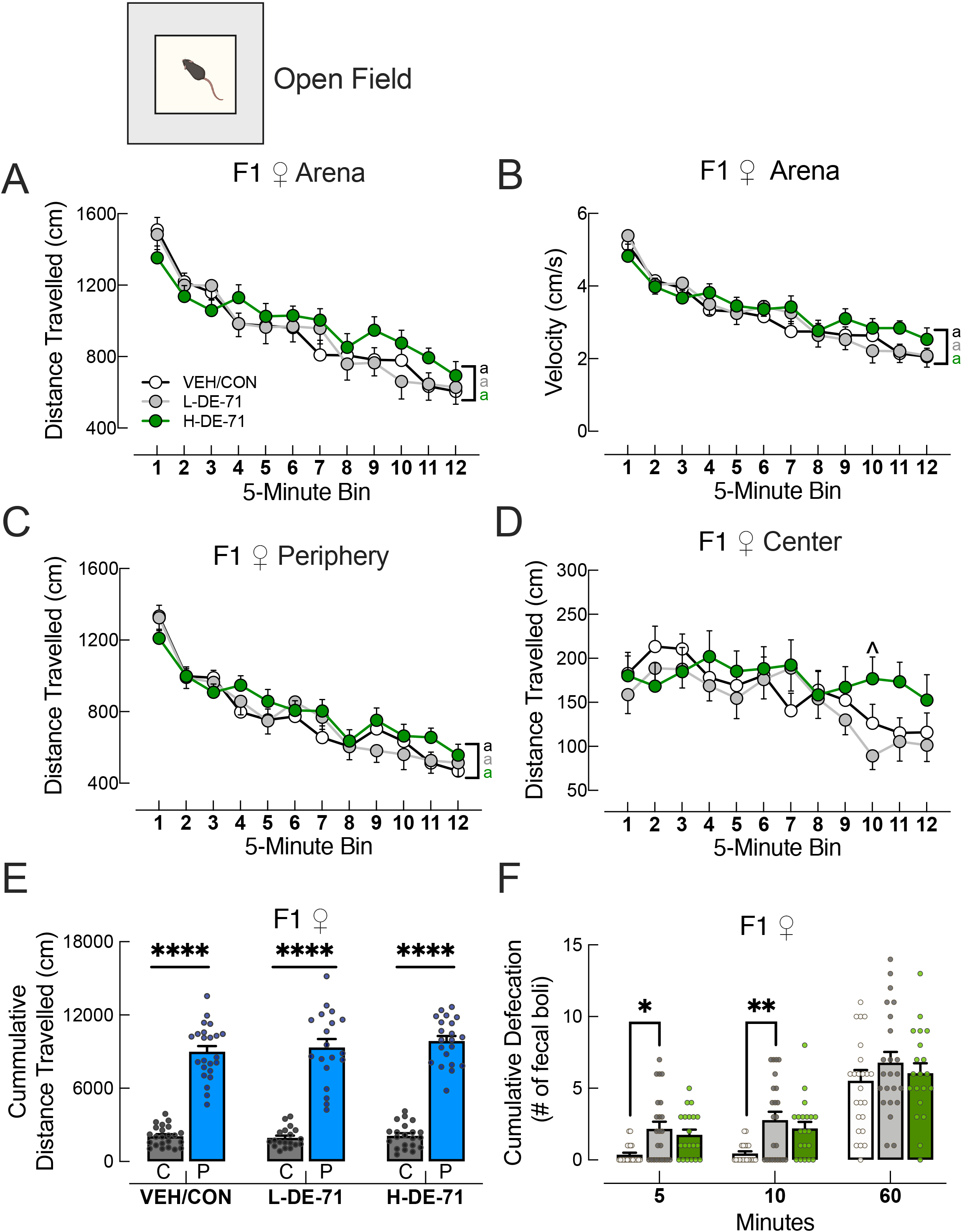
Early-life PBDE exposure does not alter locomotion or anxiety on the open field test. **a, b** Distance traveled in open field arena. All F1 exposure groups showed similar reduced exploratory activity and velocity over the 1 h. **c, d** Exploration time in periphery and center for all groups showed habituation only in the periphery. **e** Exploration time in center was significantly less than in periphery for all groups, suggesting no exposure effects on anxiety. **f** Another measure of anxiety, number of fecal boli, indicated increased emotional reactivity in the L-DE-71 F1 relative to VEH/CON. \**P*<.05, \*\**P*<.01. \*\*\*\**P*<.0001 compared to center (e) or VEH/CON (f). ^*P*<.05 compared to corresponding L-DE-71. ^a^P<.0001 compared to initial time bin for corresponding treatment group. *n*=19-23 subjects/group. C, center zone; P, periphery zone

Exploration time in center and periphery zones for all exposure groups (**Fig. 7c,d**) showed habituation only in the periphery (*P*<.0001). **Figure 7e** shows that total distance travelled in periphery was similarly and significantly greater in all exposure groups (*P*<.0001). Another measure of anxiety, number of fecal boli at 5 and 10 min into the test, indicated increased emotional reactivity in the L-DE-71 F1 relative to VEH/CON, respectively (**Fig**. **7f**, *P*<.05, *P*<.01). For F0 H-DE-71 induced greater distance travelled and velocity in the arena and exploration in the periphery zone as compared to VEH/CON (*P*<.05-.01) and L-DE-71 (*P*<.05-.0001). L-DE-71 mice produced more fecal boli at 60 min relative to VEH/CON (**Supplementary Figure 7**).

### DE-71 alters prosocial gene expression in brain regions involved in social behavior

To correlate the behavioral findings with changes in gene expression of the social neuropeptides that are key mediators of complex social behavior such as vasopressin (*Avp*), oxytocin (*Oxt*), PACAP (*Adcyap1)* and their receptors, we measured the relative expression of these genes from micropunches of discrete brain nuclei involved in social behavior: lateral septum, amygdala, bed nucleus of the stria terminalis (BNST), SON and PVN. **Figure 8** shows that *Avp* was decreased in BNST of L-DE-71 (*P*<.05) and SON of H-DE-71 (*P*<.05). Similarly, *Oxt* mRNA transcripts were decreased in the BNST of L-DE-71 and H-DE-71 (*P*<.05) and SON of L-DE-71 (*P*<.05). *Oxtr* levels were increased in PVN of L-DE-71 (*P*<.05) and the BNST and amygdala of H-DE-71 (*P*<.05). For *Avp1ar,* BNST levels were upregulated in L-DE-71 (*P*<.05) and downregulated in SON in H-DE-71 (*P*<.05). No changes in *Adcyap1* or *Adcyap1r1* were observed.

**Fig. 8.**
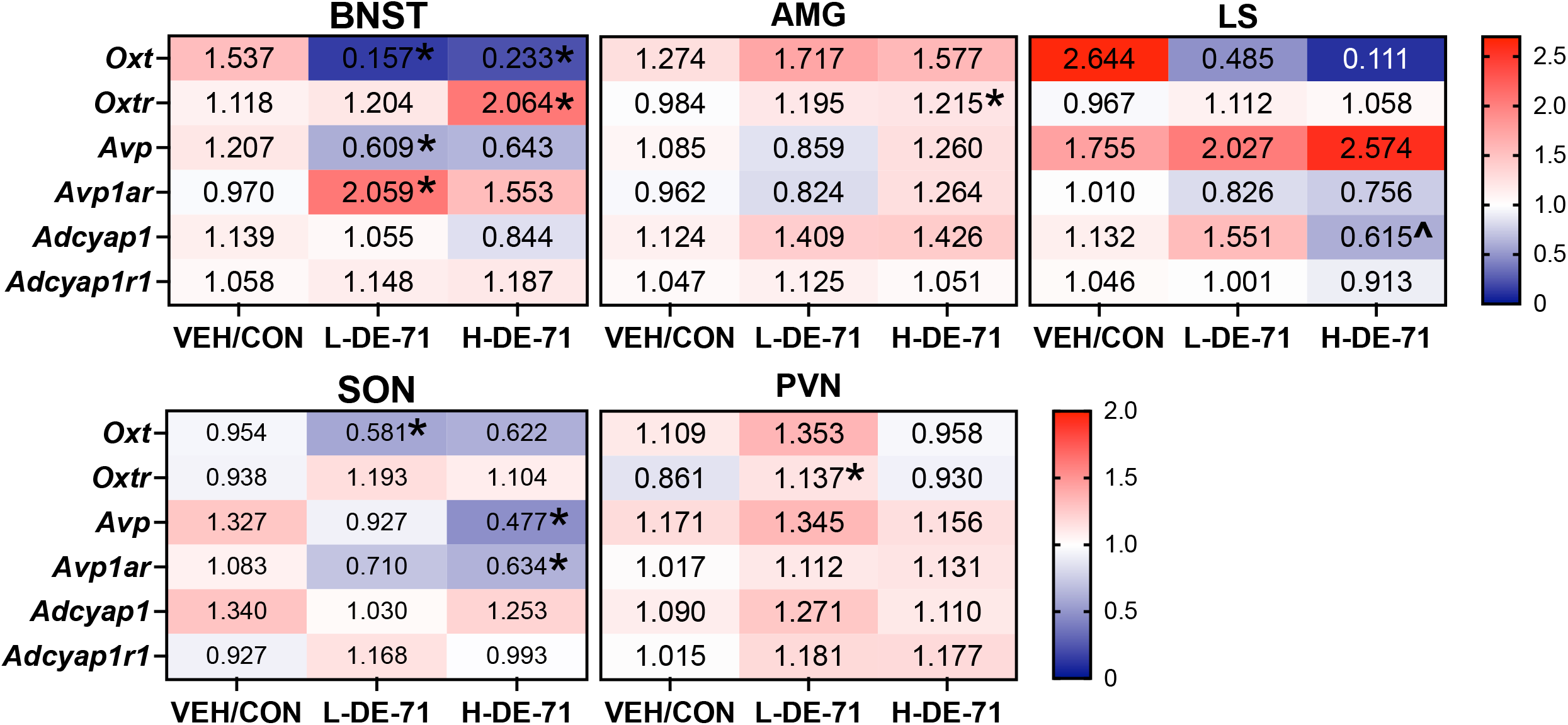
DE-71 exposure alters gene expression in select brain regions involved in social behavior in F1 females. Heatmap representation (double gradient, blue—minus; red—plus) of RT-qPCR analysis with the respective fold-change value (mean) of each gene studied by brain region. *n*= 4-17/group. \**P*<.05 compared to VEH/CON. ^*P*<.05 compared to L-DE-71. BNST, bed nucleus of the stria terminalis; AMG, amygdala; LS, lateral septum; SON, supraoptic nucleus; PVN, paraventricular nucleus.

### DE-71 alters plasma vasopressin but not oxytocin levels in F1 offspring

We next measured plasma oxytocin and vasopressin concentrations and their association with social behavior phenotypes. Figure 9 shows that plasma AVP levels in L-DE-71 F1 females were significantly elevated relative to VEH/CON (*P*<.05). In contrast, there were no group differences in plasma OXT levels.

**Fig. 9.**
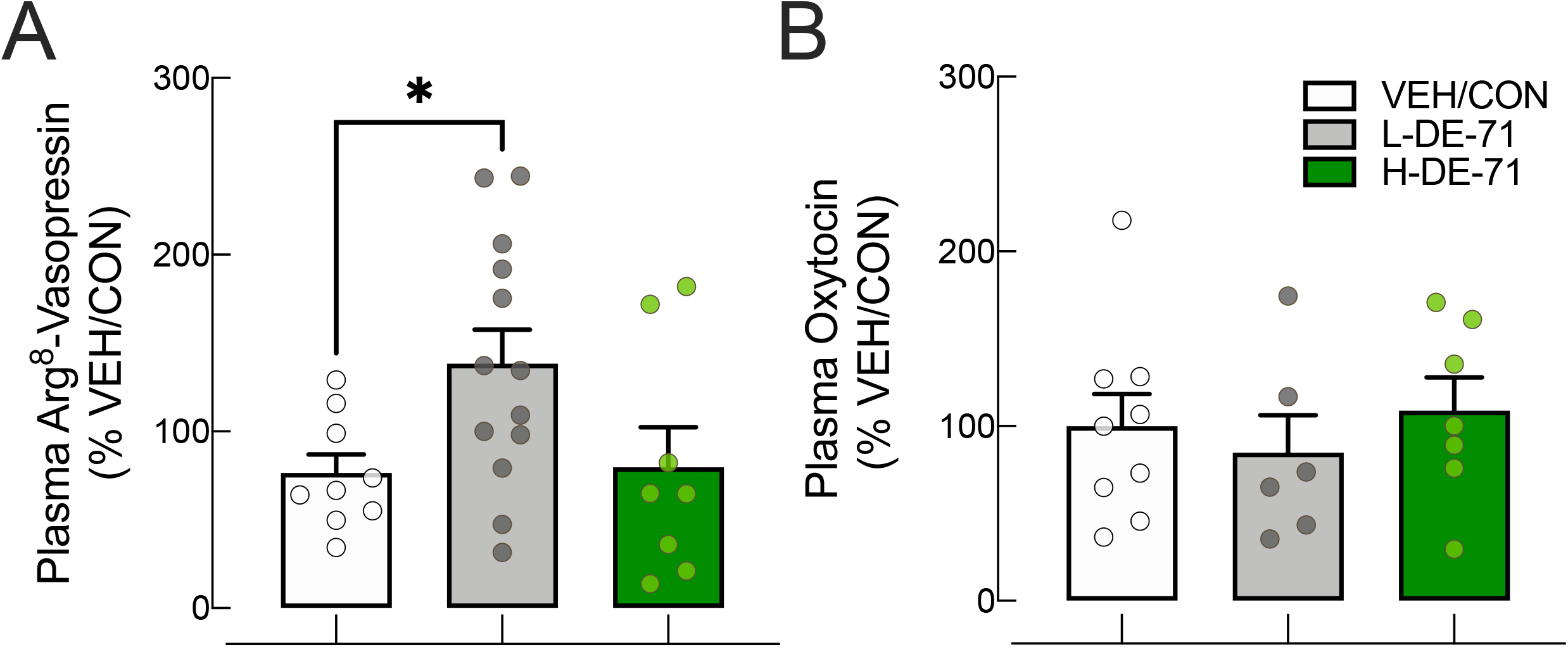
Perinatal exposure to DE-71 exaggerates plasma ARG^8^-vasopressin but not oxytocin (OXT) levels in adult F1 female offspring. **a** Plasma Arg-8 vasopressin measured using EIA using blood taken at sacrifice. L-DE-71 exposed offspring showed elevated levels. **b** OXT levels showed no changes. \**P*<.05 compared to VEH/CON. *n*=8-13 subjects/group (a); *n*=6-8 subjects/group (b).

## Discussion

Growing evidence suggests a positive association between early-life exposure to PBDEs and neurodevelopmental alterations^97^. Environmental factors, including xenobiotic chemical exposures, may provide a plausible explanation for the rising incidence of NDDs with social deficits^4^, however, experimental evidence has not established a direct link with specific candidate chemicals. With this purpose, our study is the first to investigate the effects of the penta-PBDE mixture DE-71 on behaviors and neurochemical/endocrine profiles relevant to several core ASD symptom domains. Our experimental design exposes progeny to the full complement of congeners found in human breast milk^98^. The major findings reveal that developmental DE-71 exposure produces enduring deficits in social recognition, repetitive behavior and social odor discrimination in female offspring. The behavioral phenotypes occurred concomitantly with changes in peripheral AVP and in the neuromolecular phenotype of *Oxt* and/or *Avp* signaling pathways in brain regions that coordinate complex social behaviors. Together, the behavioral, sensory and neurochemical phenotypes produced by DE-71 may provide a novel, comprehensive ASD-relevant model with high translational impact. Our results are congruent with a disrupted developmental trajectory of the Social Processing Domain as outlined in the 2010 NIMH Research Domain Criteria (RDoC) framework^1^; a key characteristic of ASD pathology. Our work is further strengthened by the use of the litter as the unit of statistical analysis, thus overcoming risk of bias (RoB) of individual studies^99^ and inter-individual variability. Further, DE-71 produced the common hormetic response, such that only 0.1 but not 0.4 mg/kg exhibited most of the behavior changes even though there was a dose-dependent increase in brain accumulation of ∑PBDE congeners. Moreover, confirmed the augmented susceptibility to developmental relative to adult exposure, highlighting the significance of chemical exposures during critical neurodevelopmental windows. Collectively, these data support the conclusion that environmental xenobiotics impact social behavior and related neurochemical signaling pathways in mice relevant to NDDs.

### Perinatal DE-71 exposure produces deficient social recognition and increases repetitive behavior in adult female offspring

Our main finding was that *in utero* and lactational transfer of DE-71 produces behavioral phenotypes resembling two core behavioral features of a ASD DSM-V diagnosis: deficits in social reciprocity and communication and repetitive/stereotyped behaviors^100^. With respect to the latter, female offspring exposed to L-DE-71 showed increased activity on a marble burying test indicative of repetitive behaviors in rodent models of ASD^79^. Developmental L-DE-71 exposure also produced deficient short-term social memory (SNP) and long-term social recognition memory (SRM), while sociability (SOC) was not affected, ruling out a lack of the ‘social motivation’ component of social cognition. SRM is considered to be another distinct behavioral domain and important for the ‘knowledge of self and others’ component of social cognition^65^.

Though much is still unknown about the neural correlates of social behavior, the social motivation and social recognition domains have been shown to be independent of each other. For instance, deficits in SNP can occur without decrements in sociability in other models of deficient social behavior induced by high fat diet^101^ or C-section delivery^102^. The former can be restored by OXT administration. In other reports, restoring OXT content in the PVN with probiotic therapy (*L. reuteri)*, in maternal high fat diet and valproic acid offspring, rescues SOC and SNP, but not other ASD endophenotypes^52, 53^. In a maternal immune activation (MIA) model, both SNP and SOC are deficient but unable to be restored by *B. fragilis*, while repetitive behavior is rescued^103^. Taken together, these results suggest that different mechanisms and/or circuitry govern the various social behavior domains that can be selectively isolated by experimental contrast and susceptibility to early-life PBDEs. Specifically, perinatal DE-71 exposure significantly compromises the social recognition domain of social cognition, which is more relevant to ASD since the behaviors related to knowledge of self and others such as facial recognition, empathy and evaluation of emotion of others are disrupted in ASD patients^65, 104^.

Our findings indicate deficient short-term social recognition and long-term social recognition memory in L-DE-71 F1, suggesting our results may be translational to ASD and other NDDs characterized by psycho-social deficiencies. While findings of epidemiological studies evaluating associations between PBDEs and social deficits/ASD are mixed (Gibson et al, 2018; Braun et al, 2014; Vuong et al, 2016), a higher risk of poor social competence has been found with increasing postnatal exposure to BDE-47 (4 yr old child serum) (Gascon et al, 2011). BDE-47 levels in cord blood have also been positively associated with poor social domain development in 24 month-old toddlers^35^. Previous rodent studies examining the effects of environmental pollutants on social behavior have produced inconsistent results perhaps due to heterogeneity of brominated (BFR) flame retardants used, timing of exposure, sex and/or model organism used. Importantly, the only other study examining the effects of DE-71 (0.3 and 1.6 ppm) on social behavior supports our findings. Fernie and colleagues (2005) found fewer and less appropriate pair-bonding and courtship behaviors in exposed captive kestrels^105^. In contrast, female offspring exposed to BDE-47 perinatally via mother showed reduced sociability relative to controls ^46^ but no effect of BDE-47 (at 0.03 mg/kg) was detected on SNP unless administered to genetically altered mice lacking methyl–CPG binding protein 2 (*Mecp2)*, a frontal cortical protein negatively associated with ASD^106^. Male CD-1 mice developmentally exposed to BDE-47 (0.2 mg/kg) display reduced time with conspecfics but show no effect on SNP relative to controls^47^. In rats perinatally exposed to PBDE-47 (50 mg/kg) Li and others (2021) report preference for stranger over familiar conspecific and for social stimulus over empty corral but with reduced time spent in exploration^107^.

Our findings are also supported by perinatal exposure studies using other BFRs such as Firemaster 550 (6.6 mg/kg/day) and its BFR and organophosphate components alone (3.3 mg/kg/d), which produce deficits in social recognition after 24 h retention in a sex and exposure-specific manner in rats^108^. Perinatal exposure to Firemaster 550 also produces abnormal partner preference in female prairie voles (1 mg/kg)^109^. Using low doses of BDE-209 (0.12 ng/mouse/day, s.c.), Chen and colleagues (2019) did not observe deficient sociability nor SNP in exposed male mice offspring^110^. Therefore, it appears that PBDE effects on social behavior may be congener-and dose-specific.

### Perinatal DE-71 exposure produces deficient novel object recognition memory in dams and adult female offspring

A complex interplay between forebrain regions is responsible for normal social recognition^65,66,67^ including hippocampal circuits underlying social memory formation and amygdalar circuits that process social signals such as volatile odorant pheromones that trigger social and reproductive behaviors^111, 112^. DE-71-exposed socially deficient mice also showed abnormal NOR memory suggesting abnormal function in hippocampus since it serves as an integration hub underlying both social recognition memory and recognition memory^64, 114, 115^. Toxicological studies of developmentally administered single BDE congeners or DE-71 have not examined effects on NOR or SMR^99^. However, previous studies using peri/postnatally administered single BDE congeners such as BDE-153 (0.9 mg/kg bw) and -47 (0.03 mg/kg bw) have showed neurotoxic actions on hippocampal-dependent function related to spatial memory^95, 99, 113^. In support of our findings, evidence from human studies suggest that more than one environmental BDE congener may produce risk for cognitive impairments in children. For example, several PBDEs found in maternal samples (BDE-47, 99, 100, 153) are associated with children’s lowered IQ and cognitive scores^36, 114, 115^ mental/physical development^116^ and fine motor skills, attention and cognition^22^.

It is not surprising that L-DE-71 F1 mice showed both deficient NOR memory and SRM. However, while deficient SRM was seen at both short and long-term retention times, NOR memory deficits were evident at short-term retention time only. Moreover, F0 showed deficits only in short-term NOR memory indicating that short-term social recognition ability and short-term novel object recognition memory are distinct constructs. Therefore, PBDEs may target different brain circuits participating in general and social memory processes and/or different neurochemical systems within each circuit. For example, hippocampal OXTRs are necessary for short-term social recognition but not novel object recognition memory in male mice^56^.

### Perinatal exposure to DE-71 alters social odor discrimination in adult female offspring

Recognition of conspecifics in rodents depends on proper identification, discrimination and processing of olfactory cues present in urine and secretions from skin, reproductive tract and scent glands^111, 112^. We found that the disruption of social behavior after perinatal DE-71 exposure is coincident with abnormal profiles of olfactory habituation/dishabituation to social odors. For example, socially-deficient L-DE-71 mice also displayed a reduction in habituation to social odors and in dishabituation from one to another social odor. In H-DE-71 mice the deficits in olfactory discrimination and social behavior (reduced sociability with normal social recognition ability and memory) were relatively less severe. In combination, these results suggest that DE-71 effects on olfactory discrimination are specific to social odors and that olfactory as well as social deficits are different depending on dose. An olfactory preference test of non-social odors showed no deficits in general olfactory processing. Therefore, reduced social and recognition memory produced by DE-71 is concomitant with deficient social odor discrimination. It is unclear why PBDEs are more neurotoxic to social odor processing but may depend on the different CNS pathways taken by signals from neutral and social odors. Chemosensory cues are processed through two olfactory systems; neutral odors (banana and almond) are processed through the main olfactory epithelium (MOE) and social odors through both MOE and the vomeronasal organ (VNO)^92^. Signals are then processed through amygdala and hypothalamus to trigger innate social and reproductive behaviors. There are no previous studies on PBDEs and olfactory function although BDE-47, -85, - 99 can concentrate to the epithelium of the nasal cavity^117^ and developmental exposure to BDE-209 impairs subventricular zone (SVZ) neurogenesis and olfactory granule cell morphology in mice^118^. However, a recent study indicates that prenatal exposure to PCBs (Aroclor 1221, 1 mg/kg) impair mate preference behavior based on olfactory cues concomitant with impaired odor preference for mates with different hormone status in adult female offspring^119^. Our findings that early-life PBDE exposure alters social odor discrimination may translate to autistic humans which are prone to hypo- or hyper-reactivity to sensory stimulation (American Psychiatric Association, 2013 DSM-V); recent studies suggest that this may include atypical olfaction^120^. Indeed, several olfactory outcomes have been reported in children with ASD, i.e., abnormal odor responses, difficulties in emotional reaction to odors, impaired detection thresholds and odor identification as well as heightened olfactory sensitivity^120,121,122^. Further research is needed to discern the mechanisms by which PBDEs may act to alter social odor discrimination and if this contributes to their social recognition deficits. Interestingly, extrahippocampal OXT and AVP systems, that contribute to short-term social recognition, also modulate detection and processing of social _odors_^123, 124^.

### Perinatal DE-71 alters AVP and OTergic neuromolecular phenotypes in brain regions that coordinate complex social behaviors

Our lab has previously shown that in vitro and early-life exposure to PBDEs (and PCBs) produce neuroendocrine disruption of the prosocial neuropeptide, vasopressin, under osmotically stimulated state^59, 60, 63^. Therefore, the observed PBDE-induced deficits in SNP and SMR may result from altered function of AVP and/or OXT neurochemical systems. Here we show that L-DE-71 downregulates *Avp* in BNST, which provides sexually dimorphic AVPergic innervation to LS^125^. Diminished AVPergic signaling to LS may explain reduced social recognition memory in L-DE-71 F1 females, since AVP1a receptor antagonism in LS compromises social discrimination especially well in females^126^. DE-71-mediated upregulation in BNST *Avp1ar* may represent a compensatory effect. Interestingly, *Avp1ar* in the ventromedial nucleus (VMN) is upregulated by the PCB mixture A1221 in female rat (but not male) offspring and is not dependent on estrogenic pathways^127^. The observed downregulation of *Avp* in SON may also impact social recognition ability indirectly via reduced AVPergic-mediated activation of BNST^128^.

At 0.1 mg/kg, DE-71 also produced elevated plasma AVP which is consistent with less inhibitory regulation over axonal secretion of AVP hormone resulting from reduced levels of central AVP^129^. DE-71 can interfere with intracellular calcium dynamics and increase exocytosis in PC12 pheochromocytoma endocrine cells^130^ and potentially increase secretion of stored AVP depots in axonal terminals located in the posterior pituitary releasing AVP into the bloodstream. DE-71 appears to alter the central OXTergic system which is also necessary for social recognition and partner preference^67^. For example, mice with OXT gene deletion fail to remember recently encountered individuals and do not show the typical decline in preference during subsequent exposures to the familiar mouse, an effect which can be rescued by central administration of OXT^131^. A recent report has demonstrated that OXT receptor blockade, in the extrahypothalamic population of oxytocinergic neurons of the BNST, impairs social recognition in female and male rats^132^. Here we show that in the BNST, L-DE-71 female F1 display significantly reduced *Oxt* mRNA transcripts. Assuming that there is a positive correlation between gene and peptide content and release, one interpretation of our data is that L-DE-71 exposure reduces OXTergic signaling which is necessary for normal social discrimination^132^. Our results further indicate that BNST-originating OXT may be sufficiently important for activating BNST OXTR relative to PVN-originating OXT^133^. H-DE-71 F1 females display reduced BNST *Oxt* mRNA in conjunction with upregulated *Oxtr*, a likely compensatory mechanism to maintain OXT receptor signaling at normal levels. Importantly, L-DE-71 also reduced *Oxt* transcripts in the SON. Because local release of OXT from SON dendrites that extend to MeA promotes social recognition through amygdalar OXTR^134, 135^, downregulated SON *Oxt* may underlie, in part, the associated SNP and SMR deficits. OXT in the BNST also drives stress-induced social vigilance and avoidance that may be at play in social behavior domains examined here^136^. Since the promoter regions for genes of both oxytocin^137^ and vasopressin systems^138^ are susceptible to epigenetic modification^138^, these genes may be altered by global DNA methylation measured after developmental BDE-47 exposure^46, 139^. Our findings may have translational value since altered OXT and AVP mechanisms in humans have been implicated in ASD^140,141,142,143^.

### Specificity and comprehensive profile of PBDE toxicant model of ASD

A recent meta-review has put forth recommendations to improve ASD model characterization in rodent studies such that information about reciprocal social communication and stereotyped repetitive behavior domains are characterized in the same animals^7^. To this end, we used established protocols to measure ASD-relevant and other comorbid behaviors in order to fully characterize the DE-71-induced phenotypes^50, 144^. We found that the effects of DE-71 were specific to social novelty preference and social recognition memory as well as repetitive behavior and olfactory discrimination of social odors. Alterations were specific to offspring exposed perinatally via maternal transfer of environmentally relevant BDE congeners; adult exposed mothers were mostly unaffected. DE-71 had little to no effects on behaviors representing the domains of anxiety, depression and locomotion indicating ASD-relevant specificity without general neurological effects. In addition, there were no indications of reduced general health, i.e., body weight in pups nor gross abnormalities in maternal nest conditions. We have recently reported that similarly exposed (L-DE-71) female offspring, and to a lesser degree, their exposed mothers, display diabetic symptomatology, effects which may relate to the present findings^71^. Importantly, we used multiple behavioral tests to validate social and other constructs studied (locomotion and anxiety). For example, for all F1 groups, the frequency of total entries on EPM and distance travelled on OFT yielded similar results on locomotion. In addition, results on time spent in open arm on EPM, and latency to leave center on Suok was consistent with duration in center of OFT.

Our DE-71 model of ASD also shows altered prosocial peptide neurotransmitters/neurohormones that are critical to ensuring proper development of social brain networks. In particular, the vasopressin and oxytocin systems are critically involved in social cognition with mutations having sociobehavioral impact that have been implicated in core symptoms of autism^145^. These neurochemical systems are being actively studied as potential targets of future therapeutic interventions for ASD^146, 147^. In light of incongruent findings reported by past rodent and human studies^7^, we believe that our findings brings us closer to understanding the risk of ASD posed by xenobiotic endocrine disrupting chemicals. Nevertheless, human and rodent studies reporting on the relationship between PBDE exposure and autistic phenotype are few in number or have yielded inconclusive results and this field would benefit from additional detailed epidemiological and animal studies on the relationship between persistent organic pollutants (POPs) and risk of ASD.

### Maternal transfer of BDE congeners in DE-71 and their brain accumulation in female offspring is dose- and time-dependent

BDE congener composition found in PND15 exposed brains mimics that found in humans. BDE-28, -47, -99, -100, -153 were common congeners found at ppb in both DE-71-exposed offspring groups at PND15 with three-fold greater levels in H-DE-71 than L-DE-71. ∑PBDE values for adult serum are 30–100ng/g lipid^29^ and 3 to 9-fold higher in infants because of exposure through breastmilk and in toddlers because of exposure through house dust and the diet^25, 26, 148, 149^. Serum ∑PBDE values can reach 482 ng/g l.w. in toddlers (California 18 month-old) (Fischer et al., 2006) but lesser values have also been reported, i.e., 127 ng/g l.w.^150^ in Ohio 2 year-olds and 100 ng/g l.w. for North Carolina 12-36 month-old toddlers^151^. Using a divisor factor of .095 to convert w.w. to l.w. (unpublished observations), we estimate our mean ∑PBDE in L-DE-71 F1 at PND 15 to be 1.7- to 8.2-fold greater, suggesting ours represents a translational model of maternal PBDE transfer. The main congeners in PND15 brains, BDE-47, -85, -99, -100, -153, and -154, accounting for 97% of the mean ∑PBDEs, also comprise the majority of congeners (96%) in DE-71^32^. These and other congeners found in offspring brain samples, i.e., BDE-17, 28, 49, 138, 139, 140, 183 and 184, have also been detected in human serum and/or breastmilk^152^. Importantly, to our knowledge, BDE-49, -140, -183, -184 have not been previously reported in DE-71 exposed rodent brain^32^.

Although using a mixture like DE-71 closely models the PBDE contamination previously shown in human breastmilk, there are some congeners, found at low levels in breastmilk, that we did not detect in offspring brain, i.e., BDE-7, 15, 71, 77, 119, 126^152, 153^. Of these BDE-71 and - 126 are present in DE-71^154^ Little or no information is available about the penetrance and/or neuroactivity of the missing congeners. In most rodent studies, which have focused on a single PBDE congener, BDE-47, dominantly detected in humans, have not reported pervasive effects on social behavior as we do here using DE-71. We speculate that BDE-47 alone is not effective in producing deficits in social recognition and memory and that, instead, several PBDE congeners may act synergistically and/or additively to generate these abnormal phenotypes, reinforcing the need for *in vivo* studies using PBDE formulations that mimic child exposure. By PND 110 the BDE composition in F1 brain was limited to BDE-153, which may be partly responsible for neurotoxicity seen. BDE-153 has been positively associated with lower IQ in children and can cause impaired learning and memory in animal studies^113, 115^. However, while BDE-153 (and an additional 6 congeners) is detected at ppb in *postmortem* brain samples from 4-71 year-old born 1940 to 2000, it is significantly depleted in autistics relative to normal subjects^88^. The relatively lower retention of BDE-47 is in line with a previous report of differential tissue accumulation and disposition of BDE congeners attributed to their toxicokinetic properties^155^. *Cyp*-mediated biotransformation of BDE-47 and -99 (but not 153) may contribute since these congeners contain sites with adjacent unsubstituted carbons where the metabolism occurs^156^. By PND 110, most of the congeners were eliminated from brain except for BDE-153; minimal metabolism of this congener is observed in rodents due to its high lipophilicity as determined by a high octanol-water partition coefficient (Log K_ow_)^156^. Our findings suggest that elevated brain levels of key BDE congeners during early postnatal development may predispose children to neurobehavioral alterations related to ASD.

## Conclusion

Though the role of environmental toxicants in the etiology of NDDs is poorly understood, our data support a link between maternal toxicant exposures and abnormal social and repetitive behavior in offspring that is relevant to ASD. We have shown that early-life exposure to DE-71 leading to these phenotypes is associated with human-relevant levels and composition of BDE congeners penetrating the postnatal offspring brain via maternal transfer. DE-71 actions almost exclusively affect F1 progeny, supporting previous studies showing the particular susceptibility of developing nervous system to neurotoxic actions of PBDEs. These abnormal social behavior phenotypes are specific to social novelty preference and social recognition memory and are also associated with excessive repetitive behavior, as well as neurochemical and social odor processing correlates – suggesting that discrete brain systems are targeted by PBDEs to promote neurodevelopmental abnormalities.. Future studies are needed to discern if DE-71 actions are sexually dimorphic and extend to exposed male offspring. We believe that our environmental toxicant mouse model has utility in future studies examining the relationship between environmental xenobiotics, neurodevelopmental reprogramming and the rising incidence of NDDs.

### Limitations of the Study

The results of the PCR analysis provide novel results on the effects of PBDE exposure on the expression of gene markers for small ‘prosocial’ neuropeptides and their receptors in specific regions of the social brain network. However, these restricted regions vary in cell density and limit the RNA yield for genes of interest (GOIs), especially in the amygdala and LS. Moreover, relative expression was more variable for ROIs that have low expression of GOIs, i.e., *Oxt* for LS. To improve our experimental data, we followed MIQE guidelines to optimize oligonucleotide primer efficiency and target specificity. Since the methodological approach we outlined depends on the level and variability of gene expression and quantity of RNA collected, our results should be interpreted alongside these limitations. BDE congener analysis was performed using two mass spectrometry methods utilized by teams at different institutions. The GC/ECNI-MS method uses an ECNI ionization mode to improve sensitivity. This method provides equal sensitivity to HRGC/HRMS that uses electron impact ionization. Therefore, the reduction in brain BDE congener at PND110 is likely due to elimination and not to methodological factors. All social behavioral tests were analyzed using litter as the unit of statistical analysis. However, for practical reasons most others tests used individual subjects. Our findings pertain to exposed female offspring and their mothers but male offspring were omitted due to limited resources. Further research is needed to determine if the ASD phenotypes evoked using the PBDE model are sex-specific.

## Acknowledgements

We acknowledge Drs. G. Hicks, M. Collin, D. Carter and C. and H. Clark at UCR Institute for Integrative Genome Biology, Genomics and Imaging Cores, and E. Grace (IDT) and M. Kuhn (NEB) for advice on PCR analysis. We thank UCR graduate students R. Bottom, K. Conner and D. Rohac for assistance with Suok and FST. We thank Drs. K. Huffman (Dept. Psychology), W. Saltzman (Dept. Biology) for access to behavioral apparatus. We are grateful to Dr. B. Wong (Noldus) for Ethovision software training. Dr. J. Porter and M. Colon at the Brain Behavioral Core (RR003050/MD007579), Ponce Health Sciences University, provided additional support on Ethovision data analysis. We thank B. O’Hara and R. Hart (Arbor Assays) for advice on EIA analysis and Dr. M. Adams for the use of speed vacuum evaporator. We thank J. Phan for help with animal husbandry. We thank Jim from GraphPad Support for helpful software assistance. We are grateful to Drs. I. Ethell, F. Sladek, K. Huffman, M. Riccomagno for gift of mice used as breeders and stimulus animals. Illustrations were created with BioRender.com. The authors also thank Drs. Michael Hughes and Andrew Johnstone of USEPA for their helpful comments on an earlier version of this manuscript.

## Dedication

This report is dedicated to Dr. Elizabeth R. Gillard, who set us on the path to study the neurotoxicity of endocrine disrupting chemicals and who embodied an intense passion for discovery. Moreover, she cultivated an inclusive culture and herculean work ethic that has promoted the highest standards for excellence in the lab. We immortalize her memory here.

## Declarations

### Disclaimer

J.M.K. is now a 2nd Lieutenant at the Uniformed Services University, Department of Defense. Her work was performed at the University of California, Riverside before becoming a military officer. However, we want to emphasize that the opinions and assertions expressed herein are those of the authors and do not necessarily reflect the official policy or position of the Uniformed Services University or the Department of Defense.

The research described in this article has been reviewed by the Center for Public Health and Environmental Assessment, U.S. Environmental Protection Agency (EPA) and approved for publication. Approval does not signify that the contents necessarily reflect the views and policies of the agency nor does the mention of trade names of commercial products constitute endorsement or recommendation for use.

## Availability of Data and Material

Not applicable.

## Code Availability

Not applicable.

## CRediT authorship contribution statement

**Elena V. Kozlova:** Conceptualization, Data curation, Formal Analysis, Funding acquisition, Investigation, Methodology, Project administration, Software, Supervision, Validation, Visualization, Writing – original draft, Writing – review & editing. **Matthew C. Valdez**: Conceptualization, Data curation, Formal Analysis, Funding acquisition, Investigation, Methodology, Project administration, Software, Supervision, Validation. **Maximilian E. Denys:** Formal Analysis, Funding Acquisition Investigation, Software, Writing – original draft. **Anthony E. Bishay:** Formal Analysis, Funding Acquisition, Investigation, Writing – original draft. **Julia M. Krum**: Data curation, Funding Acquisition, Investigation, Methodology, Software, Visualization. **Kayhon M. Rabbani:** Formal Analysis, Funding Acquisition, Investigation, Software, Validation, Data curation. **Valeria Carrillo:** Investigation, Funding Acquisition, Data curation. **Gwen M. Gonzalez:** Funding acquisition, Investigation, Methodology, Validation. **Jasmin D. Tran:** Formal Analysis, Investigation, Funding acquisition. **Brigitte M. Vazquez:** Investigation, Funding Acquisition. **Gregory Lampel:** Investigation, Funding Acquisition. **Laura M. Anchondo:** Investigation, Software. **Syed A. Uddin:** Investigation, Software, Validation. **Nicole M. Huffman:** Investigation, Software, Validation. **Eduardo Monarrez:** Investigation, Data curation, Software, Validation. **Duraan S. Olomi:** Investigation, Data curation. **Bhuvaneswari D. Chinthirla:** Investigation. **Richard E. Hartman:** Resources, Software, Methodology, Validation, Writing – review & editing. **Prasada Rao S. Kodavanti:** Funding acquisition, Resources, Writing – review & editing. **Gladys Chompre:** Investigation. **Allison L. Phillips:** Formal Analysis, Investigation, Writing – review & editing. **Heather M. Stapleton**: Formal Analysis, Funding acquisition, Methodology, Resources, Supervision, Validation, Writing – review & editing. **Bernhard Henkelmann:** Investigation, Methodology, Validation, Writing - original draft. **Karl-Werner Schramm:** Methodology, Resources, Funding acquisition, Supervision, Writing – review & editing. **Margarita C. Curras-Collazo:** Conceptualization, Formal Analysis, Funding acquisition, Methodology, Project administration, Resources, Supervision, Validation, Visualization, Writing – original draft, Writing – review & editing

## Ethics approval

Care and treatment of animals was performed in accordance with guidelines from and approved by the University of California, Riverside Institutional Animal Care and Use Committee (AUP #00170026 and 20200018).

## Consent to participate

Not applicable.

## Consent for publication

All authors reviewed and approved the final manuscript.

