## Supplementary data 1 for "Persistent autism-relevant phenotype produced by *in utero* and lactational exposure of female mice to the commercial PBDE mixture, DE-71"

Orchid IDs: E.V.K.: 0000-0002-4691-6618; A.E.B.: 0000-0002-6057-3770; K.W.S.: 0000-0002-8945-4062; M.C.C.: 0000-0002-0189-4179

[1]Department of Molecular, Cell and Systems Biology, University of California, Riverside, CA 92521, USA

[2] Neuroscience Graduate Program, University of California, Riverside, CA, 92521, USA

[3] Duke University, Nicholas School of the Environment, Durham, NC 27710, USA

[4]Department of Psychology, Loma Linda University, Loma Linda CA 92350, USA

[5]Biotechnology Department, Pontifical Catholic University of Puerto Rico, Ponce, Puerto Rico 00717-9997 USA.

[6] HelmholtzZentrum Munchen, German National Research Centre for Environmental Health (GmbH), Molecular EXposomics (MEX), Ingolstaedter Landstrasse 1, Neuherberg, Munich, Germany

[7]TUM, Wissenschaftszentrum Weihenstephan für Ernährung, Landnutzung und Umwelt, Department für Biowissenschaftliche Grundlagen, Weihenstephaner Steig 23, 85350 Freising, Germany

[8]Neurological and Endocrine Toxicology Branch, Public Health and Integrated Toxicology Division, CPHEA/ORD, U.S. Environmental Protection Agency, Research Triangle Park, NC 27711 USA

***Corresponding author**:

Dr. Margarita C. Curras-Collazo, Ph.D

Professor of Neuroscience

Department Molecular, Cell and Systems Biology

University of California, Riverside

Riverside, CA 92521

951-827-3960

**Declarations**

**Funding**

We acknowledge funding from UCR Committee on Research (CoR) Grants to M.C.C.; UC MEXUS Awards to M.C.C., E.V.K., M.C.V.; NSF GRFP to M.C.V.; MARC U STAR Fellowship and NIH T34 (T34GM062756) to G.M.G.; UCR GRMP to E.V.K.; Sigma Xi Grant-in-Aid of Research award to E.V.K., K.M.R., M.E.D.; UCR Undergraduate Minigrant to E.V.K., K.M.R., A.E.B., V.C., G.L., B.M.V.; STEM-HSI Department of Education Award to E.V.K.; UCR

Chancellor's Fellowship to J.M.K., APS IOSP Scholarship to L.M.A, APS STRIDE to A.E.B., and NIH R01 ES016099 to H.M.S.

**Conflicts of interests/Competing interests**

The authors report no conflicts of interests and have no competing interests to declare.

**Disclaimer:** J.M.K. is now a 2nd Lieutenant at the Uniformed Services University, Department of Defense. Her work was performed at the University of California, Riverside before becoming a military officer. However, we want to emphasize that the opinions and assertions expressed herein are those of the authors and do not necessarily reflect the official policy or position of the Uniformed Services University or the Department of Defense.

The research described in this article has been reviewed by the Center for Public Health and Environmental Assessment, U.S. Environmental Protection Agency (EPA) and approved for publication. Approval does not signify that the contents necessarily reflect the views and policies of the agency nor does the mention of trade names of commercial products constitute endorsement or recommendation for use.

**Availability of Data and Material**

Not applicable.

**Code Availability**

Not applicable.

**CRediT authorship contribution statement**

**Elena V. Kozlova:** Conceptualization, Data curation, Formal Analysis, Funding acquisition, Investigation, Methodology, Project administration, Software, Supervision, Validation,

Visualization, Writing – original draft, Writing – review & editing. **Matthew C. Valdez**: Conceptualization, Data curation, Formal Analysis, Funding acquisition, Investigation, Methodology, Project administration, Software, Supervision, Validation. **Maximilian E. Denys:** Formal Analysis, Funding Acquisition Investigation, Software, Writing – original draft. **Anthony E. Bishay:** Formal Analysis, Funding Acquisition, Investigation, Writing – original draft. **Julia M. Krum**: Data curation, Funding Acquisition, Investigation, Methodology, Software, Visualization. **Kayhon M. Rabbani:** Formal Analysis, Funding Acquisition, Investigation, Software, Validation, Data curation. **Valeria Carrillo:** Investigation, Funding Acquisition, Data curation. **Gwen M. Gonzalez:** Funding acquisition, Investigation, Methodology, Validation. **Jasmin D. Tran:** Formal Analysis, Investigation, Funding acquisition. **Brigitte M. Vazquez:** Investigation, Funding Acquisition. **Gregory Lampel:** Investigation, Funding Acquisition. **Laura M. Anchondo:** Investigation, Software. **Syed A. Uddin:** Investigation, Software, Validation. **Nicole M. Huffman:** Investigation, Software, Validation. **Eduardo Monarrez:** Investigation, Data curation, Software, Validation. **Duraan S. Olomi:** Investigation, Data curation. **Bhuvaneswari D. Chinthirla:** Investigation. **Richard E. Hartman:** Resources, Software, Methodology, Validation, Writing – review & editing. **Prasada Rao S. Kodavanti:** Funding acquisition, Resources, Writing – review & editing. **Gladys Chompre:** Investigation. **Allison L. Phillips:** Formal Analysis, Investigation, Writing – review & editing. **Heather M. Stapleton**: Formal Analysis, Funding acquisition, Methodology, Resources, Supervision, Validation, Writing – review & editing. **Bernhard Henkelmann:** Investigation, Methodology, Validation, Writing - original draft. **Karl-Werner Schramm:** Methodology, Resources, Funding acquisition, Supervision, Writing – review & editing. **Margarita C. Curras-Collazo:** Conceptualization,

Formal Analysis, Funding acquisition, Methodology, Project administration, Resources, Supervision, Validation, Visualization, Writing – original draft, Writing – review & editing

**Ethics approval**

Care and treatment of animals was performed in accordance with guidelines from and approved by the University of California, Riverside Institutional Animal Care and Use Committee (AUP #00170026 and 20200018).

**Consent to participate**

Not applicable.

**Consent for publication**

All authors reviewed and approved the final manuscript.

**Supplementary Table 1.**

GC/MS parameters for the isomer specific detection of PBDE.

| | |
|---|---|
| GC Type | Agilent 6890 |
| Column | Rtx-1614, 15 m, 0.25 mm ID, 0.1 µm film thickness (Restek) |
| Temperature program | 75 °C, 1.5 min, 18 °C/min, 210 °C, 8 °C/min, 310 °C, 5 min |
| Carrier gas | helium |
| Flow | constant flow 1.6 mL/min |
| Injector | cooled injection system CIS 4 (Gerstel) |
| Temperature transfer line | 320 °C |
| Injection volume | 1 µL splitless |
| MS Type | MAT 95XL (Thermo) |
| Ionization mode | EI+, 47 eV, 260°C |
| Resolution | >9000 |

**Supplementary Table 2.**
Limits of quantification for PND 15 samples.

| | Mean | Standard deviation |
|---|---|---|
| | μg/kg wet weight | μg/kg wet weight |
| BDE-7 | 0.008 | 0.006 |
| BDE-10 | 0.006 | 0.005 |
| BDE-15 | 0.15 | 0.04 |
| BDE-17 | 0.008 | 0.002 |
| BDE-28 | 0.15 | 0.04 |
| BDE-30 | 0.01 | 0.01 |
| BDE-47 | 1.5 | 0.37 |
| BDE-49 | 0.02 | 0.006 |
| BDE-66 | 0.06 | 0.05 |
| BDE-71 | 0.02 | 0.004 |
| BDE-77 | 0.05 | 0.04 |
| BDE-85 | 0.05 | 0.04 |
| BDE-99 | 0.95 | 0.23 |
| BDE-100 | 0.43 | 0.10 |
| BDE-119 | 0.05 | 0.01 |
| BDE-126 | 0.05 | 0.04 |
| BDE-138 | 0.07 | 0.06 |
| BDE-139 | 0.06 | 0.05 |
| BDE-140 | 0.06 | 0.05 |
| BDE-153 | 0.75 | 0.18 |
| BDE-154 | 0.05 | 0.04 |
| BDE-156 | 0.07 | 0.06 |
| BDE-171 | 0.12 | 0.05 |
| BDE-180 | 0.12 | 0.05 |
| BDE-183 | 0.09 | 0.04 |
| BDE-184 | 0.09 | 0.04 |
| BDE-191 | 0.09 | 0.04 |
| BDE-196 | 0.02 | 0.02 |
| BDE-197 | 0.02 | 0.02 |
| BDE-201 | 0.02 | 0.02 |
| BDE-203 | 0.02 | 0.02 |
| BDE-204 | 0.02 | 0.02 |
| BDE-205 | 0.03 | 0.04 |
| BDE-206 | 0.31 | 0.07 |
| BDE-207 | 0.36 | 0.09 |
| BDE-208 | 0.21 | 0.05 |
| BDE-209 | 5.0 | 1.2 |

Content was measured as ng/g wet weight
Values are expressed as ug/kg wet weight
Values expressed as mean ± S.D.
BDE, brominated diphenyl ether
Values ≥ 0.15 indicate detected blanks

**Supplementary Table 3.** Mass spectrometric analysis (HRGC/HRMS) of PBDE congeners in PND 15 F1 female offspring brain on a wet-weight basis after transplacental and lactational low dose exposure to DE-71 through the dam. (Related to Figure 1)

| Compound/Substituents | IUPAC Number | Female Offspring Brain PND 15 | | |
|---|---|---|---|---|
| Treatment | | VEH/CON | 0.1 mg/kg DE-71 | 0.4 mg/kg DE-71 |
| n | | 3 | 3 | 3 |
| 2,4-Dibromdiphenylether | BDE-7 | <LOQ | 0.03 | <LOQ |
| 2,6-Dibromdiphenylether | BDE-10 | <LOQ | <LOQ | <LOQ |
| 4,4'-Dibromdiphenylether | BDE-15 | <LOQ | <LOQ | <LOQ |
| 2,2',4-Tribromdiphenylether | BDE-17 | 0.007 | 0.01 | 0.02±0.003* |
| 2,4,4'-Tribromdiphenylether | BDE-28 | <LOQ | <LOQ | 0.37±0.11* |
| 2,4,6-Tribromdiphenylether | BDE-30 | <LOQ | 0.02 | <LOQ |
| 2,2',4,4'-Tetrabromdiphenylether | BDE-47 | <LOQ | 9.51±3.12* | 51.6±12.1* |
| 2,2',4,5'-Tetrabromdiphenylether | BDE-49 | 0.025 | <LOQ | 0.07 |
| 2,3',4,4'-Tetrabromdiphenylether | BDE-66 | <LOQ | <LOQ | <LOQ |
| 2,3',4',6-Tetrabromdiphenylether | BDE-71 | <LOQ | <LOQ | <LOQ |
| 3,3',4,4'-Tetrabromdiphenylether | BDE-77 | <LOQ | <LOQ | <LOQ |
| 2,2',3,4,4'-Pentabromdiphenylether | BDE-85 | 0.02±0.002 | 1.27±0.23 | 4.17±1.58 |
| 2,2',4,4',5-Pentabromdiphenylether | BDE-99 | <LOQ | 28.3±7.7* | 100.5±24.4* |
| 2,2',4,4',6-Pentabromdiphenylether | BDE-100 | <LOQ | 9.52±2.87* | 33.4±7.7* |
| 2,3',4,4',6-Pentabromdiphenylether | BDE-119 | 0.19 | <LOQ | <LOQ |
| 3,3',4,4',5-Pentabromdiphenylether | BDE-126 | <LOQ | <LOQ | <LOQ |
| 2,2',3,4,4',5'-Hexabromdiphenylether | BDE-138 | <LOQ | 0.45±0.08* | 1.44±0.33* |
| 2,2',3,4,4',6-Hexabromdiphenylether | BDE-139 | 0.03 | 2.27±0.66* | 7.99±1.67* |
| 2,2',3,4,4',6'-Hexabromdiphenylether | BDE-140 | <LOQ | 0.32±0.07* | 0.85±0.21* |
| 2,2',4,4',5,5'-Hexabromdiphenylether | BDE-153 | <LOQ | 23.6±6.80* | 87.2±16.5* |
| 2,2',4,4',5,6'-Hexabromdiphenylether | BDE-154 | 0.05±.01 | 2.44±0.42* | 7.32±1.55* |
| 2,3,3',4,4',5-Hexabromdiphenylether | BDE-156 | <LOQ | <LOQ | 0.05 |
| 2,2',3,3',4,4',6-Heptabromdiphenylether | BDE-171 | <LOQ | <LOQ | <LOQ |
| 2,2',3,4,4',5,5'-Heptabromdiphenylether | BDE-180 | <LOQ | <LOQ | 0.13 |
| 2,2',3,4,4',5',6-Heptabromdiphenylether | BDE-183 | 0.078 | 0.19±0.03* | 0.50±0.06** |
| 2,2',3,4,4',6,6'-Heptabromdiphenylether | BDE-184 | <LOQ | 0.19±0.02* | 0.54±0.08* |
| 2,3,3',4,4',5',6-Heptabromdiphenylether | BDE-191 | <LOQ | <LOQ | <LOQ |
| 2,2',3,3',4,4',5,6'-Octabromdiphenylether | BDE-196 | NR | NR | NR |
| 2,2',3,3',4,4',6,6'-Octabromdiphenylether | BDE-197 | NR | <LOQ | NR |
| 2,2',3,3',4,5',6,6'-Octabromdiphenylether | BDE-201 | NR | <LOQ | <LOQ |
| 2,2',3,4,4',5',5',6-Octabromdiphenylether | BDE-203 | NR | NR | <LOQ |
| 2,2',3,4,4',5',6,6'-Octabromdiphenylether | BDE-204 | <LOQ | <LOQ | <LOQ |
| 2,3,3',4,4',5,5',6-Octabromdiphenylether | BDE-205 | <LOQ | <LOQ | <LOQ |
| 2,2',3,3',4,4',5,5',6-Nonabromdiphenylether | BDE-206 | <LOQ | NR | <LOQ |
| 2,2',3,3',4,4',5,6,6'-Nonabromdiphenylether | BDE-207 | <LOQ | NR | <LOQ |
| 2,2',3,3',4,5,5',6,6'-Nonabromdiphenylether | BDE-208 | <NR | NR | <LOQ |
| 2,2',3,3',4,4',5,5',6,6'-Decabromdiphenylether | BDE-209 | NR | NR | <LOQ |
| ΣPBDEs (mean ± S.E.M.) | | 0.33±0.07 | 78.1±37.5** | 296±115*^ |

Content was measured as ng/g wet weight
Values expressed as mean ± S.E.M.
Values for all biological replicates are listed; all values were identical when <LOQ is listed
**P*<.05, ***P*<.01 compared to VEH/CON; ^*P*<.05 compared to L-DE-71 (Student's t-test with Welch's Correction)

‘n’ indicates the number of biological replicates per group.
BDE, brominated diphenyl ether; IUPAC, International Union of Pure and Applied Chemistry; LOQ, limit of quantification
NR, not repeatable, only 1 sample yielded detectable values

**Supplementary Table 4.** Mass spectrometric analysis (GC/ECNI/MS) of PBDE congeners in PND 110 F1 female offspring brain on a lipid-weight basis after transplacental and lactational low dose exposure to DE-71 through the dam. (Related to Figure 1)

| Compound/Substituents | IUPAC Number | Female Offspring Brain PND 110 | | |
|---|---|---|---|---|
| Treatment | | VEH/CON | 0.1 mg/kg DE-71 | 0.4 mg/kg DE-71 |
| n | | 4 | 4 | 4 |
| 2,2′,4-tri BDE | BDE 17 | <MDL | <MDL | <MDL |
| 2,3′,4-tri BDE | BDE 25 | <MDL | <MDL | <MDL |
| 2,4,4′-tri BDE, 2′,3,4-tri BDE | BDE 28, 33 | <MDL | <MDL | <MDL |
| 2,4,6,-tri BDE | BDE 30 | <MDL | <MDL | <MDL |
| 2,2′,4,4′-tetra BDE | BDE 47 | <MDL | <MDL | <MDL |
| 2,2′,4,5′-tetra BDE | BDE 49 | <MDL | <MDL | <MDL |
| 2,3′,4,4′-tetra BDE | BDE 66 | <MDL | <MDL | <MDL |
| 2,3′,4′,6-tetra BDE | BDE 71 | <MDL | <MDL | <MDL |
| 2,4,4′,6-tetra BDE | BDE 75 | <MDL | <MDL | <MDL |
| 2,2′,3,4,4′-penta BDE, 2,2′,4,4′,6,6′-hexa BDE | BDE 85, 155 | <MDL | <MDL | <MDL |
| 2,2′,4,4′,5-penta BDE | BDE 99 | <MDL | <MDL | <MDL |
| 2,2′,4,4′,6-penta BDE | BDE 100 | <MDL | <MDL | 13.0[#],<MDL,<MDL,<MDL |
| 2,3,4,5,6-penta BDE | BDE 116 | <MDL | <MDL | <MDL |
| 2,3′,4,4′,6 –penta BDE | BDE 119 | <MDL | <MDL | <MDL |
| 2,2′,3,4,4′,5′-hexa BDE | BDE 138 | <MDL | <MDL | <MDL |
| 2,2′,4,4′,5,5′-hexa BDE (mean ± S.E.M.) | BDE 153 | <MDL | 57.2, 179, 137, 79.9 (113 ± 27.5)* | 115, 185, 205,<MDL (126 ± 92.7)* |
| 2,2′,4,4′,5,6′-hexa BDE | BDE 154 | <MDL | <MDL | <MDL |
| 2, 3,3′,4,4′,5-hexa BDE | BDE 156 | <MDL | <MDL | <MDL |
| 2,2′, 3,4,4′,5,6-hepta BDE | BDE 181 | <MDL | <MDL | <MDL |
| 2,2′, 3, 4,4′,5′,6-hepta BDE | BDE 183 | <MDL | <MDL | <MDL |
| 2,3,3′,4,4′,5,6-hepta BDE | BDE 190 | <MDL | <MDL | <MDL |
| 2, 3,3′,4,4′,5′,6-hepta BDE | BDE 191 | <MDL | <MDL | <MDL |
| 2,2′,3,3′,4,5,6,6′-octa BDE, 2,2′,3,4,4′,5,5′,6-octa BDE | BDE 200, 203 | <MDL | <MDL | <MDL |
| 2, 3,3′,4,4′,5,5′,6-octa BDE | BDE 205 | <MDL | <MDL | <MDL |
| 2,2′,3,3′,4,4′,5,5′,6-nona BDE | BDE 206 | <MDL | <MDL | <MDL |
| 2,2′,3,3′,4,4′,5,5′,6,6′-deca BDE | BDE 209 | <MDL | <MDL | <MDL |
| ΣPBDEs | | <MDL | 113 ± 27.5* | 169 ± 27.3** |

Content was measured as ng/g lipid weight
Values expressed as mean ± S.E.M.
Values for all biological replicates are listed; all values were identical when <MDL is listed
*$P$<.05, **$P$<.01 compared to VEH/CON (Student's t-test with Welch's Correction)
### indicates values detected in one biological replicate were above MDL.
'n' indicates the number of biological replicates per group. BDE-100 was detected in only one 0.4 mg/kg DE-71 replicate.
BDE, brominated diphenyl ether; IUPAC, International Union of Pure and Applied Chemistry; MDL, method detection limit

**Supplementary Table 5.** Mass spectrometric analysis (GC/ECNI/MS) of PBDE congeners in PND 110 F1 female offspring brain on a wet-weight basis after transplacental and lactational low dose exposure to DE-71 through the dam. (Related to Figure 1)

| Compound/Substituents | IUPAC Number | Female Offspring Brain PND 110 | | |
|---|---|---|---|---|
| Treatment | | VEH/CON | 0.1 mg/kg DE-71 | 0.4 mg/kg DE-71 |
| n | | 4 | 4 | 4 |
| 2,2',4-tri BDE | BDE 17 | <MDL | <MDL | <MDL |
| 2,3',4-tri BDE | BDE 25 | <MDL | <MDL | <MDL |
| 2,4,4'-tri BDE, 2',3,4-tri BDE | BDE 28, 33 | <MDL | <MDL | <MDL |
| 2,4,6,-tri BDE | BDE 30 | <MDL | <MDL | <MDL |
| 2,2',4,4'-tetra BDE | BDE 47 | <MDL | <MDL | <MDL |
| 2,2',4,5'-tetra BDE | BDE 49 | <MDL | <MDL | <MDL |
| 2,3',4,4'-tetra BDE | BDE 66 | <MDL | <MDL | <MDL |
| 2,3',4',6-tetra BDE | BDE 71 | <MDL | <MDL | <MDL |
| 2,4,4',6-tetra BDE | BDE 75 | <MDL | <MDL | <MDL |
| 2,2',3,4,4'-penta BDE, 2,2',4,4',6,6'-hexa BDE | BDE 85, 155 | <MDL | <MDL | <MDL |
| 2,2',4,4',5-penta BDE | BDE 99 | <MDL | <MDL | <MDL |
| 2,2',4,4',6-penta BDE | BDE 100 | <MDL | <MDL | 0.19#, <MDL,<MDL,<MDL |
| 2,3,4,5,6-penta BDE | BDE 116 | <MDL | <MDL | <MDL |
| 2,3',4,4',6 –penta BDE | BDE 119 | <MDL | <MDL | <MDL |
| 2,2',3,4,4',5'-hexa BDE | BDE 138 | <MDL | <MDL | <MDL |
| 2,2',4,4',5,5'-hexa BDE (mean ± S.E.M.) | BDE 153 | <MDL | 0.61, 0.41, 0.64, 0.45 (0.53 ± 0.11)** | 1.71, 2.35, 1.91, <MDL (1.51 ± 1.0)* |
| 2,2',4,4',5,6'-hexa BDE | BDE 154 | <MDL | <MDL | <MDL |
| 2, 3,3',4,4',5-hexa BDE | BDE 156 | <MDL | <MDL | <MDL |
| 2,2', 3,4,4',5,6-hepta BDE | BDE 181 | <MDL | <MDL | <MDL |
| 2,2', 3, 4,4',5',6-hepta BDE | BDE 183 | <MDL | <MDL | <MDL |
| 2,3,3',4,4',5,6-hepta BDE | BDE 190 | <MDL | <MDL | <MDL |
| 2, 3,3',4,4',5',6-hepta BDE | BDE 191 | <MDL | <MDL | <MDL |
| 2,2',3,3',4,5,6,6'-octa BDE, 2,2',3,4,4',5,5',6-octa BDE | BDE 200, 203 | <MDL | <MDL | <MDL |
| 2, 3,3',4,4',5,5',6-octa BDE | BDE 205 | <MDL | <MDL | <MDL |
| 2,2',3,3',4,4',5,5',6-nona BDE | BDE 206 | <MDL | <MDL | <MDL |
| 2,2',3,3',4,4',5,5',6,6'-deca BDE | BDE 209 | <MDL | <MDL | <MDL |
| ΣPBDEs | | <MDL | 0.53 ± 0.06** | 1.51 ± 0.50* |

Content was measured as ng/g wet weight
Values expressed as mean ± S.E.M.
Values for all biological replicates are listed; all values were identical when <MDL is listed
**P*<.05, ***P*<.01 compared to VEH/CON (Student's t-test with Welch's Correction)
'n' indicates the number of biological replicates per group. BDE-100 was detected in only one 0.4 mg/kg DE-71 replicate.
### indicates values detected in one biological replicate were above MDL.
BDE, brominated diphenyl ether; IUPAC, International Union of Pure and Applied Chemistry; MDL, method detection limit

**Supplementary Figure 1.**

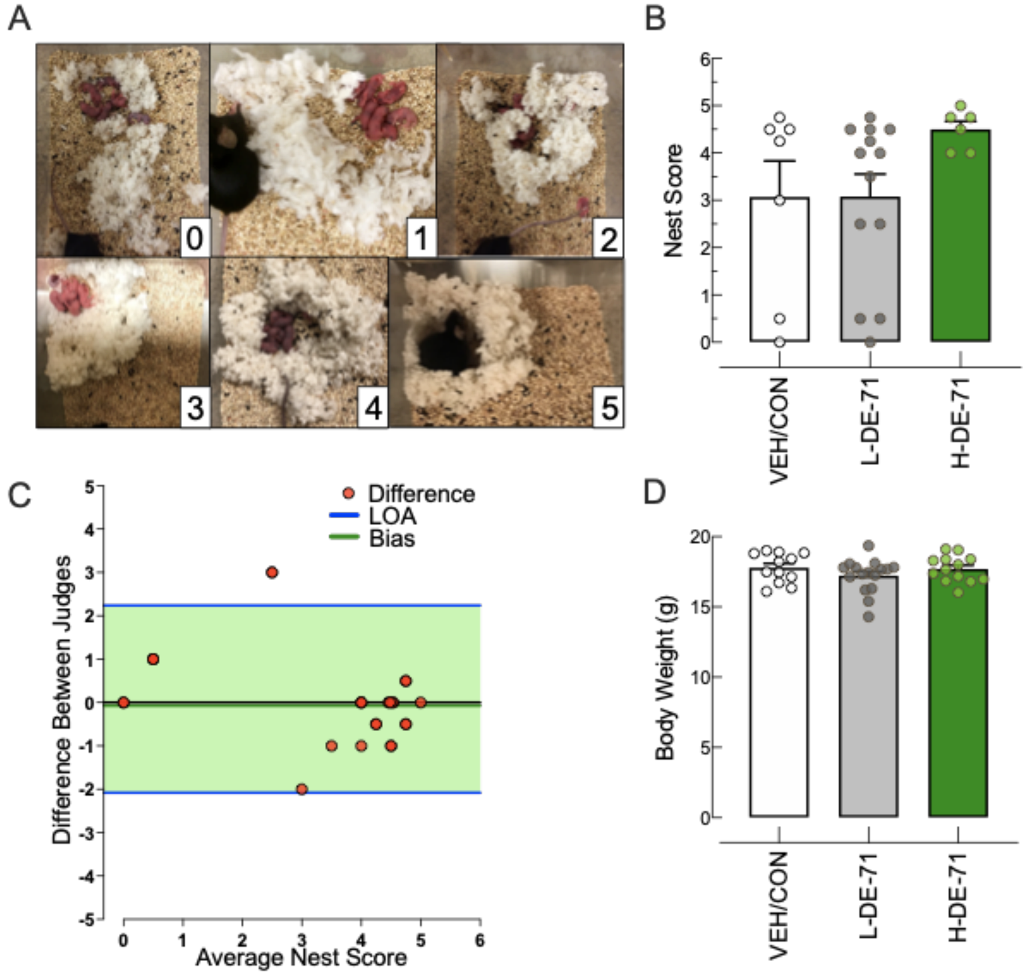


**Supplementary Figure 1.** Nest Scores of PBDE dams assessed on PND 0-1 and body weights of perinatally exposed female juvenile offspring (PND46). (Related to Figure 1) **a** Representative images for the nests receiving a score of 0-5 using established criteria. **b** Scores of nests (mean +/- s.e.m) built by dams are not different across exposure groups. **c** Bland-Altman bias plot (mean$\pm$s.d.) was used to test the validity and reproducibility between two independent judges blind to exposure group. Analysis revealed a very small mean of the differences between judge scores (Bias, 0.08±1.1) and a precision measured as limits of agreement (LOA), average difference ± 1.96 standard deviation of the difference, of -2.1-2.2, indicating negligible skewing by either judge. **d** Average body weights (mean +/- s.e.m) of female offspring litters at PND 46. b *n*=4-13 nests/group; d *n*=12-16 litters/group.

**Supplementary Figure 2.**

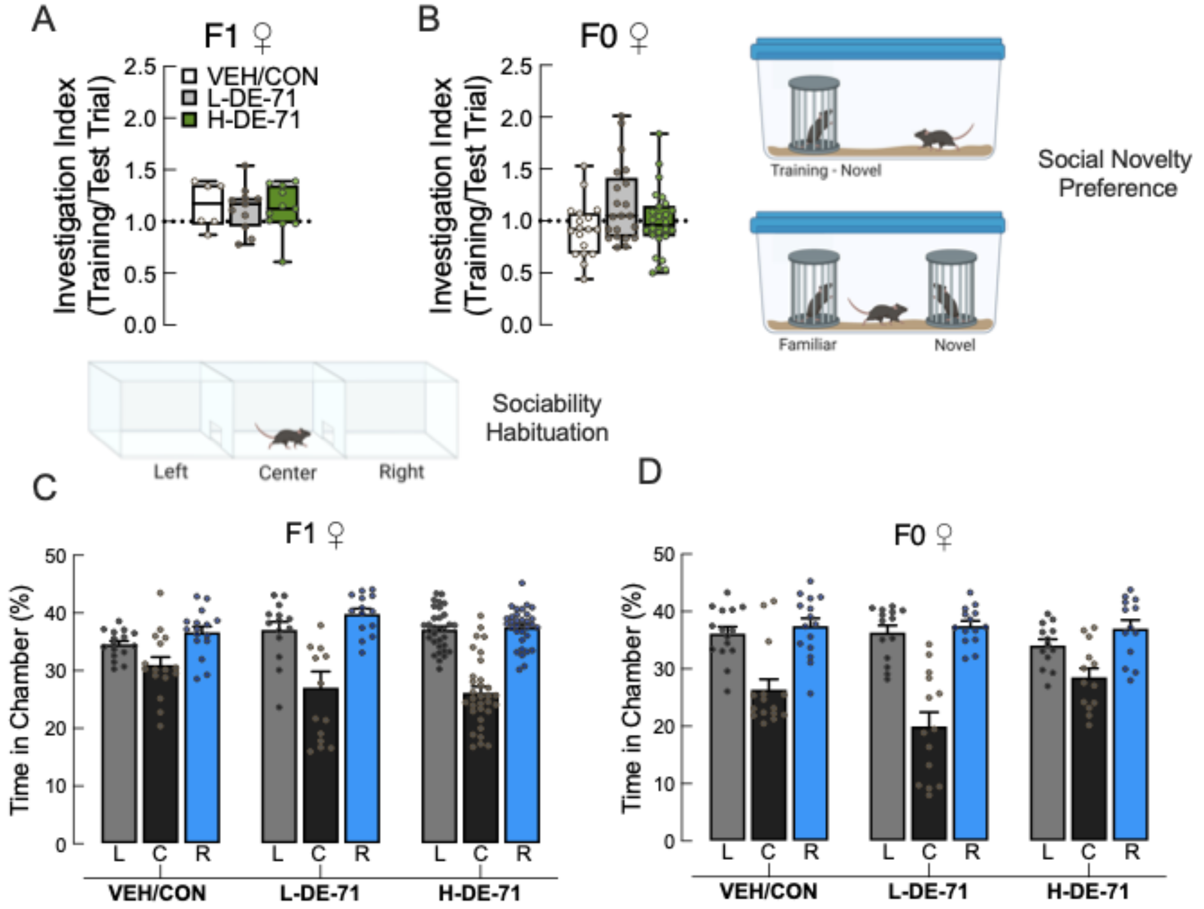


**Supplementary Figure 2.** Scores for investigation index on social novelty preference test (SNP) and time in chamber on the three-chamber sociability test (SOC) obtained prior to testing indicated mouse engagement in test and no inherent side preference, respectively**.** (Related to Figure 2) **a, b** Investigation Index scores on SNP indicate the ratio of time spent investigating the first social stimulus on training trial to the total investigation time during test trial (when exploring both social stimuli). A score of near 1 shown here indicates equal investigation time on training and test trials indicating that SNP deficit is not due to lack of social engagement. No effects of exposure on the investigation index was observed for F1 and F0. In SOC, adult female mice were tested for sociability in a 3-chamber apparatus. In the first habituation phase, mice are allowed to explore the middle chamber only. During the second habituation phase a test mouse is allowed to explore all three chambers of an empty apparatus and time spent in left, right and center chambers is recorded over 10 min. F1 (**c**) and F0 (**d**) females in all exposure groups showed similar times spent in left and right chambers during the second habituation phase indicating no inherent side preference. Sample size: F1: 6-11 litters/group (a), F0: 18-26 subjects/group (b), F1: 13-33 litters/group (c), F0: 13-24 subjects/group (d). L, left, C, center, R, right chambers

**Supplementary Figure 3.**

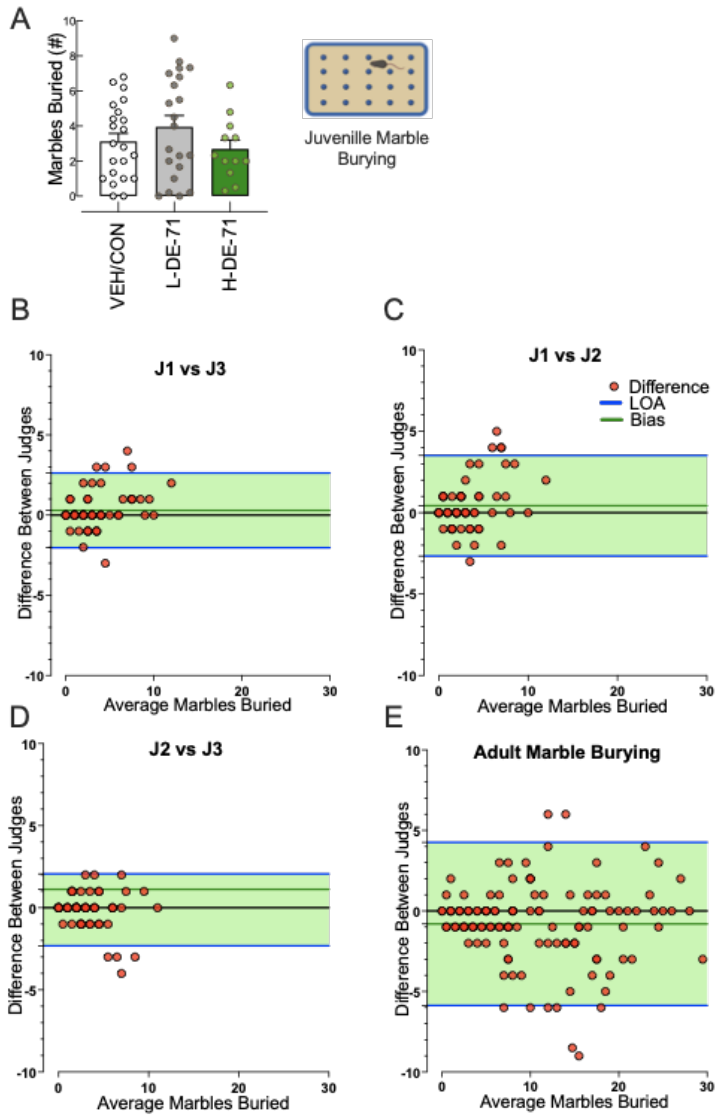


**Supplementary Figure 3.** Perinatal exposure to DE-71 does not alter repetitive behavior in female offspring tested as juveniles (PND35). (Related to Figure 2) **a** Mean scores for marbles buried were not significantly different across exposure groups. **b-d** Juvenile marble burying scores were determined by 3 judges (J) and inter-judge reliability assessed by Bland-Altman analysis (mean±s.d). Precision was measured as limits of agreement (LOA), where LOA is the average difference ± 1.96 standard deviation of the difference. (b) J1 vs J3, Bias: 0.30±1.2, LOA: -2.0-2.6. (c) J1 vs J2, Bias: 0.43±1.6, LOA: -2.7-3.5. (d) J2 vs J3, Bias: -0.13±1.1 and LOA: -2.3-2.1. (**e**) Adult marble burying scores were determined by 2 judges, Bias: -0.81±2.6 and LOA: -5.9-4.3. Outliers were removed if they fell outside the LOA between all three judges for b-d and two judges for e. a *n*=12-22 subjects/group.

**Supplementary Figure 4.**

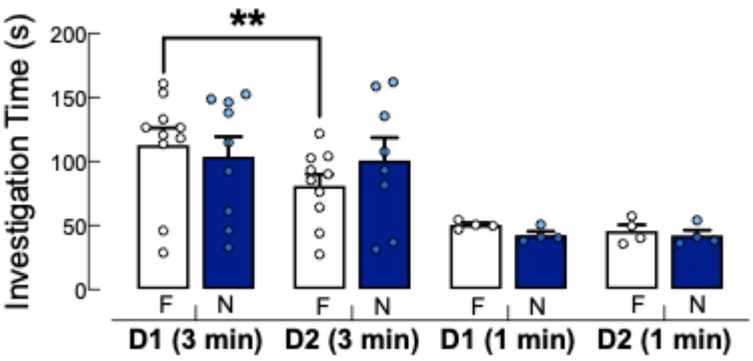


**Supplementary Figure 4.** (Related to Figure 3) Optimization experiments showed that long-term memory in F1 females, represented as reduced investigation of a familiar mouse after a 24 h retention, was achievable with a 3 but not 1 min social exposure on Day 1. *n*=4-10 subjects/group

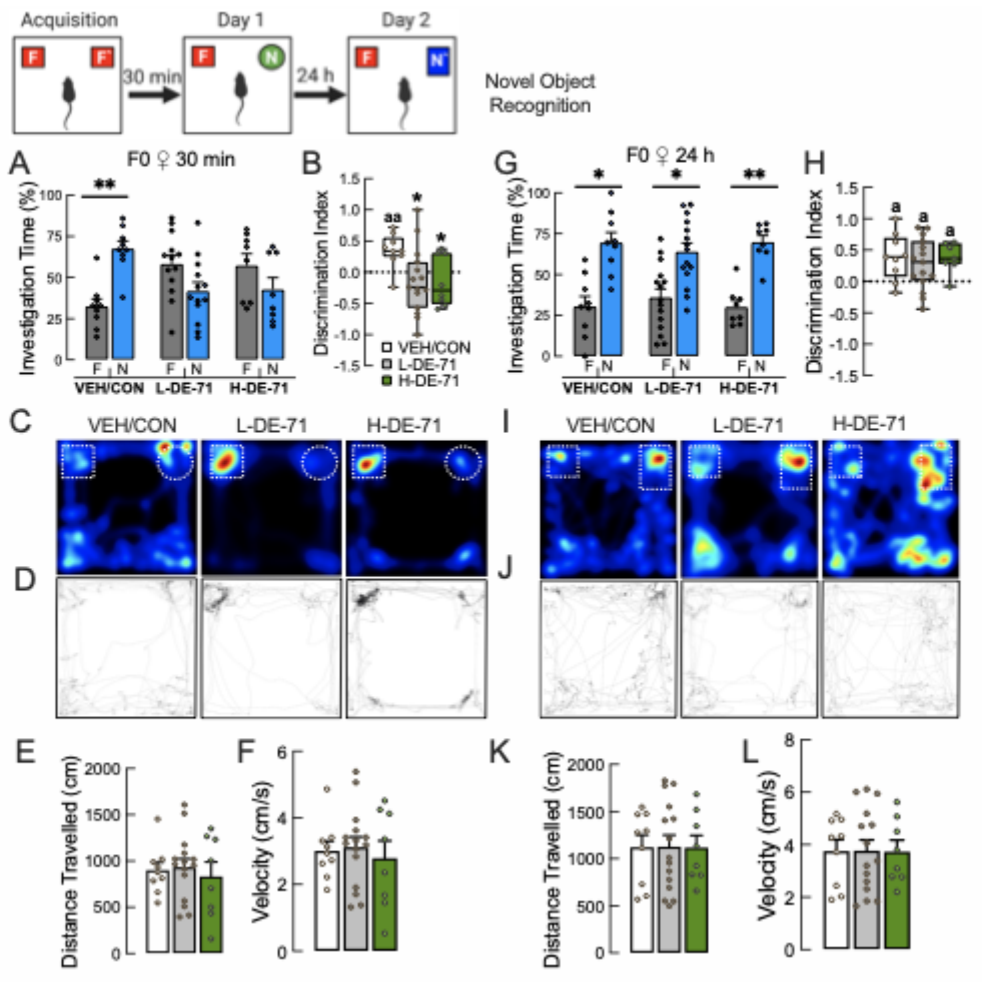


**Supplementary Figure 5.** Adult exposure to DE-71 reduces short-term but not long-term non-social memory in F0 dams. (Related to Figure 4) **a, g** Investigation time on NORT. **b, h** Discrimination index for 30 min and 24 h. **c, i** Representative heat maps (double gradient, blue—minus; red—plus) of time spent exploring novel and familiar objects placed in upper left and right corners of arena. **d, j** Representative raster plots. **e, k** Distance traveled in open field arena. **f, l** Velocity in open field arena.

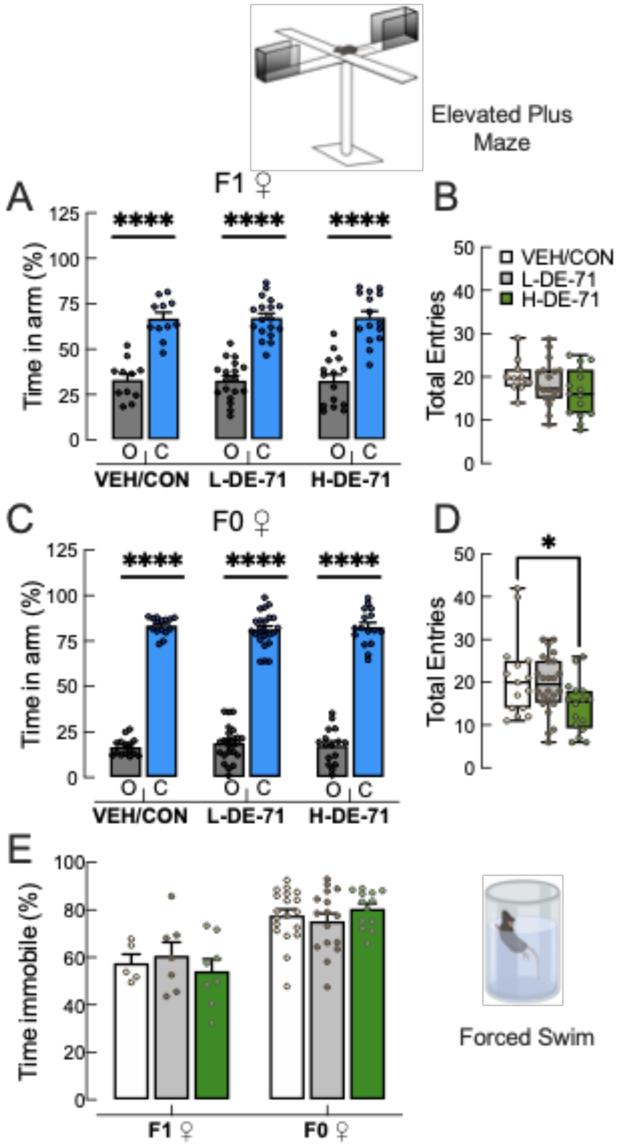


**Supplementary Figure 6.** DE-71 exposure does not affect anxiety nor produce depressive-like behavior in adult F1 nor F0. **a, c** Time spent in open and closed arms on an elevated plus maze. Both F1 and F0 spent significantly greater time in closed arm regardless of exposure. **b, d** Total entries into open and closed arms were similar across groups in F1 but H-DE-71 F0 group showed reduced total entries. **e** Time spent immobile on Forced Swim test showed no exposure effect. ****$P$<.0001 compared to closed arm time (a, c), (d). *compared to VEH/CON, *$P$<.05. $n$=11-18 litters/group, (a, b); 15-24 subjects/group (c, d); F1, $n$=5-8 litters/group, F0 $n$=13-19 subjects/group (e). O, open arm, C, closed arm

**Supplementary Figure 7.**

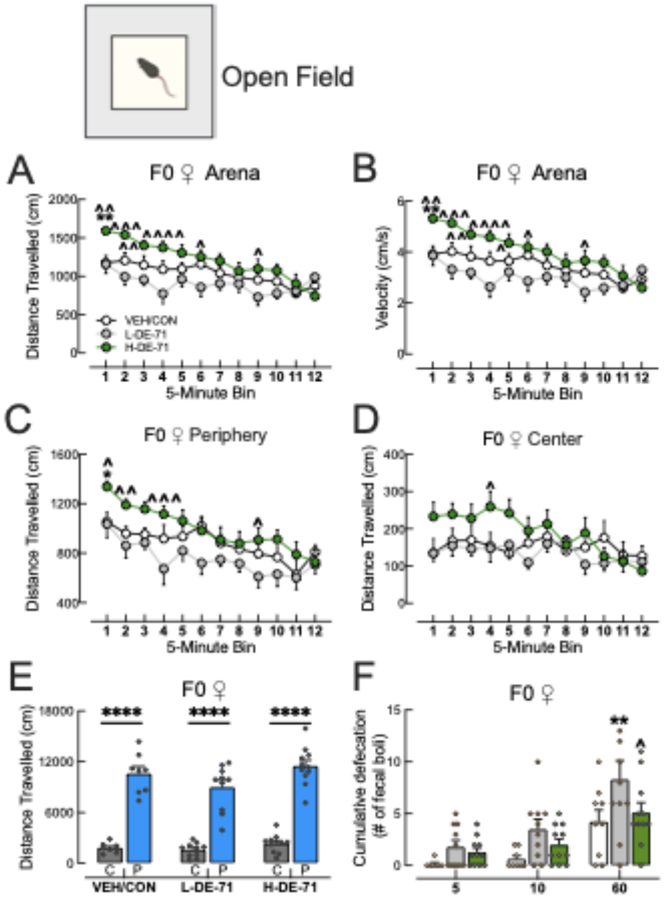


**Supplementary Figure 7.** Adult DE-71 exposure affects locomotion and anxiety on the Open Field Test (OFT). (Related to Figure 7**)**. H-DE-71 F0 exposure group showed increased exploratory activity measured by increased distance traveled (**a**) and velocity over the 1h-long test (**b)** indicating hyper-mobility relative to VEH/CON and L-DE-71. **c, d** Exploration time in periphery (but not center) of arena was significantly greater in H-DE-71 vs VEH/COn especially at early time points. **e** Group comparison of total exploration time in center was significantly less than in periphery. **f** Significant increase in the number of fecal boli indicated increased emotional reactivity in the L-DE-71 F1 relative to VEH/CON. *$P$<.05, **$P$<.01 compared to VEH/CON (a-d, f). ^$P$<.05, ^^$P$<.01, ^^^$P$<.01, ^^^^$P$<.0001 compared to corresponding L-DE-71 (a-d, f). ****$P$<.0001 compared to center (e). $n$=8-11 subjects/group. C, center zone; P, periphery zone

#### Supplementary Methods

##### *Elevated Plus Maze*

Anxiety was assessed using the elevated plus maze (20 x 22 cm; elevated 92 cm) (EPM), constructed from black Plexiglass as described (Lister, 1987). Two open arms and two closed arms received differential lighting, i.e., 300 and 30 LUX, respectively, to create anxiogenic (open) and anxiolytic conditions (closed). At the beginning of each trial, the test mouse was placed in the middle of the apparatus facing the open arm and was allowed to explore the apparatus for a total of 5 min. Activity was monitored by an overhead digital camera and video recordings were analyzed for time spent in each arm using BORIS.

##### *Forced Swim Test*

The forced swim test (FST) was used to assess depressive-like behavior using an apparatus consisting of a vertical Plexiglass cylinder (127 X 305 mm) filled to a depth of 150 mm with water maintained at 23-25°C (Lucki et al., 2001). Testing was conducted under dim room lighting conditions with an overhead light placed above the cylinders. Mice were placed individually into the water column for a 6 min test trial. After the test, mice were removed and dried before returning to their home cage. Swimming behavior was recorded using an overhead digital camera and video recordings were analyzed during the last 4 min using Ethovision. Active swimming was considered as time spent struggling/paddling with more than one limb. Time spent immobile, i.e., floating/absence of excessive movement and drifting or passive movements used to maintain verticality represented depressive-like behavior.

#### Supplementary Statistical Results

**Figure 1. Maternal dosing paradigm for DE-71 produces BDE congener penetration in adult female F1 offspring brain.** (**b**) $\Sigma_{14}$PBDEs in PND 15 F1 brain. One-way ANOVA, Exposure: $F_{(2,6)}$=14.3, *P*<.01. Tukey's post hoc, VEH/CON vs H-DE-71, *P*=.01, L-DE-71 vs H-DE-71, *P*=.05. $\sum_{14}$PBDE in PND 15 F1 brain. Welch's correction one-sided t-test, VEH/CON vs L-DE-71, $t_{(2)}$=3.5, *P*<.05; VEH/CON vs H-DE-71, $t_{(2)}$=4.4, *P*<.05. (**c**) $\sum$PBDE (BDE-153) in PND 110 F1 brain. Welch's ANOVA, Exposure: $F_{(2,4.3)}$=32.3, *P*<.01. Dunnet's T3 post-hoc: VEH/CON vs L-DE-71, *P*<.01. VEH/CON vs H-DE-71, ns. L-DE-71 vs H-DE-71, ns. Welch's one-sided t-test, VEH/CON vs L-DE-71, $t_{(3.5)}$=8.1, *P*<.01; VEH/CON vs H-DE-71, $t_{(3)}$=3.0, *P*<.05. (**e**) BDE-17. One-sided t-test, VEH/CON vs L-DE-71, $t_{(4)}$=1.0, ns; VEH/CON vs H-DE-71, $t_{(4)}$=2.7, *P*<.05; BDE-28. One-sided t-test, VEH/CON vs L-DE-71, $t_{(4)}$=0.49, ns; Welch's correction one-sided t-test, VEH/CON vs H-DE-71, $t_{(2.1)}$=2.7, *P*<.05; BDE-47. Welch's correction one-sided t-test, VEH/CON vs L-DE-71, $t_{(2.0)}$=2.7, *P*<.05; VEH/CON vs H-DE-71, $t_{(2.0)}$=4.2, *P*<.05. BDE-49. Welch's correction one-sided t-test, VEH/CON vs L-DE-71, $t_{(2.0)}$=1.3, ns; VEH/CON vs H-DE-71, $t_{(2.4)}$=1.8, *P*=.09. BDE-85. Welch's correction one-sided t-test, VEH/CON vs L-DE-71, $t_{(2)}$=5.5, *P*<.05; VEH/CON vs H-DE-71, $t_{(2)}$=2.6, *P*<.05. BDE-99. Welch's one-sided t-test, VEH/CON vs L-DE-71, $t_{(2)}$=3.6, *P*<.05; VEH/CON vs H-DE-71, $t_{(2)}$=4.1, *P*<.05. BDE-100. Welch's correction one-sided t-test, VEH/CON vs L-DE-71, $t_{(2)}$=3.3, *P*<.05; VEH/CON vs H-DE-

71, $t_{(2)}$=4.3, $P$<.05. BDE-138. Welch's correction one-sided t-test, VEH/CON vs L-DE-71, $t_{(2)}$=5.2, $P$<.05; VEH/CON vs H-DE-71, $t_{(2)}$=4.3, $P$<.05. BDE-139. Welch's correction one-sided t-test, VEH/CON vs L-DE-71, $t_{(2)}$=3.4, $P$<.05; VEH/CON vs H-DE-71, $t_{(4)}$=4.8, $P$<.05. BDE-140. Welch's correction one-sided t-test, VEH/CON vs L-DE-71, $t_{(2.3)}$=4.3, $P$<.05; VEH/CON vs H-DE-71, $t_{(2)}$=4.0, $P$<.05. BDE-153. Welch's correction one-sided t-test, VEH/CON vs L-DE-71, $t_{(2)}$=3.4, $P$<.05; VEH/CON vs H-DE-71, $t_{(2)}$=5.3, $P$<.05. BDE-154. Welch's correction one-sided t-test, VEH/CON vs L-DE-71, $t_{(2)}$=5.7, $P$<.05; VEH/CON vs H-DE-71, $t_{(2)}$=4.7, $P$<.05. BDE-183. Welch's correction one-sided t-test, VEH/CON vs L-DE-71, $t_{(2.4)}$=3.8, $P$<.05; VEH/CON vs H-DE-71, $t_{(2.1)}$=7.8, $P$<.01. BDE-184. Welch's correction one-sided t-test, VEH/CON vs L-DE-71, $t_{(2.1)}$=1.9, $P$=.09; VEH/CON vs H-DE-71, $t_{(2)}$=5.8, $P$<.05. (**f**) BDE 153. Welch's one-sided t-test, VEH/CON vs L-DE-71, $t_{(3)}$=9.0, $P$<.01; VEH/CON vs H-DE-71, $t_{(3)}$=3.0, $P$<.05.

**Figure 2**. **Early-life exposure to DE-71 produces deficits relevant to core symptoms of autism.** (**a**) F1 social novelty preference, Effect of stimulus. Paired t-test, VEH/CON: $t_{(5)}$=3.6, $P$<.05, Paired t-test, L-DE-71: $t_{(10)}$=1.5, ns. Paired t-test, H-DE-71: $t_{(10)}$=2.5, $P$<.05. (**b**) F1 Recognition Index. One-way ANOVA: $F_{(2,25)}$=2.77, $P$=.08. Dunnet's post-hoc test, VEH/CON vs L-DE-71, $P$<.05. (**c**) F0 social novelty preference, Effect of stimulus. Paired t-test, VEH/CON: $t_{(18)}$=13.5, $P$<.0001, Paired t-test, L-DE-71: $t_{(20)}$=6.9, $P$<.0001. Paired t-test, H-DE-71: $t_{(25)}$=7.5, $P$<.0001. (**e**) F0 Recognition Index. Brown-Forsythe ANOVA: $F_{(2,52.8)}$=1.3, ns. Dunnet's T3 post-hoc test, ns. (**e**) F1 Sociability, Sniffing Time, Effect of stimulus. Paired t-test, VEH/CON: $t_{(5)}$=3.7, $P$<.05, Paired t-test, L-DE-71, $t_{(6)}$=5.6, $P$<.01. Paired t-test, H-DE-71: $t_{(8)}$=2.9, $P$<.05. (**g**) F0 Sociability, Sniffing Time, Effect of stimulus. Paired t-test, VEH/CON: $t_{(7)}$=3.7, $P$<.01, Paired t-test, L-DE-71: $t_{(7)}$=3.6, $P$<.01. Paired t-test, H-DE-71: $t_{(7)}$=3.9, $P$<.01. (**g**) F1 Sociability, Time in Chamber. Paired t-test, VEH/CON: $t_{(15)}$=3.1, $P$<.01. L-DE-71 $t_{(12)}$=2.7, $P$<.05. H-DE-71: $t_{(32)}$=1.6, ns. (**h**) F0 Sociability, Time in Chamber. Paired t-test, VEH/CON: $t_{(12)}$=2.6, $P$<.05. L-DE-71 $t_{(16)}$=2.7, $P$<.05. H-DE-71: $t_{(23)}$=3.0, $P$<.01. (**i**) F1 Marble Burying. Brown-Forsythe ANOVA: $F_{(2,.6)}$=4.1, $P$<.05. Dunnet's T3 post-hoc test, VEH/CON vs L-DE-71, $P$<.05. F0 Marble Burying. One-way ANOVA: $F_{(2,39)}$=0.2, ns. (**j**) F1 Nestlet Square Shredding. One-way ANOVA: $F_{(2,85)}$=0.9, ns. F0 Nestlet Shredding. Brown Forsythe ANOVA: $F_{(2,39.1)}$=4.2, $P$<.05. Dunnet's T3 post-hoc test, VEH/CON vs L-DE-71, $P$<.05.

**Figure 3. Exposure to L-DE-71 but not H-DE-71 reduces long-term social recognition memory in F1.** (**a**) F1 Investigation Time Familiar Stimulus, Day 1 vs Day 2: VEH/CON paired t-test: $t_{(4)}$=3.7, $P$<.05; L-DE-71 $t_{(5)}$=2.0, ns; H-DE-71 $t_{(4)}$=16.7, $P$<.0001. (**b**) Recognition Index: One-Sample t- test comparing the mean vs a hypothetical value of 0.65: VEH/CON, $t_{(4)}$=1.1, ns., L-DE-71, $t_{(5)}$=2.3, $P$<.07; H-DE-71, $t_{(4)}$=3.9, $P$<.05; Across-groups: Brown-Forsythe ANOVA, $F_{(2,8.5)}$=5.7, $P$<.03. Dunnet's T3, VEH/CON vs H-DE-71, $P$<.05. (**c**) F1 Investigation Time Novel Stimulus, Day 1 vs Day 2: VEH/CON paired t-test: $t_{(4)}$=1.6, ns; L-DE-71 $t_{(4)}$=2.7, $P$<.05; H-DE-71 $t_{(4)}$=0.7, ns. (**d**) Recognition Index: One-Sample t-test comparing the mean vs a hypothetical value of 1.0: L-DE-71, $t_{(4)}$=2.8, $P$<.05; H-DE-71, $t_{(4)}$=.74, ns; VEH/CON, $t_{(4)}$=1.6, ns. Across-groups: One-Way ANOVA, $F_{(2,12)}$=0.2, ns.

**Figure 4. Perinatal exposure to DE-71 affects short- but not long-term novel object recognition memory. (a)** F1 30 min Investigation Time Familiar vs Novel: VEH/CON paired t-test: $t_{(3)}$=4.9, *P*<.01; L-DE-71 $t_{(9)}$=4, *P*<.01; H-DE-71 $t_{(5)}$=2.7, *P*<.05. (**b**) Discrimination Index: One-way ANOVA, Exposure: $F_{(2,17)}$=18.8, Tukey's post hoc, VEH/CON vs L-DE71 *P*<.001; L-DE-71 vs DE-71 *P*<.001. One-Sample t- test comparing the mean vs a hypothetical value of 0: VEH/CON, $t_{(3)}$=4.9, P<.05, L-DE-71, $t_{(9)}$=4.0, *P*<.01; H-DE-71, $t_{(5)}$=4.7, *P*<.05 (**e**) Distance travelled: One-way ANOVA, Exposure: $F_{(2,17)}$=1.7, ns**.** (**f**) Velocity: One-way ANOVA, Exposure: $F_{(2,17)}$=1.3, ns. (**g**) F1 24 h Investigation Time Familiar vs Novel: VEH/CON paired t-test: $t_{(3.1)}$=3.1, *P*<.05; L-DE-71 $t_{(9)}$=5.3, *P*<.001; H-DE-71 $t_{(5)}$=5.3, *P*<.0001. (**h**) Discrimination Index: One-way ANOVA, Exposure: $F_{(2,17)}$=1.9, ns. One-Sample t- test comparing the mean vs a hypothetical value of 0: VEH/CON, $t_{(3)}$=3.0, P<.05, L-DE-71, $t_{(9)}$=5.3, *P*<.001; H-DE-71, $t_{(5)}$=5.3, *P*<.01 (**k**) Distance travelled: One-way ANOVA, Exposure: $F_{(2,17)}$=.16, ns**.** (**l**) Velocity: One-way ANOVA, Exposure: $F_{(2,17)}$=2.0, ns.

**Figure 5. Perinatal exposure to DE-71 does not alter general olfaction but disrupts response to social odors.** (**a**) F1 Olfactory Preference Test: RM Mixed Effects Two-way ANOVA: Exposure $F_{(2,26)}$=2.0; Odor stimulus $F_{(3,77)}$=18.3, *P*<.0001; Interaction $F_{(6,77)}$=0.30; Tukey's post hoc test, odor stimulus: water vs peanut butter: VEH/CON *P*<.05, L-DE-71, *P*<.0001, H-DE-71, *P*<.05; peanut butter vs butyric acid: VEH/CON *P*<.01, L-DE-71, *P*<.001, H-DE-71, *P*<.001; peanut butter vs vanilla: VEH/CON *P*<.05, L-DE-71, *P*<.001, H-DE-71, *P*<.05; Between groups comparisons, exposure, ns. (**b**) F0 Olfactory Preference Test: RM Mixed Effects Two-way ANOVA: Exposure $F_{(1.7,64.7)}$=31.4, *P*<.001; Odor stimulus $F_{(2,38)}$=0.54, ns; Interaction $F_{(6,112)}$=0.41, ns; Tukey's post hoc test, odor stimulus: water vs peanut butter: VEH/CON *P*<.05, L-DE-71, *P*<.05, H-DE-71, *P*<.01; peanut butter vs butyric acid: VEH/CON *P*<.01, L-DE-71, *P*<.05, H-DE-71, *P*<.05; peanut butter vs vanilla: VEH/CON *P*<.05, L-DE-71, *P*<.01, H-DE-71, *P*<.05; Between groups comparison, exposure, ns. (**c**) F1 Olfactory Habituation/Dishabituation Test: RM Mixed Effects Two-way ANOVA: Odor stimulus trial $F_{(14,650)}$=64.7, *P*<.0001; Exposure $F_{(2,50)}$=0.94, ns; Interaction $F_{(28,650)}$=2.7, *P*<.0001; Sidak's post hoc test (see Table 1). Between groups comparisons, exposure (see Table 1). (**d**) F0 Olfactory Habituation/Dishabituation Test: RM Mixed Effects Two-way ANOVA: Odor stimulus trial $F_{(14,738)}$=59.3, *P*<.0001; Exposure $F_{(2,55)}$=1.1, ns; Interaction $F_{(28,738)}$=0.52, ns; Sidak's post hoc test (see Table 1). Between groups comparisons, exposure, ns.

**Figure 6. Selective effects of perinatal DE-71 on Suok Test**.

(**a**) F1 Horizontal Activity: One-way ANOVA, Exposure: $F_{(2,28)}$=5.5, $P$<.01, Tukey's post hoc test, VEH/CON vs H-DE-71, $P$<.05, L-DE-71 vs H-DE-71, $P$<.05; F0 Horizontal Activity: One-way ANOVA, Exposure: $F_{(2,30)}$=1.6, ns. (**b**) F1 Locomotor Activity: One-way ANOVA, Exposure: $F_{(2,27)}$=4.1, $P$<.05, Tukey's post hoc test, VEH/CON vs H-DE-71, $P$<.05, L-DE-71 vs H-DE-71, $P$<.05; F0 Locomotor Activity: One-way ANOVA, Exposure: $F_{(2,55)}$ =0.91, ns. (**C**) F1 Falls: One-way ANOVA, Exposure $F_{(2,26)}$=5.4, $P$<.05, Tukey's post hoc test, VEH/CON vs H-DE-71, $P$<.05. F0 Falls: One-way ANOVA, Exposure: $F_{(2,67)}$=0.58, ns. (**d**) F1 Falls per Segments Crossed: Kruskal-Wallis ANOVA, Exposure: $H_{(3)}$=9.6, $P$<.01. Dunn's post hoc test, L-DE-71 vs H-DE-71, $P$<.01. F0 Falls per Segments Crossed: One-way ANOVA, Exposure $F_{(2,47)}$=0.89, ns. (**e**) F1 Hind Leg Slips: One-way ANOVA, Exposure $F_{(2,31)}$=4.4, $P$<.05, Tukey's post hoc tests, L-DE-71 vs H-DE-71, $P$<.05. F0 Hind-Leg slips: Brown-Forsythe ANOVA, Exposure: $F_{(2,52.9)}$= 3.1, $P$<.05, Dunnet's T3 post hoc test, VEH/CON vs H-DE-71, $P$<.05. (**f**) F1 Hind Leg slips per segments crossed: One-way ANOVA, Exposure $F_{(2,30)}$=2.6, ns. Sidak's post hoc test, L-DE-71 vs H-DE-71, $P$<.05. F0 Hind Leg slips per segments crossed: One-way ANOVA, Exposure: $F_{(2,50)}$=0.59, ns. (**g**) F1 Head Dips and Side Looks: One-way ANOVA, Exposure $F_{(2,31)}$=2.6, ns. Sidak's post hoc test, VEH/CON vs H-DE-71, $P$<.05. F0 Head Dips and Side Looks: One-way ANOVA, Exposure $F_{(2,68)}$=0.27, ns. (**h**) F1 Latency to Leave Center: Kruskal-Wallis ANOVA, Exposure $H_{(6)}$=2.3, ns. F0 Latency to Leave Center: One-way ANOVA, Exposure $F_{(2,29)}$ =0.34, ns. (**i**) F1 Stretch and Attend Postures: One-way ANOVA, Exposure: $F_{(2,19)}$=3.0, ns. Sidak's post-hoc tests, VEH/CON vs H-DE-71, $P$<.05. F0 Stretch and Attend Postures: One-way ANOVA, Exposure: $F_{(2,59)}$=7.3, $P$<.01. Tukey's post hoc test, VEH/CON vs L-DE-71, $P$<.05, L-DE-71 vs H-DE-71, $P$<.01. (**j**) F1 Vegetative Responses: One-way ANOVA, Exposure: $F_{(2,27)}$=0.44, ns. F0 Vegetative Responses: One-way ANOVA, Exposure: $F_{(2,66)}$=0.24, ns. (**k**) F1 Autogrooming: One-way ANOVA, Exposure: $F_{(2,29)}$=6.3, $P$<.01, Tukey's post hoc test, VEH/CON vs H-DE-71, $P$<.05, L-DE-71 vs H-DE-71, $P$<.01. F0 Autogrooming: Kruskal-Wallis ANOVA, Exposure: $H_{(3)}$=2.7, ns.

**Figure 7. DE-71 exposure has minimal effects on locomotion and anxiety on the Open Field Test.** (**a**) F1 Distance Travelled Arena: RM Mixed Effects Two-way ANOVA: Exposure $F_{(2,60)}$=0.5, ns; Time $F_{(11,637)}$=51.5, $P$<.0001; Interaction $F_{(22,637)}$=1.5, ns; (**b**) F1 Velocity Arena: RM Mixed Effects Two-way ANOVA: Exposure $F_{(2,60)}$=0.47, ns; Time $F_{(11,651)}$=51.7, $P$<.0001; Interaction $F_{(2,651)}$=1.3, ns; (**c**) F1 Distance Travelled Periphery: RM Mixed Effects Two-way ANOVA: Exposure $F_{(2,60)}$=0.7, ns; Time $F_{(11,633)}$=66.1, $P$<.0001; Interaction $F_{(22,633)}$=1.2, ns; (**d**) F1 Distance Travelled Center: RM Mixed Effects Two-way ANOVA: Exposure $F_{(2,59)}$=0.69, ns; Time $F_{(11,626)}$=5.5, $P$<.0001; Interaction $F_{(22,626)}$=1.1, ns; (**e**) F1 Cumulative Distance Travelled Center vs Periphery: VEH/CON paired t-test: $t_{(22)}$=16.4, $P$<.0001; L-DE-71 $t_{(18)}$=12.7, $P$<.0001; H-DE-71 $t_{(20)}$=27.1, $P$<.0001. (**f**) F1 Cumulative Fecal Boli: Two-way ANOVA: Exposure $F_{(2,187)}$=8.5, $P$<.001; Time $F_{(2,187)}$=66.8, $P$<.0001; Interaction $F_{(4,187)}$=0.4, ns

**Figure 8. DE-71 alters gene expression in brain regions involved in social behavior.**
**BNST:** *Oxt*, One-way ANOVA, Exposure: $F_{(2,16)}$=4.6, P<.05. Dunnet's post hoc test, VEH/CON vs L-DE-71, $P$<.05; VEH/CON vs H-DE-71, $P$<.05. n=5-7/group. *Oxtr*, One-way ANOVA, Exposure: $F_{(2,18)}$=3.5, P<.05. Bonferroni's post hoc test, VEH/CON vs H-DE-71, $P$<.05. n=6-8/group. *Avp*, One-way ANOVA, Exposure: $F_{(2,19)}$=3.3, P<.05. Bonferroni's post hoc test, VEH/CON vs H-DE-71, $P$<.05. n=6-8/group. *Avp1ar*, One-way ANOVA, Exposure: $F_{(2,13)}$=2.7,

ns. Bonferroni's post hoc test, VEH/CON vs L-DE-71, *P*<.05. n=4-7/group. *Adcyap1*, One-way ANOVA, Exposure: $F_{(2,18)}$=0.79, ns. n=7/group *PAC1r*, One-way ANOVA, Exposure: $F_{(2,16)}$=.45, ns. n=6-7/group. **AMG:** *Oxt*, Kruskal-Wallis Test, Exposure: Exposure: H(2)=.46, ns. n=5-7/group. *Oxtr*, Welch's ANOVA: Exposure: $F_{(2,36.9)}$=3.3, P<.05. Dunnet's T3 post hoc test, VEH/CON vs H-DE-71, *P*<.05. n=15-17/group. *Avp*, One-way ANOVA, Exposure: $F_{(2,24)}$=.54, ns. n=7-10/group. *Avp1ar*, One-way ANOVA, Exposure: $F_{(2,13.5)}$=1.3, ns. n=9-14/group. *Adcyap1*, One-way ANOVA, Exposure: $F_{(2,46)}$=1.1, ns. n=16-17/group. *PAC1r*, One-way ANOVA, Exposure: $F_{(2,18)}$=.15, ns. n=7/group. **LS:** *Oxt*, One-way ANOVA, Exposure: $F_{(2,12)}$=2.0, ns. n=3-7/group. *Oxtr*, One-way ANOVA, Exposure: $F_{(2,32)}$=.43, ns. n=8-14/group. *Avp*, One-way ANOVA, Exposure: $F_{(2,12)}$=.18, ns. n=5/group. *Avp1ar*, One-way ANOVA, Exposure: $F_{(2,18)}$=2.0, ns. n=6-8/group. *Adcyap1*, Welch's ANOVA: Exposure: $F_{(2,20)}$=4.1, P<.05. Dunnet's T3 post hoc test, L-DE-71 vs H-DE-71, *P*<.05. *PAC1r*, One-way ANOVA, Exposure: $F_{(2,30)}$=.34, ns. n=9-12/group. **SON:** *Oxt*, One-way ANOVA, Exposure: $F_{(2,24)}$=2.5, ns. Bonferroni's post hoc test, VEH/CON vs L-DE-71, *P*<.05 n=9/group. *Oxtr*, One-way ANOVA, Exposure: $F_{(2,24)}$= 1.2, ns. n=8-10/group. *Avp*, One-way ANOVA, Exposure: $F_{(2,20)}$=5.1, P<.05. Tukey's post hoc test, VEH/CON vs H-DE-71, *P*<.05. n=7-9/group. *Avp1ar*, One-way ANOVA, Exposure: $F_{(2,23)}$=4.2, P<.05. Tukey's post hoc test, VEH/CON vs H-DE-71, *P*<.05. n=8-9/group. *Adcyap1*, One-way ANOVA, Exposure: $F_{(2,20)}$=.43, ns. n=6-9/group. *PAC1r*, One-way ANOVA, Exposure: $F_{(2,18)}$=1.5, ns. n=6-8/group. **PVN:** *Oxt*, One-way ANOVA, Exposure: $F_{(2,36)}$=1.2, ns. n=13/group. *Oxtr*, Brown-Forsythe ANOVA, Exposure: $F_{(2,21.1)}$=3.7, *P*<.05. Dunnet's T3 post hoc test, VEH/CON vs L-DE-71, *P*<.05. n=12-15/group. *Avp*, One-way ANOVA, Exposure: $F_{(2,40)}$=.32, ns. n=13-16/group. *Avp1ar*, One-way ANOVA, Exposure: $F_{(2,16)}$=.61, ns. n=6-7/group. *Adcyap1*, One-way ANOVA, Exposure: $F_{(2,46)}$=.74, ns. n=16-17/group. *PAC1r*, One-way ANOVA, Exposure: $F_{(2,43)}$=1.6, ns. n=14-17/group

**Figure 9. Perinatal exposure to DE-71 exaggerates plasma ARG8-vasopressin but not oxytocin levels in adult F1 female offspring.** (**a**) Plasma ARG8-vasopressin F1 offspring: One-way ANOVA, Exposure: $F_{(2,27)}$=2.2, *P*<.05, Dunnet's post-hoc test, VEH/CON vs L-DE-71 *P*<.05. (**b**) Plasma Oxytocin F1 offspring: One-way ANOVA, Exposure: $F_{(2,19)}$=0.34, ns.

**Supplementary Figure 1. Dam nest scores and offspring body weight:** One-way ANOVA, Exposure: $F_{(2,21)}$=1.1, ns. Body weights: One-way ANOVA, Exposure: $F_{(2,38)}$=1.1, ns.

**Supplementary Figure 2. Social novelty preference test (SNP) and time in chamber on the three-chamber sociability test (SOC) in F1 and F0**. (**a**) F1 SNP Investigation Index. One-way ANOVA: $F_{(2,25)}$=0.06, ns. (**b**) F0 SNP Investigation Index. One-way ANOVA: $F_{(2,62)}$=2.9, ns. (**c**) F1 Time in Left vs Right Chamber, VEH/CON: Paired t-test, $t_{(15)}$=1.6, ns. L-DE-71: $t_{(13)}$=1.7, ns. H-DE-71: $t_{(31)}$=0.5, ns. (**d**) F0 Time in Left vs Right Chamber, VEH/CON: Paired t-test, $t_{(14)}$=0.7, ns. L-DE-71: $t_{(13)}$=0.8, ns. H-DE-71: $t_{(12)}$=1.4, ns.

**Supplementary Figure 3. Repetitive behavior in juvenile and adult F1.**
(a) F1 juveniles, Marbles Buried: Brown-Forsythe ANOVA, Exposure: $F_{(2,45.3)}$=2.2, ns.

**Supplementary Figure 4. SMRT Test Optimization,** Investigation Time, Day 1 vs Day 2 Familiar, 3 min paired t-test: $t_{(9)}$=3.6, *P*<.01. Day 1 vs Day 2 Novel, 3 min paired t-test: $t_{(7)}$=.38, ns. Day 1 vs Day 2 Familiar, 1 min paired t-test: $t_{(3)}$=1.1, ns. Day 1 vs Day 2 Novel, 1 min paired t-test: $t_{(3)}$=.03, ns.

**Supplementary Figure 5. NORT in F0.** (**a**) F0 30 min Investigation Time Familiar vs Novel: VEH/CON paired t-test: $t_{(8)}$=3.8, *P*<.01; L-DE-71 $t_{(14)}$=1.0, ns; H-DE-71 $t_{(8)}$=0.4, ns. (**b**) Discrimination Index: One-way ANOVA, Exposure: $F_{(2,29)}$=4.1, *P*<.05. (**c**) Representative heat maps (double gradient, blue—minus; red—plus) of time spent exploring novel and familiar objects located in upper left and right corners of open field arena (dashed), respectively, showed differences in dwell times for different exposure groups for 30 min test. (**d**) Representative raster plots indicate no significant effect of exposure on general locomotor activity. (**e**) F0 Distance travelled: One-way ANOVA, Exposure: $F_{(2,29)}$=0.21, ns**.** (**f**) F0 Velocity: One-way ANOVA, Exposure: $F_{(2,29)}$=0.21, ns. (**g**) F0 24 h Investigation Time Familiar vs Novel: VEH/CON paired t-test: $t_{(8)}$=3.1, *P*<.05; L-DE-71 $t_{(14)}$=2.7, *P*<.05; H-DE-71 $t_{(7)}$=4.8, *P*<.01. (**h**) F0 Discrimination Index: One-way ANOVA, Exposure: $F_{(2,29)}$=0.25, ns. (**i**) heatmaps (**j**) raster plots (**k**) F0 Distance travelled: One-way ANOVA, Exposure: $F_{(2,29)}$=0.002, ns**.** (**l**) F0 Velocity: One-way ANOVA, Exposure: $F_{(2,29)}$=.002, ns. [a]Significantly different from 0, [a]*P*<.05, [aa]*P*<.01. F, familiar object; N and N`, novel object.

**Supplementary Figure 6. Elevated plus maze and forced swim testing in F1 and F0.** (**a**) F1 Time in Arm: RM Two-way ANOVA: Stimulus Arm: $F_{(1,41)}$=87.6, P<.0001; Exposure $F_{(2,41)}$=100.0, *P*<.001; Interaction $F_{(6,41)}$=0.006, ns; Sidak's post-hoc tests, open vs closed for all groups, *P*<.001; Between group comparison, effect of exposure, ns; (**c**) F0 Time in Arm: Two-way ANOVA: Stimulus Arm: $F_{(1,52)}$=746, *P*<.0001; Exposure $F_{(2,52)}$=0, ns; Interaction $F_{(2,52)}$=0.32, ns; Sidak's post-hoc tests, open vs closed for all groups, *P*<.0001. Between group comparison, effect of exposure, ns; (**b**) F1 Total arm entries: One-way ANOVA: $F_{(2,41)}$=1.8, ns; (**d**) F0 Total arm entries: One-way ANOVA: $F_{(2,52}$=3.7, *P*<.05, Tukey's post-hoc test, VEH/CON vs H-DE-71, *P*<.05. (**e**) F1 time spent immobile: One-way ANOVA: $F_{(2,17)}$=0.4, ns. F0 time spent immobile: One-way ANOVA: $F_{(2,45)}$=0.8, ns. *P*<.05, Tukey's post-hoc test, VEH/CON vs H-DE-71, *P*<.05.

**Supplementary Figure 7. Open Field Testing in F0.** (**a**) F0 Distance Travelled Arena: RM Mixed Effects Two-way ANOVA: Exposure $F_{(2,26)}$=4.9, *P*<.05; Time $F_{(11,274)}$=13.9, *P*<.0001; Interaction $F_{(2,274)}$=2.4, *P*<.001; (**b**) F0 Velocity Arena: RM Mixed Effects Two-way ANOVA: Exposure $F_{(2,26)}$=5.0, *P*<.05; Time $F_{(11,274)}$=13.5, *P*<.0001; Interaction $F_{(2,274)}$=2.2, *P*<.0001; (**c**) F0 Distance Travelled Periphery: RM Mixed Effects Two-way ANOVA: Exposure $F_{(2,26)}$=3.9, *P*<.05; Time $F_{(11,277)}$=15.4, *P*<.0001; Interaction $F_{(22,277)}$=1.4, ns; (**d**) F0 Distance Travelled Center: RM Mixed Effects Two-way ANOVA: Exposure $F_{(2,26)}$=2.0, ns; Time $F_{(11,280)}$=3.7, *P*<.0001; Interaction $F_{(22,280)}$=1.6, *P*<.05; (**e**) F0 Cumulative Distance Travelled Center vs Periphery: VEH/CON paired t-test: $t_{(7)}$=11.7, *P*<.0001; L-DE-71 $t_{(9)}$=11.2, *P*<.0001; H-DE-71 $t_{(10)}$=14.8, *P*<.0001. (**f**) F1 Cumulative Fecal Boli: Two-way ANOVA: Exposure $F_{(2,78)}$=6.6, *P*<.01; Time $F_{(2,78)}$=21.5, *P*<.0001; Interaction $F_{(4,78)}$=0.7, ns.
