## Supplementary data 2 for "Persistent autism-relevant phenotype produced by *in utero* and lactational exposure of female mice to the commercial PBDE mixture, DE-71"

Orchid IDs: E.V.K.: 0000-0002-4691-6618; A.E.B.: 0000-0002-6057-3770; K.W.S.: 0000-0002-8945-4062; M.C.C.: 0000-0002-0189-4179

[1]Department of Molecular, Cell and Systems Biology, University of California, Riverside, CA 92521, USA
[2] Neuroscience Graduate Program, University of California, Riverside, CA, 92521, USA
[3] Duke University, Nicholas School of the Environment, Durham, NC 27710, USA
[4]Department of Psychology, Loma Linda University, Loma Linda CA 92350, USA
[5]Biotechnology Department, Pontifical Catholic University of Puerto Rico, Ponce, Puerto Rico 00717-9997 USA.
[6] HelmholtzZentrum Munchen, German National Research Centre for Environmental Health (GmbH), Molecular EXposomics (MEX), Ingolstaedter Landstrasse 1, Neuherberg, Munich, Germany
[7]TUM, Wissenschaftszentrum Weihenstephan für Ernährung, Landnutzung und Umwelt, Department für Biowissenschaftliche Grundlagen, Weihenstephaner Steig 23, 85350 Freising, Germany
[8]Neurological and Endocrine Toxicology Branch, Public Health and Integrated Toxicology Division, CPHEA/ORD, U.S. Environmental Protection Agency, Research Triangle Park, NC 27711 USA

***Corresponding author**:
Dr. Margarita C. Curras-Collazo, Ph.D
Professor of Neuroscience
Department Molecular, Cell and Systems Biology
University of California, Riverside
Riverside, CA 92521
951-827-3960


#### **Declarations**

##### **Funding**

We acknowledge funding from UCR Committee on Research (CoR) Grants to M.C.C.; UC MEXUS Awards to M.C.C., E.V.K., M.C.V.; NSF GRFP to M.C.V.; MARC U STAR Fellowship and NIH T34 (T34GM062756) to G.M.G.; UCR GRMP to E.V.K.; Sigma Xi Grant-in-Aid of Research award to E.V.K., K.M.R., M.E.D.; UCR Undergraduate Minigrant to E.V.K., K.M.R., A.E.B., V.C., G.L., B.M.V.; STEM-HSI Department of Education Award to E.V.K.; UCR Chancellor's Fellowship to J.M.K., APS IOSP Scholarship to L.M.A, APS STRIDE to A.E.B., and NIH R01 ES016099 to H.M.S.

**Conflicts of interests/Competing interests**

The authors report no conflicts of interests and have no competing interests to declare.

**Disclaimer:** J.M.K. is now a 2nd Lieutenant at the Uniformed Services University, Department of Defense. Her work was performed at the University of California, Riverside before becoming a military officer. However, we want to emphasize that the opinions and assertions expressed herein are those of the authors and do not necessarily reflect the official policy or position of the Uniformed Services University or the Department of Defense.

The research described in this article has been reviewed by the Center for Public Health and Environmental Assessment, U.S. Environmental Protection Agency (EPA) and approved for publication. Approval does not signify that the contents necessarily reflect the views and policies of the agency nor does the mention of trade names of commercial products constitute endorsement or recommendation for use.

**Availability of Data and Material**

Not applicable.

**Code Availability**

Not applicable.

**CRediT authorship contribution statement**

**Elena V. Kozlova:** Conceptualization, Data curation, Formal Analysis, Funding acquisition, Investigation, Methodology, Project administration, Software, Supervision, Validation, Visualization, Writing – original draft, Writing – review & editing. **Matthew C. Valdez**: Conceptualization, Data curation, Formal Analysis, Funding acquisition, Investigation, Methodology, Project administration, Software, Supervision, Validation. **Maximilian E. Denys:** Formal Analysis, Funding Acquisition Investigation, Software, Writing – original draft. **Anthony E. Bishay:** Formal Analysis, Funding Acquisition, Investigation, Writing – original draft. **Julia M. Krum**: Data curation, Funding Acquisition, Investigation, Methodology, Software, Visualization. **Kayhon M. Rabbani:** Formal Analysis, Funding Acquisition, Investigation, Software, Validation, Data curation. **Valeria Carrillo:** Investigation, Funding Acquisition, Data curation. **Gwen M. Gonzalez:** Funding acquisition, Investigation, Methodology, Validation. **Jasmin D. Tran:** Formal Analysis, Investigation, Funding acquisition. **Brigitte M. Vazquez:** Investigation, Funding Acquisition. **Gregory Lampel:** Investigation, Funding Acquisition. **Laura M. Anchondo:** Investigation, Software. **Syed A. Uddin:** Investigation, Software, Validation. **Nicole M. Huffman:** Investigation, Software, Validation. **Eduardo Monarrez:** Investigation, Data curation, Software, Validation. **Duraan S. Olomi:** Investigation, Data curation. **Bhuvaneswari D. Chinthirla:** Investigation. **Richard E. Hartman:** Resources, Software, Methodology, Validation, Writing – review & editing. **Prasada Rao S. Kodavanti:** Funding acquisition, Resources, Writing – review & editing. **Gladys Chompre:** Investigation. **Allison L. Phillips:** Formal Analysis, Investigation, Writing – review & editing. **Heather M. Stapleton**: Formal Analysis, Funding acquisition, Methodology, Resources, Supervision, Validation, Writing – review & editing. **Bernhard Henkelmann:** Investigation, Methodology, Validation, Writing - original draft. **Karl-Werner Schramm:** Methodology, Resources, Funding acquisition, Supervision, Writing – review & editing. **Margarita C. Curras-Collazo:** Conceptualization, Formal Analysis, Funding acquisition, Methodology, Project administration, Resources, Supervision, Validation, Visualization, Writing – original draft, Writing – review & editing

**Ethics approval**

Care and treatment of animals was performed in accordance with guidelines from and approved by the University of California, Riverside Institutional Animal Care and Use Committee (AUP #00170026 and 20200018).

**Consent to participate**

Not applicable.

**Consent for publication**

All authors reviewed and approved the final manuscript.

### Supplementary Information

#### Contents Page

Supplementary Table 1 – MIQE checklist ……………………………………………………………………………..5-13
Supplementary Table 2 – RT-qPCR Primers……………………………………………………………………….....14
Supplementary Table 3 – RT-qPCR Primers for Additional Reference Genes……………………………………………14
Supplementary Table 4 – Primer Optimization Parameters……………………………………………………………...15
Supplementary Table 5 – RNA quantity and quality ……………………………………………………………………16
Supplementary Data 1 – Splice variants and secondary structure analysis of amplicons and primers …………..………...17-33
Supplementary Data 2 – Amplification plot and Melting curve analysis (RT-qPCR) ……………………………......34-41
Supplementary Data 4 – qPCR primer efficiency ………………………………...……………………………..41-57

2 **Supplementary Table 1.** MIQE checklist for authors, reviewers and editors. E = essential information; D = desirable information.

| ITEM TO CHECK | IMPORTANCE | Description how item was addressed in study / article |
|---|---|---|
| **EXPERIMENTAL DESIGN** | | |
| Definition of experimental and control groups | E | Control group: Corn oil vehicle (VEH/CON); Experimental groups: 0.1 mg/kg/d DE-71 (L-DE-71) and 0.4 mg/kg/d DE-71 (H-DE-71) For details see materials and methods and Figure 1. RT-qPCR experiments were performed in F1 female offspring. |
| Number within each group | E | See supplementary statistical information for Figure 9. |
| Assay carried out by core lab or investigator's lab? | D | All experiments were carried out in the investigator's laboratory with the exception of use of thermocyclers which were located at the University of California Riverside Genomics Core Facility. |
| Acknowledgement of authors' contributions | D | M.C.V., M.C.C. and E.V.K. conceived the idea of the study/study design as well as designed/validated the primer pairs and probes used. E.V.K., M.C.C., B.M.V., K.M.R., V.C., L.A., M.E.D. conducted the experiments. E.V.K., M.C.C., V.C. and K.M.R. analysed the results. E.V.K and M.C.C. wrote the manuscript. E.V.K., A.E.B., K.M.R. created the figures, tables and the supplementary material. All authors reviewed the manuscript. |
| **SAMPLE** | | |
| Description | E | The Palkovitz Punch technique was used to allow for extraction of minute quantities of brain tissue from specific nuclei. Mouse brain tissue micropunches were obtained using brain cryosections (300 uM) and immediately placed in 1.5 mL autoclaved Eppendorf tubes containing 350 uL of Trizol reagent, homogenized and snap frozen over dry ice. |
| Volume/mass of sample processed | D | Brain tissue punches were obtained using custom-made micropunchers. Micropunchers consist of three components: Precision Glide Hypodermic Needles (BD Biosciences, USA) ; 16G x 1 in (25mm), Disposable Syringes with Luer-Lok Tips 3 mL, and T-pins. The inner diameter of the 16-gauge needle was 1.651 mm. One micropunch from a 300 µm section yielded a volume of ~.67 $mm^3$. The average density of brain tissue is 1.04 $g/cm^3$, yielding ~667 µg of tissue per punch. |
| Microdissection or macrodissection | E | Microdissection. Micropunch technique yields nuclei specific brain samples. |

| | | |
|---|---|---|
| Processing procedure | E | Tissue punches were obtained by a trained neuroscientist using a mouse brain atlas (Paxinos and Franklin, *The Mouse Brain in Stereotaxic Coordinates*, 5th Edition), immediately placed in Trizol, homogenized and snap frozen. |
| If frozen - how and how quickly? | E | Samples were frozen over a slurry of dry ice (78ºC) immediately after homogenization in Trizol reagent in Eppendorf tubes. |
| If fixed - with what, how quickly? | E | Not fixed. |
| Sample storage conditions and duration (especially for FFPE samples) | E | Samples were stored at -80ºC in cryoboxes until further use. |
| **NUCLEIC ACID EXTRACTION** | | |
| Procedure and/or instrumentation | E | Total RNA from brain tissue micropunches was extracted using the Qiagen RNeasy Micro or RNeasy Plus Micro Kits according to the manufacturer's instructions. The resulting RNA pellet was eluted in nuclease-free water (14µl) with immediate ice-cooling. The procedure enriches for mRNA since all RNA molecules longer than 200 nucleotides are purified. |
| Name of kit and details of any modifications | E | RNeasy Plus Micro Kit (74004), RNeasy Micro Kit (74034; Qiagen, USA). We modified the manufacturers RNA isolation protocol by substituting the supplied RLT lysis buffer, which we found was inadequate for lysis of brain tissue for a partial phenol-chloroform extraction. |
| Source of additional reagents used | D | QIAzol Lysis Reagent (79306; Qiagen, USA); Chloroform (C297-4; Fisher Chemical, USA), Ethanol (V1016; Koptec, USA), UltraPure Distilled Water DNase, RNAse free (10977-015; Invitrogen, USA) |
| Details of DNase or RNAse treatment | E | DNase treatment was performed with the RNase-Free DNase set supplied with the Qiagen RNeasy Micro kit. Each sample was treated with 27 Kunitz Units of DNAse I suspended in Buffer RDD while the RNA was suspended on the spin column membrane and allowed to sit for 15 min at RT. Samples processed with the RNeasy Plus Micro Kit were spun through the gDNA Eliminator columns which efficiently remove the majority of the genomic DNA without DNase treatment. No RNAse treatment was performed. |
| Contamination assessment (DNA or RNA) | E | In each experiment, a no-template-control (NTC) without mRNA and a -RT control (complete RNA synthesis reaction without the addition of the enzyme reverse transcriptase (RT)) were added for each primer pair or probe per region of interest(s) ROI(s) in order to detect primer-dimer formation, contamination or presence of genomic DNA (gDNA). |

| | | |
|---|---|---|
| Nucleic acid quantification | E | RNA concentration was determined by measuring the absorbance of UV light at 260 nm. The RNA conversion factor derived from Beer-Lambert's law states that a sample with $A_{260}$ of 1.0 will theoretically contain 40 µg/ml pure RNA. |
| Instrument and method | E | NanoDrop-2000 UV spectrophotometer (Thermo Fisher Scientific, Waltham, MA, USA) |
| Purity (A260/A280) | D | RNA purity was determined by measuring the absorbance ratio $OD_{260nm/280nm}$ as well as $OD_{260nm/230nm}$. An $OD_{260nm/280nm}$ ratio of >1.6 was considered protein-free RNA, and an $OD_{260nm/230nm}$ ratio of >1.0 phenol-/ethanol-free RNA (Supplementary Table 5). The increased A230 in some RNA samples was due to the high concentrations of guanidine thiocyanate present in the extraction reagent. Low 230 ratios were determined by the manufacturer to not compromise the reliability of downstream applications, nevertheless, we chose a cutoff of 1.0 for our samples. RNA quality was assessed with a spectrophotometer. The mean $OD_{260nm/230nm}$ and $OD_{260nm/280nm}$ ratios by ROI were as follows: **SON**$_{260nm/230nm}$ : 1.22±0.049; **SON**$_{260nm/280nm}$ : 1.77±0.049; **PVN**$_{260nm/230nm}$: 1.29±0.053; **PVN**$_{260nm/280nm}$: 1.76±0.022; **AMG**$_{260nm/230nm}$ : 1.16±0.053; **AMG 280**: 1.74±0.017; **LS**$_{260nm/230nm}$: 1.29±0.043; **LS**: 1.69±0.016$_{260nm/280nm}$; **BNST**$_{260nm/230nm}$: 1.31±0.065; **BNST**$_{260nm/280nm}$: 1.78±0.019 |
| Yield | D | RNA yield was measured on the spectrophotometer. The mean total RNA concentration ∓SEM by region of interest (ROI) was as follows: **SON**: 46.3±3.15 ng/uL, **PVN**: 77.5±5.39 ng/uL, **AMG**: 67.8±3.98 ng/uL, **LS**: 60.2±2.93 ng/uL , **BNST**: 88.7±5.76 ng/uL |
| RNA integrity method/instrument | E | RNA integrity was determined with an Agilent 2100 Bioanalyzer (Agilent Technologies Inc. Santa Clara, CA, USA) according to the manufacturer's protocol (Supplementary Table 5). |
| RIN/RQI or Cq of 3' and 5' transcripts | E | RIN values ranged from 6.9 to 8.7 (mean 7.6, SD 0.60), indicating high RNA integrity (Supplementary Table 5) |
| Electrophoresis traces | D | Electrophoresis traces were determined with an Agilent 2100 Bioanalyzer (Agilent Technologies Inc. Santa Clara, CA, USA) according to the manufacturer's protocol (Supplementary Table 5) |
| Inhibition testing (Cq dilutions, spike or other) | E | To evaluate primer efficiency and the absence of inhibitor a Log10 serial dilution series of a RNA sample was amplified for each gene of interest and reference gene candidate. The highest dilution at which all technical replicates were texted was set as the limit of detection (LOD). A standard curve was generated within the BioRad Maestro software using a linear regression of the Cq values and the dilution factor within the linear dynamic range (LDR). The coefficient of determination ($r^2$) and the reaction efficiency (E) were determined from the slope of the standard curve (E = (10-1/slope -1) x 100%). Primer pairs with a $r^2$ of >.996 were acceptable and efficiencies (E) with 90-110% were considered acceptable. (Supplementary Table 2 and 3 ) |

| | | |
|---|---|---|
| **REVERSE TRANSCRIPTION** | | |
| Complete reaction conditions | E | cDNA was synthesized using the Luna Universal One-Step RT-qPCR or Luna Universal One-Step Probe RT-qPCR kits (New England Biolabs, USA). During the cycling protocol, a 55°C RT step temperature was used for a single cycle of 10 min, which is the optimal temperature for the Luna WarmStart Reverse Transcriptase. |
| Amount of RNA and reaction volume | E | **Amount of RNA**: 4 ng, with the exception one lateral septum experiment which used 1 ng; **Reaction volume**: 20 µl |
| Priming oligonucleotide (if using GSP) and concentration | E | Full composition details are not provided by New England Biolabs (NEB) |
| Reverse transcriptase and concentration | E | Luna WarmStart Reverse Transcriptase (1 uL of Luna WarmStart RT Enzyme Mix was added/rxn). Full composition details are not provided by NEB. |
| Temperature and time | E | 10 min at 55°C; 1 cycle |
| Manufacturer of reagents and catalogue numbers | D | Specified in "Complete reaction conditions" |
| Cqs with → without RT | D | The signal of the amplification plot without reverse transcriptase was very late and there was a high Cq value difference between the -RT control and all samples. ***Avp*:** 18.24 → 36.22; ***Avp1ar*:** 23.18 → 43.06; ***Adcyap1*:** 20.04 → 36.72; ***Adcyap1r1*:** 20.32 → 37.65**;** ***Oxt*:** 17.67→ 37.34; ***Actb*:** 15.19 → 37.93; ***Oxtr* with hydrolysis probe:** 27.19 → not detected**;** ***Actb* with hydrolysis probe:** 18.44 → not detected; Additional candidate reference genes: ***GAPDH*:** 14.69 → 31.85; ***HPRT*:** 19.02 → 35.79 |

| | | |
|---|---|---|
| Storage conditions of cDNA | D | Not applicable |
| **qPCR TARGET INFORMATION** | | |
| If multiplex, efficiency and LOD of each assay. | E | ***Oxtr E***: 107.1%, LOD: 100 pg RNA equivalent (weak signal at 10 pg and 1 pg); ***Actb*** **E**: 101.7%, LOD: 10 pg (weak signal at 1 pg) |
| Sequence accession number | E | We design our primers using target gene nucleotide sequences from the NCBI Nucleotide database (GeneBank, http://www.ncbi.nlm.nih.gov/nuccore). *Actb*, *Oxt*, *Oxtr* primers were purchased as PrimeTime pre-designed assays from IDT. (Provided in Supplementary Table 2) |
| Location of amplicon | D | Provided in Supplementary Table 2 |
| Amplicon length | E | Provided in Supplementary Table 2 |
| In silico specificity screen (BLAST, etc) | E | Provided in Supplementary Data 1. No secondary structures present at annealing temperature were detected as determined in silico by UNAFold (http://eu.idtdna.com/UNAFold?, Suboptimality 50%; Integrated DNA Technologies Inc., Coralville, IA, USA). |
| Pseudogenes, retropseudogenes or other homologs? | D | Sequence alignment, possible splicing and targeted transcript variants as well as absence of targeted pseudogenes, retropseudogenes or other homologs were assessed upon primer design by NCBI PrimerBLAST (National Center for Biotechnology Information, Bethesda MD, USA, https://www.ncbi.nlm.nih.gov/tools/primer-blast) and PrimerCheck (SpliceCenter der Genomics and Sequence alignment D Bioinformatics Group, LMP, CCR, NCI, http://projects.insilico.us/SpliceCenter/PrimerCheck.jsp). |
| Sequence alignment | D | |
| Secondary structure analysis of amplicon | D | Provided in Supplementary Data 1. No secondary structures present at annealing temperature were detected as determined in silico by UNAFold (http://eu.idtdna.com/UNAFold?, Suboptimality 50%; Integrated DNA Technologies Inc., Coralville, IA, USA). |
| Location of each primer by exon or intron (if applicable) | E | Provided in Supplementary Table 2 |
| What splice variants are targeted? | E | Provided in Supplementary Table 2 |
| **qPCR OLIGONUCLEOTIDES** | | |

| | | |
|---|---|---|
| Primer sequences | E | Provided in Supplementary Table 2 |
| RTPrimerDB Identification Number | D | Not applicable, primers were constructed and validated by the authors or purchased as pre-designed assays from IDT. |
| Probe sequences | D | Provided in Supplementary Table 2 |
| Location and identity of any modifications | E | Primers received no terminal or other modifications. |
| Manufacturer of oligonucleotides | D | Primers were synthesized by Integrated DNA Technologies (Coralville, IA, USA). |
| Purification method | D | Primers were purified by standard desalting by Integrated DNA Technologies (Coralville, IA, USA). |
| **qPCR PROTOCOL** | | |
| Complete reaction conditions | E | For RT-qPCR reactions we used CFX Connect thermocycles (Bio-Rad, USA) with 96 well PCR plates (MLL9601, Bio-Rad Laboratories Inc., USA) in combination with Microseal 'B' seal Seals (MSB1001, Bio-Rad Laboratories, Inc., USA). Into each well, NEB Luna Universal or probe one-step master mix (New England Biolabs, USA) consisting of 10uL of Luna Universal One-Step Reaction Mix (2X), 1 uL of Luna WarmStart RT Enzyme Mix, 0.8 uL of forward and reverse primer (200-500nM final), 4uL of 2-4 ng of mRNA and 5-5.4 uL of nuclease-free H20 (10977-015; Invitrogen, USA) for a total of 20uL/rxn. Master-mixes of all components except mRNA were prepared to minimize technical errors during manual pipetting. Amplification reactions for genes of interest were performed in triplicate in 50 cycles (RT 55°C/10 min; initial denaturation 95°C/1 min; per cycle 95°C/10 s denaturation, 60°C or 55 °C/30 s extension; 65-95°C in 0.5°C, 5s increments melt curve analysis). Reactions were ran in triplicate. |
| Reaction volume and amount of cDNA/DNA | E | Reaction volume: 20 µl; Amount of mRNA: 4 uL of an 1000 pg/uL dilution of the mRNA stock solution which yields **4 ng/rxn**. With the exception of one lateral septum experiment which used 4 uL of a 250 pg/uL dilution which yields **1ng/rxn**. |
| Primer, (probe), Mg++ and dNTP concentrations | E | Full composition details are not provided by NEB. |
| Polymerase identity and concentration | E | Hot Start Taq. Full composition details are not provided by NEB. |

| | | |
|---|---|---|
| Buffer/kit identity and manufacturer | E | Luna Universal One-Step RT-qPCR or Luna Universal One-Step Probe RT-qPCR kits (E3005, E3006, New England Biolabs, USA) |
| Exact chemical constitution of the buffer | D | Full composition details are not provided by NEB. |
| Additives (SYBR Green I, DMSO, etc.) | E | The NEB kits contain a dsDNA intercalating dye that is functionally equivalent to SYBR-Green. Full composition details are not provided by NEB. |
| Manufacturer of plates/tubes and catalog number | D | 96 well PCR plates, clear (MLL9601, Bio-Rad Laboratories Inc., USA) in combination with Microseal 'B' seal Seals (MSB1001, Bio-Rad Laboratories, Inc., USA) |
| Complete thermocycling parameters | E | RT 55°C/10 min; initial denaturation 95°C/1 min; per cycle 95°C/10 s denaturation, 60°C or 55 °C/30 s extension; 65-95°C in 0.5°C, 5s increments melt curve analysis. See Supplementary Data 2 |
| Reaction setup (manual/robotic) | D | Manual |
| Manufacturer of qPCR instrument | E | CFX Connect Real-Time PCR Detection System (Bio-Rad Laboratories Inc., USA) |
| **qPCR VALIDATION** | | |
| Evidence of optimisation (from gradients) | D | Primer optimization is evidenced by melt curve analysis and agarose gel electrophoresis, qPCR efficiency. Melting temperatures $T_m$ of primers as validated by the IDT are provided in Table 2. |
| Specificity (gel, sequence, melt, or digest) | E | Specific amplification of target reference genes was assessed byhaving a single band of correct size during agarose gel electrophoreses and a specific peak in melting curve analysis (95°C for 15s, 60°C for 15s, then continuous temperature increase to 95°C and fluorescence measurement). For each primer pair and qPCR run we also tested a no-template-control (NTC) without cDNA and a -RT control (cDNA synthesis without enzyme reverse transcriptase added) on the same plate to exclude false positives caused by unspecific amplification such primer dimers, contamination or genomic DNA. (Supplementary Data 2). |

| | | |
|---|---|---|
| For SYBR Green I, Cq of the NTC | E | The signal of the amplification plot during efficiency analysis for standard curve generation was very late or not detected (ND) and there was a high Cq value difference between the negative control and all mRNA dilutions. ***Avp*:** ND**;** ***Avp1ar*:** ND**;** ***Adcyap1*:** 38.48; ***Adcyap1r1*:** 41.23; ***Oxt*:** ND; ***ActB*:** 39.77; ***Oxtr* with hydrolysis probe:** 39.04; ***ActB* with hydrolysis probe:** ND; **Additional candidate reference genes: *GAPDH*:** 37.91; ***HPRT*:** 36.05 (Supplementary Table 4) |
| Standard curves with slope and y-intercept | E | ***Avp*:** y = 38.097, slope: -3.371**;** ***Avp1ar*:** y=44.248, slope: -3.478; ***Adcyap1*:** y=39.564, slope: -3.279; ***Adcyap1r1*:** y=40.648, slope: -3.415; ***Oxt*:** y=38.441, slope: -3.400; ***Actb*:** y=35.122, slope: -3.331**;** ***Oxtr* with hydrolysis probe:** y=46.429, slope: -3.162**;** ***ActB* with hydrolysis probe:** y=38.090, slope: -3.282; **Additional candidate reference genes: *GAPDH*:** y=34.763, slope: -3.493; ***HPRT*:** y=38.421, slope: -3.256 (Supplementary Table 4) |
| PCR efficiency calculated from slope | E | Provided in Table 2. |
| Confidence interval for PCR efficiency or standard error | D | CIs of qPCR efficiencies (E) for genes analyzed in RT-qPCR experiments are provided in Supplementary Data 3. |
| $R^2$ of standard curve | E | ***Avp*:** 0.998**;** ***Avp1ar*:** 0.989**;** ***Adcyap1*:** 0.999; ***Adcyap1r1*:** 0.999; ***Oxt*:** 1.00; ***Actb*:** 1.00; ***Oxtr* with hydrolysis probe:** 1.00; ***Actb* with hydrolysis probe:** 0.999; **Additional candidate reference genes: *GAPDH*:** 0.999; ***HPRT*:** 0.997 |
| Linear dynamic range | E | The linear dynamic range (LDR) used a 10-fold dilution series of mRNA that ranged from 1:10-1:$10^6$. Standard curves were calculated with dilutions that fell within the LDR. (Supplementary Data 3) |
| Cq variation at LOD | E | Supplementary Data 3 |
| Confidence intervals throughout range | D | CIs of Cq were calculated for replicates in the serial dilution range. Supplementary Data 3. |
| Evidence for limit of detection | E | This was determined by a not detectable Cq value or standard error of >1 for one of more of the replicates at a dilution level. This criteria was used to set the LOD at the previous dilution level. Supplementary Data 3 |

| | | |
|---|---|---|
| If multiplex, efficiency and LOD of each assay. | E | Oxtr E: 107.1%, LOD: 100 pg RNA equivalent (weak signal at 10 pg and 1 pg); *β-Actin* E: 101.7%, LOD: 10 pg (weak signal at 1 pg) |
| **DATA ANALYSIS** | | |
| qPCR analysis program (source, version) | E | CFX Maestro software, version 1.1 (Bio-Rad, USA) |
| Cq method determination | E | Single Threshold Method (Baseline Subtracted Curve Fit, Fluorophore Analysis Mode, Drift Correction Off) |
| Outlier identification and disposition | E | Each biological replicate was ran in triplicate. Technical replicate Cq that had error of greater than 1 SD deviation of mean were excluded. |
| Results of NTCs | E | ***Avp*: ** 38.8; ***Avp1ar*:** 37.1; ***Adcyap1*:** 37.3; ***Adcyap1r1*:** 37.0; ***Oxt*:** 35.6; ***Actb*:**>40; ***Oxtr* with hydrolysis probe:** 37.3**;** ***ActB* with hydrolysis probe:** >40 |
| Justification of number and choice of reference genes | E | We have previously determined that DE-71 treatment does not affect reference gene stability for B-actin. For this study we ran two additional reference candidates and determined them to be unchanged by treatment as well. |
| Description of normalisation method | E | Samples were normalized according to the Pfaffl method. |
| Number and concordance of biological replicates | D | *n*=6-8 subjects/group |
| Number and stage (RT or qPCR) of technical replicates | E | qPCR reactions were performed in triplicates (technical replicates n = 3). |
| Repeatability (intra-assay variation) | E | Not available. |

| | | |
|---|---|---|
| Reproducibility (inter-assay variation, % CV) | D | Not available. |
| Power analysis | D | The number of biological replicates (n = 6-8 subjects/group) was based on previous studies and corresponds to the number of replicates generally used in brain punch RT-qPCR experiments. |
| Statistical methods for result significance | E | All biological samples were ran in triplicate and a mean and SD was determined. Technical outliers that were >1 SD from mean were excluded. Outliers were also identified from melt curve analysis as off target amplification or primer dimers. Cq values and efficiencies were used to generate Pfaffl ratios. Outliers were removed using the ROUT method (5%) in GraphPad prism. A one-way ANOVA was used to make across group comparisons. |
| Software (source, version) | E | Google Sheets (Google, USA)<br>GraphPad Prism v9.0 (La Jolla, California, USA) |
| Cq or raw data submission using RDML | D | Not provided. |

**Table 2. RT-qPCR Primers**

| Target Gene | Gene Symbol | GenBank Accession Number | Primer/Probe Sequence | Exon Location Fwd/Rv | E (%) | Tm (°C) Fwd/Rv | Product Size (bp) | Anneal Temp (°C) |
|---|---|---|---|---|---|---|---|---|
| Arginine vasopressin | *Avp* | NM_009732.2 | F: CTCAACACTACGCTCTCCGC<br>R: CAGCAGATGCTTGGTCCGA | 1/1-2 | 98 | 60.8/57.9 | 173 | 55 |
| Arginine vasopressin receptor 1A | *Avp1ar* | NM_16847.2 | F: GCTGGACACCTTTCTTCATCGTC<br>R: CTGTTCAAGGAAGCCAGTAACG | 1/2 | 89.1 | 61.7/59.5 | 115 | 55 |
| Adenylate cyclase activating polypeptide 1 | *Adcyap1* | NM_009625.3 | F: AGGTGCTGGTGTTGGAATGAATGC<br>R: AATGCATGAGGGCAAGGGTAGGAA | 5 | 95 | 60.2/60.7 | 176 | 55 |
| Adenylate cyclase activating polypeptide 1 receptor 1 | *PAC1r* | NM_007407.4 | F: TTCACTACTGCGTGGTGTCCAACT<br>R: ATATCCCAGCATCCCGCATCATCA | 10/11-12 | 96.3 | 60.3/60.3 | 199 | 55 |
| Oxytocin | *Oxt* | NM_011025.4 | F: CCGAAGCAGCGTCTTTT<br>R: CTTGGCTTACTGGCTCTGAC | 1/2 | 96.9 | 55.7/55.5 | 131 | 60 |
| Oxytocin receptor | *Oxtr* | NM_001081147.2 | F: CGCACAGTGAAGATGACCTT<br>R: ATGGCAATGATGAAGGCAGA<br>P: 6-FAM-CTTCGTGCA-ZEN-GATGTGGAGCGTTCT-IBFQ | 1/2 | 107.1 | NA | 131 | 60 |
| Beta Actin | *β-Actin* | NM_007393.5 | F: GATTACTGCTCTGGCTCCTAG<br>R: GACTCATCGTACTCCTGCTTG<br>P: HEX-CTGGCCTCA-ZEN-CTGTCCACCTTCC-IBFQ | 5/6 | 99.6<br>101.7 | 55.0/54.4<br>NA | 147 | 60 |

**Table 3. RT-qPCR Primers for Additional Reference Candidate Genes**

| Target Gene | Gene Symbol | GenBank Accession Number | Primer/Probe Sequence | Exon Location | E (%) | Tm (°C) Fwd/Rv | Product Size (bp) | Anneal Temp (°C) | $r^2$ | Cq Range |
|---|---|---|---|---|---|---|---|---|---|---|
| **Glyceraldehyde 3-phosphate dehydrogenase** | **Gapdh** | **NM_001289726.1** | **F: TTGTGATGGGTGTGAACCACGAGA<br>R: GAGCCCTTCCACAATGCCAAAGTT** | **5** | **93.3** | **60.4/60.2** | **131** | **55** | **0.999** | **21.983 - 22.667** |
| **Hypoxanthine-guanine phosphoribosyltransferase** | **Hprt** | **NM_013556.2** | **F: TACGAGGAGTCCTGTTGATGTTGC<br>R: GGGACGCAGCAACTGACATTTCTA** | **9** | **102.8** | **58.7/59.4** | **138** | **55** | **.997** | **22.813 - 23.450** |

**Table 4. Primer Optimization Parameters**

| **Target Gene** | **Gene Symbol** | **E (%)** | **$r^2$** | **Y-int** | Slope | **Cq NTC** | **Cq NRT** |
|---|---|---|---|---|---|---|---|
| Arginine vasopressin | *Avp* | 98 | 0.998 | 38.097 | -3.371 | 0.00 | 36.22 |
| Arginine vasopressin receptor 1A | *Avp1ar* | 89.1 | 0.996 | 44.824 | -3.615 | 0.00 | 43.06 |
| Oxytocin | *Oxt* | 96.9 | 1.00 | 38.441 | -3.400 | 0.00 | 37.34 |
| Adenylate cyclase activating polypeptide 1 | *Adcyap1* | 95 | 0.999 | 39.564 | -3.279 | 38.48 | 36.72 |
| Adenylate cyclase activating polypeptide 1 receptor 1 | *Adcyap1r1* | 96.3 | 0.999 | 40.648 | -3.415 | 41.23 | 37.65 |
| Oxytocin receptor | *Oxtr* | 107.1 | 0.997 | 46.429 | -3.162 | 39.04 | 0.00 |

| | | | | | | | |
|---|---|---|---|---|---|---|---|
| Beta Actin | *β-Actin* | 106.8<br>101.7 | 1.000<br>0.999 | 35.122<br>38.090 | -3.331<br>-3.282 | 39.77<br>0.00 | 37.93<br>0.00 |

**Table 5. Nanodrop and Bioanalyzer Results**

| ROI | Treatment | RNA Purity 260/280 | RNA Purity 260/230 | Yield (ng/uL) | RNA Integrity (RIN) | rRNA ratio [28s/18s] |
|---|---|---|---|---|---|---|
| BNST | 0.1 mg/kg | 1.82 | 1.52 | 64 | 6.90 | 1.2 |
| BNST | 0.4 mg/kg | 1.82 | 1.5 | 73 | 7.50 | 1.7 |
| BNST | VEH/CON | 1.74 | 1.53 | 35 | 7.30 | 0.9 |
| SON | VEH/CON | 1.73 | 1.54 | 29 | 6.90 | 0.8 |
| SON | 0.1 mg/kg | 1.89 | 1.73 | 74 | 7.10 | 1.1 |
| LS | VEH/CON | 1.81 | 1.64 | 43 | 8.40 | 1.2 |
| LS | 0.1 mg/kg | 1.71 | 1.74 | 41 | 8.30 | 1.7 |
| LS | 0.4 mg/kg | 1.67 | 1.49 | 29 | 8.70 | 1.6 |
| PVN | VEH/CON | 1.75 | 1.22 | 53 | 7.30 | 2.0 |
| PVN | 0.4 mg/kg | 1.85 | 1.75 | 61 | 7.80 | 2.3 |
| AMG | VEH/CON | 1.56 | 1.47 | 17 | 7.70 | 0.7 |
| AMG | 0.1 mg/kg | 2.00 | 1.63 | 119 | 7.30 | 1.1 |

**Supplementary Data 1. Splice variants and secondary structure analysis of amplicons and primers of the genes of interest and reference genes.**

***Avp*** PrimerBLAST (National Center for Biotechnology Information, Bethesda MD, USA, https://www.ncbi.nlm.nih.gov/tools/primer-blast )

**Primer pair 1**

| | Sequence (5'->3') | Length | Tm | GC% | Self complementarity | Self 3' complementarity |
|---|---|---|---|---|---|---|
| Forward primer | CTCAACACTACGCTCTCCGC | 20 | 60.80 | 60.00 | 2.00 | 2.00 |
| Reverse primer | CAGCAGATGCTTGGTCCGA | 19 | 60.08 | 57.89 | 6.00 | 3.00 |

**Products on target templates**

>NM_009732.2 Mus musculus arginine vasopressin (Avp), mRNA

```
product length = 173
Forward primer  1    CTCAACACTACGCTCTCCGC  20
Template        63   ....................  82

Reverse primer  1    CAGCAGATGCTTGGTCCGA  19
Template        235  ...................  217
```

***Avp*** PrimerCheck (SpliceCenter der Genomics and Bioinformatics Group, LMP, CCR, NCI, http://projects.insilico.us/SpliceCenter/PrimerCheck.jsp )

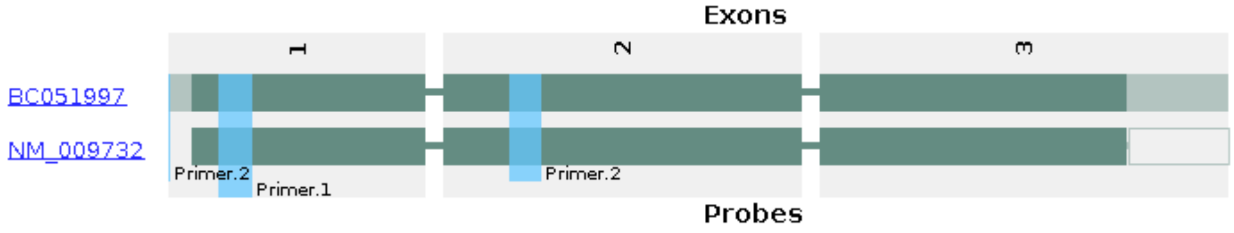

Avp

Avp Amplicon Sequence (https://www.ncbi.nlm.nih.gov/nucleotide/)

**<u>CTCAACACTACGCTCTCCGC</u>**TTGTTTCCTGAGCCTGCTGGCCTTCTCCTCCGCCTGCTACTTCCAGAACTGCCCAAGA
GGCGGCAAGAGGGCCATCTCTGACATGGAGCTGAGACAGGTACCACTGTGGTCCCTTTAGGGCTGCTGGC
AGTGCCTGTAGGGACGGGTCAGGGGCTAGGAGAGAGGGAAATGTTATCTGAGCAGTCAGACTTTATGGGA
GGTTCCTGGAAGGAGGCAGTATCTTACAGCAGAGTAGATGGACTACCCAGAAGGGTGAGAGGGGACCAGG
TGCTAGAGAAGCCGCATAAAGGATACTGTCCCCAGGCAGGGGATATGCCAGAAAATGAGAGACACTTCCT

TATGACTGGGCTTGGGATGAGAACAGGTTAAACTGGGTGCCCTGGACTCCTCTGCACACCCGGAGGTTGA
GGACTGGGCAGATTATACAAAACTATTCTTGCCTGAATTCAAATCCTTTCCCACCCAGCTCAGCCTCCCT
TGGTGCCTTTTCTAGCCAGCAGTGCCAGCTTCTTCCTGTCCACAGAAGGTGGCCAATGCCCCATGCCCAA
GTGGAGCATTTCCCGCATCGAACCTCAGCCTCTTGCTCAGATCTGTTGTATTGTATGTTCAGCTGTGAGT
CTGCCTGCCCCTCTGGCAGAGTTTGAGGGAATCTAGCTACTAGGCTCAATTCTGGTCAGGCCATGGGTGG
CTCAATTTTGAGTTGTTGAACAAGTTCGAGTGGGCAGGTAGGCAGCTCCTGTAGTCTGCCTTCCCTTTGC
AGAGTTCCTTTGGAGGTGTGTCCGGGCACCTAATTTGGTCCTTGCCACCTACCAACTAAGACATAATAGG
TTGGCGGGAGGTAAAGGCTCATATGAAGCCCACCAGCGTGGGGCAGAGGTAAGAGCAAAGCCAGAAAACG
AGTGAGCTATCTAGATGCTCTGTGGGGAGTGAGAATCTAGGGATGTGTAGGAGGACCATCTGAATGACGG
AGAGGTAAGCCTCCGAGAGATGGCTGCACACCAGTGACACTGAGAACTGAGGAAGGTCTCCCTCAAGTGT
TGCCCCGCAGCGAGAGGGTTTTGAGACCTCATGAGCTGACCACTGATCTTTCTGATGACCCAGCCGGTTA
GATTTTCACTCTTGCCCTTACCGCTGCTTCGTCCTGGACATCGCCAGAGCACCAGCAACGCAAAGCAGCA
GGTGACACTAGGTTCCCACCGCCCCTCTTGGCCTCGTTCCAGCTGACCTCCCCCACCCCTTTCCTCCACA
GTGTCTCCCCTGCGGCCCGGGCGGCAAAGGACGCTGCT**TCGGACCAAGCATCTGCTG**

***Avp*** UNAFold (Integrated DNA Technologies Inc., Coralville, IA, USA, http://eu.idtdna.com/UNAFold? , Suboptimality 50%)

#### Structures

| Structure Name | Image | ΔG (kcal.mole$^{-1}$... | $T_M$ (°C) | ΔH (kcal.mole$^{-1}$) | ΔS (cal.K$^{-1}$mole$^{-1}$) | Output |
|---|---|---|---|---|---|---|
| 1 | 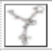 | -29.55 | 46.5 | -438.8 | -1372.64 | Ct Det |
| 2 | 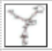 | -27.98 | 45.7 | -431.6 | -1353.76 | Ct Det |
| 3 | 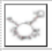 | -27.61 | 49 | -370.3 | -1149.4 | Ct Det |
| 4 | 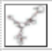 | -27.12 | 44.8 | -435.8 | -1370.71 | Ct Det |
| 5 | 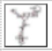 | -26.98 | 44.3 | -444.6 | -1400.7 | Ct Det |
| 6 | 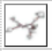 | -26.83 | 44.6 | -434.3 | -1366.67 | Ct Det |
| 7 | 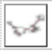 | -26.61 | 47 | -386.6 | -1207.4 | Ct Det |

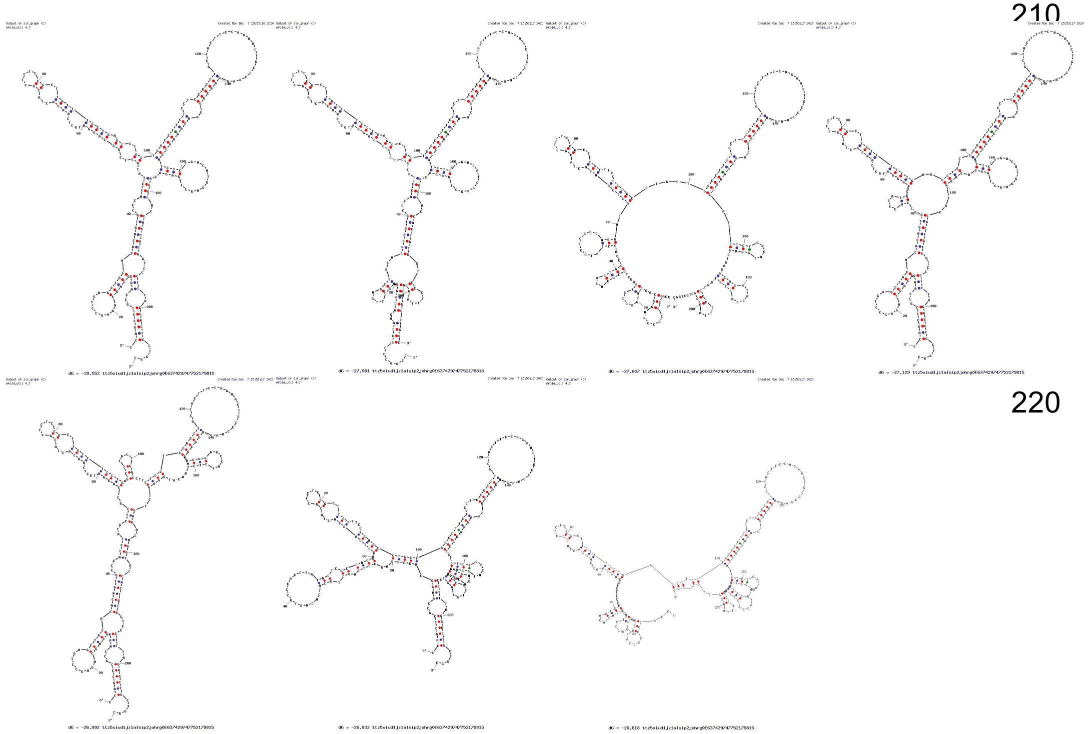

**Avp1ar** PrimerBLAST (National Center for Biotechnology Information, Bethesda MD, USA, https://www.ncbi.nlm.nih.gov/tools/primer-blast )

**Primer pair 1**

| | Sequence (5'->3') | Length | Tm | GC% | Self complementarity | Self 3' complementarity |
|---|---|---|---|---|---|---|
| **Forward primer** | GCTGGACACCTTTCTTCATCGTC | 23 | 61.71 | 52.17 | 3.00 | 1.00 |
| **Reverse primer** | CTGTTCAAGGAAGCCAGTAACG | 22 | 59.52 | 50.00 | 3.00 | 2.00 |

**Products on target templates**

>NM_016847.2 Mus musculus arginine vasopressin receptor 1A (Avpr1a), mRNA

```
product length = 115
Forward primer  1     GCTGGACACCTTTCTTCATCGTC  23
Template        1229  .......................  1251

Reverse primer  1     CTGTTCAAGGAAGCCAGTAACG  22
Template        1343  ......................  1322
```

**Avp1ar** PrimerCheck (SpliceCenter der Genomics and Bioinformatics Group, LMP, CCR, NCI, http://projects.insilico.us/SpliceCenter/PrimerCheck.jsp )

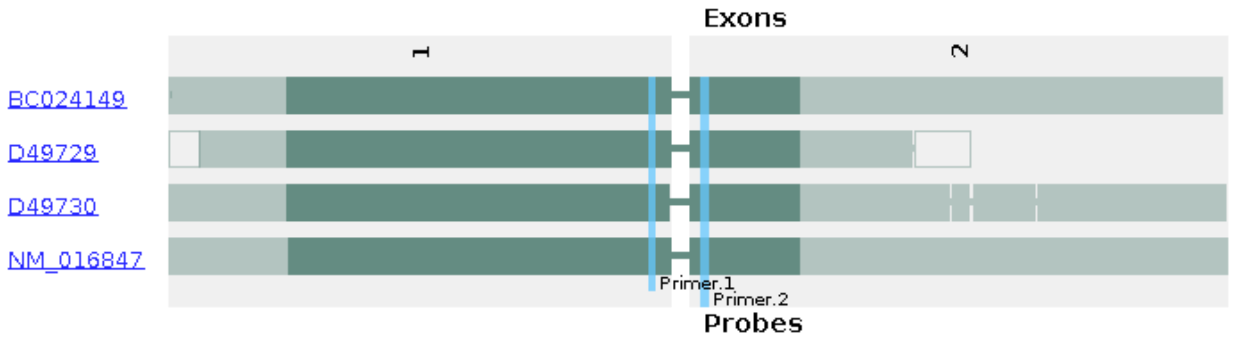

Avprla

Avp 1ar  Amplicon Sequence (https://www.ncbi.nlm.nih.gov/nucleotide/)

<u>**GCTGGACACCTTTCTTCATCGTC**</u>CAGATGTGGTCAGTCTGGGATACCAATTTCGTTTGGACCGGTATGTACTGAGACTACTATGGAAGAGGTGTGTGTGTGTGT
GTGTGTGTGTGTGTGTGCCTGTGTGTTTGTGGTGTGTGTTGGGGGGGGAGTGTTTGTGGTGTGTGTGTGTATGTATGTTATTTTTCATACAACCTATTCAATAGTTA
AGCTTCTATGTAAGAAGTCTTAAATGAACAAAATTGCAAATTCCTGGGACCTCCACCCCAGGTACACTGGAGCCCAGGATTCAGCATCTGCAGCATGTGTGACCA
CCCTAAGGCCACCAGCTGTGCCTAGAGTCACTAGTTACAAAACACTTGTGACATTTTGCCAAATTACTTAGGGTTCTTGGATATCCTGCCTTCCCATAGAGGCTCT
GGGGAATATTCTTCATGTCTTGGCTACTTTTTGTTGGATTAAGGAAACTATCACAATCAACAGAAAACTTCTCATGAGATTCTTCCAAGACCAAGGTTAGAGATAG
ACACACTGTTCTGAAACAAACAGAACGGATTTAAGAAGTTAGAAATTCCAAATTTGCATTTAGTTTGTGCAACCCATCTTCATTATAACGTTTTCATTTTTCCAACT
GTAATAACTTTCTTTTCCTGGCATCCAAATCAGAAGTCTTGATCAGCTAAATTCACTGGACTTCCCGCGGGAAGTTAACAACAAAACTATTCACGCCACACAAATC
CCTCTGTAGTTAATTGTGAAAGTTCCCCATCCCAACTGGCTGGTTTTATTTCAGTTCATTCACAAAACAGAAAAATCCATTTCCCAGGAGTTTATTGTTTATTTATT
TTGATCTTGTAATATATCTTCTTTTAATTAGGAAAAAAAAAAAAAAGACTTTCCAAACCAAGTATGTTACAGAAGATTTCCTGATGAAAACTCTAATCAGGATATAA
ATATCTAGGAGAAGCTATACCTTTCTGTAATTTTAAGATTGGGGTTTGGAATATATGCAATGCATTTAAATATATTATTTTGAATGTTCATAGATTTCGTTTAAGCA
AATACTTTTAGAGGCTGCATTGCTCATAGTAGTAAATAAATGCAAAGTCCTTTAGTCAATGCCAATCACATTTTAAAATTCTGTCCATTGCTTTATTGTGATTTAGA
CACAGTTAGTAGAATTTTCCCTTGCACTACACATGTTGAAGATGCCTGTTATTTTTAATATTGTGCGTGGCTTATGGTCACCTAGCACAGAGCAACGAATGGCTTG
CACCAGTGACTCTCAACCTTCTCAGTCCTGCCACACTCAACACAGTTCCTCGTGCTGTGATGACCCCCAACCATGAAATTGGCTTTGTCACTACTTCGTAACTGTA
ACTTTGCTACGGCTTCCAATCAGAATGTGAGTATCTGATATGCAGCCGGTCTCAGGTGACTTCGGTGACAAGGCTGTTCATGGTGAGCCACAGGTTGAGAACCAC
TGGTCTACAGTCTCTTTTGCATGACAAAAGTGGGGTGTTAGGAGTCATATACCAATCAGAAGACAGTTGATGTTTGATGGTTGTTCTTATTTTTTAATAACAGTTC
AAATAGTTTCACCATGAAAACTTTGATAGCCATGTAGCTTACTAGGGGTCATTCACTGAATAGTTCAAAATCCTGTTGGACTCCCTAACTCAGCTACTTTGGATTT
TTCTTTTTTTGTCCGTTTTAATTTTTGTTTTGTTCCTGTTGTTTGAAATGGGGCCATATACAACCCAGATGTTTTTCTTATTATAATTTAATTGCTTGAACTACAATCA
AGGACAGTCTTTTAAAATCAAACCAAATGTCTAATGATTCACAAAAATTCAATTGAAGCTGGGATTCTATTTTAAATACTCTCTTCCATACGGTTTTCATTGATATC
TGTTGAACAGCCCATATGGCATTGCCCAAGTTACTAGGATGAATACAATTCAGTTTATGTGATTATAAACACTTCAAATGTCCTTCTGTTAATACCATGTCTTTGTG
AATACCTTCACAAAGGCTCCTGCCACTGTATAGTATACAGTTTTAAAATAAGTTATTCAGTATCTTCATCTTCTTAGACTTTATATCCATTGTATTGAAGAGTTCTA
ACAGTGACTGGTTTAAACTATTTGAACTAATCTTAAGACATTTTCTTTTACAACCAGTCTCCAAAAAGTACTCATTTCCCTGAAACACCATTTCCCTAACAAATAA
GATGTCCATTTTTATCACTAATATGGAGCTTTTTGACTTGTTAGTCGGTAGGGCATAGTGTTCGGGGTCTCCCTCTGGCTGTTGATATGATCTCTCTCCTCGTTTCCC
ACACAGATTCCGAAAACCCTTCCACCACGATCACGG<u>**CGTTACTGGCTTCCTTGAACAG**</u>

#### Structures

| Structure Name | Image | ΔG (kcal.mole⁻¹... | $T_M$ (°C) | ΔH (kcal.mole⁻¹) | ΔS (cal.K⁻¹mole⁻¹) | Output |
|---|---|---|---|---|---|---|
| 1 | | -30.64 | 51.1 | -381 | -1175.11 | Ct Det |
| 2 | | -29.15 | 51.8 | -352.9 | -1085.88 | Ct Det |
| 3 | | -29.11 | 43.9 | -489.3 | -1543.49 | Ct Det |
| 4 | | -28.01 | 52.4 | -332.5 | -1021.26 | Ct Det |
| 5 | | -27.95 | 47.6 | -396.1 | -1234.78 | Ct Det |

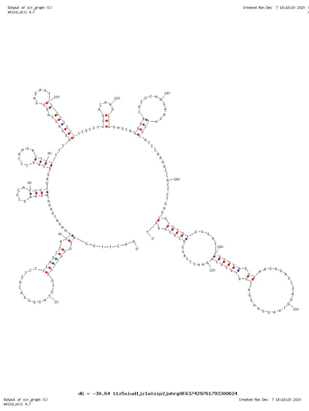
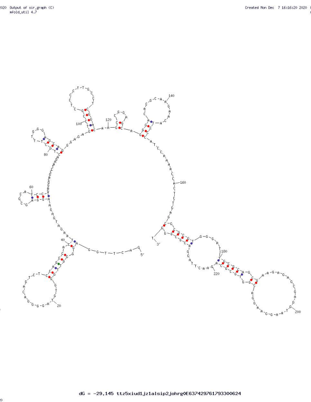
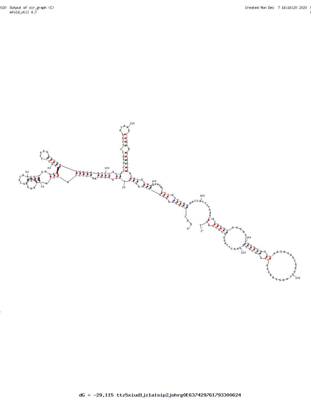
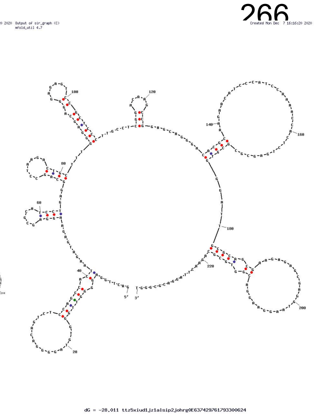

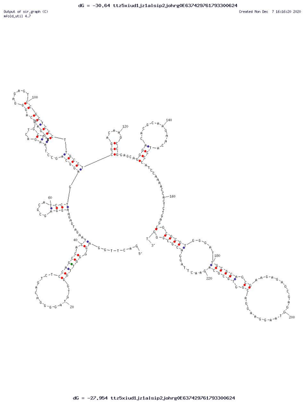

**Oxt** PrimerBLAST (National Center for Biotechnology Information, Bethesda MD, USA, https://www.ncbi.nlm.nih.gov/tools/primer-blast )

**Primer pair 1**

| | Sequence (5'->3') | Length | Tm | GC% | Self complementarity | Self 3' complementarity |
|---|---|---|---|---|---|---|
| **Forward primer** | CCGAAGCAGCGTCCTT | 16 | 56.68 | 62.50 | 5.00 | 4.00 |
| **Reverse primer** | CTTGGCTTACTGGCTCTGAC | 20 | 58.27 | 55.00 | 3.00 | 1.00 |

**Products on target templates**

>XM_006498910.5 PREDICTED: Mus musculus oxytocin (Oxt), transcript variant X1, mRNA

```
product length = 131
Forward primer  1     CCGAAGCAGCGTCCTT  16
Template        1859  ................  1844

Reverse primer  1     CTTGGCTTACTGGCTCTGAC  20
Template        1729  ....................  1748
```

>NM_011025.4 Mus musculus oxytocin (Oxt), mRNA

```
product length = 131
Forward primer  1    CCGAAGCAGCGTCCTT  16
Template        202  ................  187

Reverse primer  1   CTTGGCTTACTGGCTCTGAC  20
Template        72  ....................  91
```

**Oxt** PrimerCheck (SpliceCenter der Genomics and Bioinformatics Group, LMP, CCR, NCI, http://projects.insilico.us/SpliceCenter/PrimerCheck.jsp )

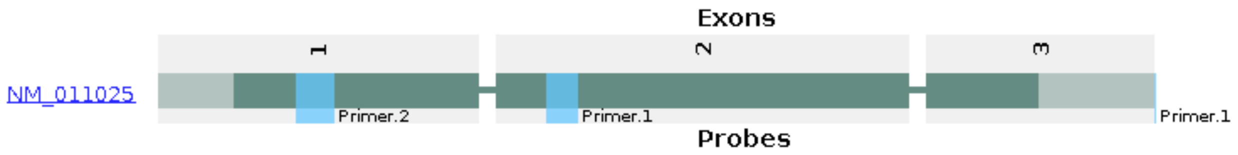

Oxt

**Oxt** Amplicon Sequence (https://www.ncbi.nlm.nih.gov/nucleotide/)

5'-
**CTTGGCTTACTGGCTCTGAC**CTCGGCCTGCTACATCCAGAACTGCCCCCTGGGCGGCAAGAGGGCTGTGCTGGACCTGGATATGCGCAAGGTGAGTCTCCCCGA
CCCTGTCCCTTCCCTTCCCGTTCTGGCGATGCTAAGGACCAGAGAAGCTCTCCCACCTACAGAGAGCATTCCCGCACACTTGCCAGCCCTACCAAGGCCTCGCGTG
GGAACCCAGGGCTTTGGGAAGTGTTAGGCTCCCTCTTGACGCCGTGAAGGTAACGACAATGCCGGAGCACCCACTGCCCCTCGCTCTGCCACAGTCCGGATTCGG
ATTGTGCACGGCGCCCACCCGCATCCTTCCCCACAGTGTCTCCCCTGCGGCCCGGGCGGC**AAAGGACGCTGCTTCGG-3'**

#### Structures

| Structure Name | Image | ΔG (kcal.mole$^{-1}$... | $T_M$ (°C) | ΔH (kcal.mole$^{-1}$) | ΔS (cal.K$^{-1}$mole$^{-1}$) | Output |
|---|---|---|---|---|---|---|
| 1 | | -13.66 | 41.1 | -266 | -846.36 | Ct Det |
| 2 | | -12.63 | 41.7 | -237.6 | -754.56 | Ct Det |
| 3 | | -12.16 | 37.5 | -303.2 | -976.16 | Ct Det |
| 4 | | -12.15 | 37.1 | -312.4 | -1007.03 | Ct Det |

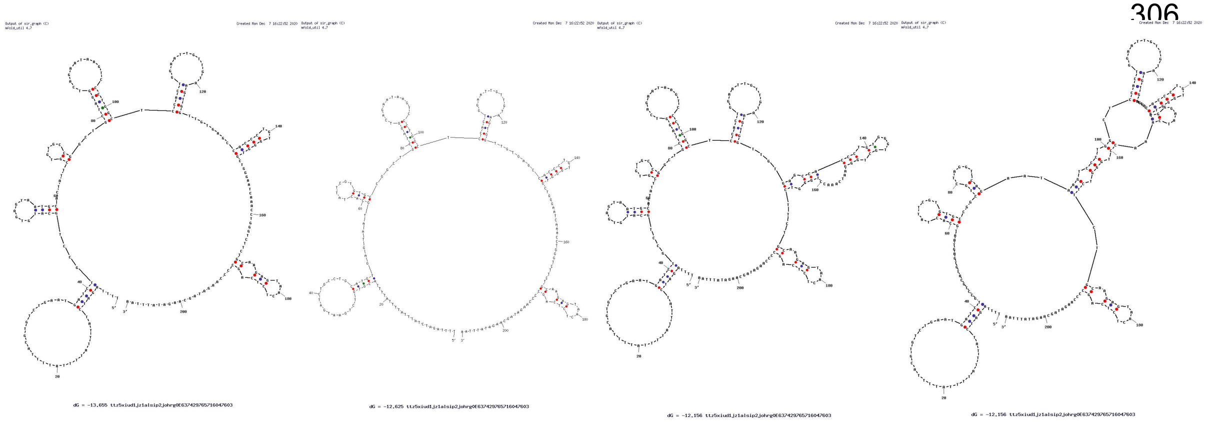
dG = -13.655 ttz5xiud1jz1alsip2johrg0E637429765716047603
dG = -12.625 ttz5xiud1jz1alsip2johrg0E637429765716047603
dG = -12.156 ttz5xiud1jz1alsip2johrg0E637429765716047603
dG = -12.156 ttz5xiud1jz1alsip2johrg0E637429765716047603

**Adcyap1** PrimerBLAST (National Center for Biotechnology Information, Bethesda MD, USA, https://www.ncbi.nlm.nih.gov/tools/primer-blast )

##### Primer pair 1

| | Sequence (5'->3') | Length | Tm | GC% | Self complementarity | Self 3' complementarity |
|---|---|---|---|---|---|---|
| **Forward primer** | AGGTGCTGGTGTTGGAATGAATGC | 24 | 64.16 | 50.00 | 2.00 | 2.00 |
| **Reverse primer** | AATGCATGAGGGCAAGGGTAGGAA | 24 | 64.27 | 50.00 | 6.00 | 0.00 |

**Products on target templates**

>XM_030249450.1 PREDICTED: Mus musculus adenylate cyclase activating polypeptide 1 (Adcyap1), transcript variant X1, mRNA

```
product length = 176
Forward primer  1     AGGTGCTGGTGTTGGAATGAATGC  24
Template        2918  ........................  2941

Reverse primer  1     AATGCATGAGGGCAAGGGTAGGAA  24
Template        3093  ........................  3070
```

>NM_001315503.1 Mus musculus adenylate cyclase activating polypeptide 1 (Adcyap1), transcript variant 2, mRNA

```
product length = 176
Forward primer  1     AGGTGCTGGTGTTGGAATGAATGC  24
Template        1665  ........................  1688

Reverse primer  1     AATGCATGAGGGCAAGGGTAGGAA  24
Template        1840  ........................  1817
```

>NM_009625.3 Mus musculus adenylate cyclase activating polypeptide 1 (Adcyap1), transcript variant 1, mRNA

```
product length = 176
Forward primer  1     AGGTGCTGGTGTTGGAATGAATGC  24
Template        1938  ........................  1961

Reverse primer  1     AATGCATGAGGGCAAGGGTAGGAA  24
Template        2113  ........................  2090
```

**Adcyap1** PrimerCheck (SpliceCenter der Genomics and Bioinformatics Group, LMP, CCR, NCI, http://projects.insilico.us/SpliceCenter/PrimerCheck.jsp )

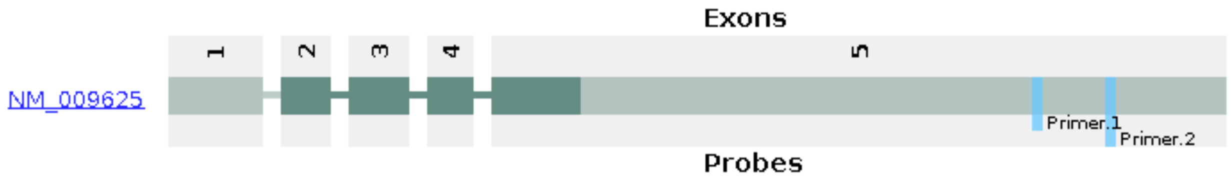

Adcyap1

**Adcyap1** Amplicon Sequence (https://www.ncbi.nlm.nih.gov/nucleotide/)

**AGGTGCTGGTGTTGGAATGAATGC**AAAGTACAATGTGTTTTTCTCCAGTGCTGTTCATGCTTTTCATGTTGTGAAATGGCCAGGATCCTCCCCTTTGAACACTGT
TCTGCAGAAGCCAGCTCTGTTCTTTGTGGATTTTCTGGAGACCCTCC**TTCCTACCCTTGCCCTCATGCATT**

378

#### Structures

| Structure Name | Image | ΔG (kcal.mole$^{-1}$... | $T_M$ (°C) | ΔH (kcal.mole$^{-1}$) | ΔS (cal.K$^{-1}$mole$^{-1}$) | Output |
|---|---|---|---|---|---|---|
| 1 | | -17.03 | 47.3 | -245 | -764.61 | Ct Det |
| 2 | | -16.63 | 45.8 | -255.2 | -800.17 | Ct Det |
| 3 | | -16.59 | 45.9 | -253 | -792.93 | Ct Det |
| 4 | | -16.38 | 45.5 | -254 | -797 | Ct Det |
| 5 | | -15.99 | 47.5 | -228.1 | -711.41 | Ct Det |
| 6 | | -15.94 | 46.4 | -237.6 | -743.47 | Ct Det |
| 7 | | -15.9 | 43.2 | -275.9 | -872.05 | Ct Det |
| 8 | | -15.56 | 44.7 | -250.5 | -787.99 | Ct Det |

413

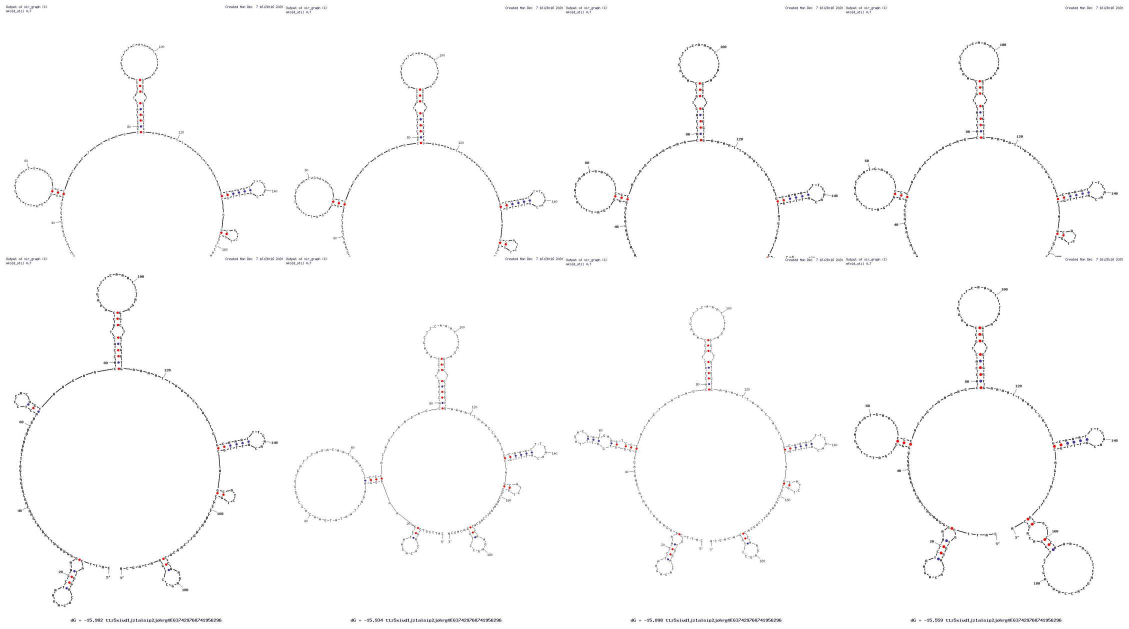

dG = -15.992 ttz5xiud1jz1alsip2johrg0E637429768741956296
dG = -15.934 ttz5xiud1jz1alsip2johrg0E637429768741956296
dG = -15.898 ttz5xiud1jz1alsip2johrg0E637429768741956296
dG = -15.559 ttz5xiud1jz1alsip2johrg0E637429768741956296

***Adcyap1r1*** PrimerBLAST (National Center for Biotechnology Information, Bethesda MD, USA, https://www.ncbi.nlm.nih.gov/tools/primer-blast )

#### Primer pair 1

| | Sequence (5'->3') | Length | Tm | GC% | Self complementarity | Self 3' complementarity |
|---|---|---|---|---|---|---|
| **Forward primer** | TTCACTACTGCGTGGTGTCCAACT | 24 | 64.28 | 50.00 | 6.00 | 1.00 |
| **Reverse primer** | ATATCCCAGCATCCCGCATCATCA | 24 | 63.99 | 50.00 | 4.00 | 2.00 |

**Products on target templates**

>XM_030255113.2 PREDICTED: Mus musculus adenylate cyclase activating polypeptide 1 receptor 1 (Adcyap1r1), transcript variant X7, mRNA

```
product length = 199
Forward primer  1     TTCACTACTGCGTGGTGTCCAACT  24
Template        1613  ........................  1636

Reverse primer  1     ATATCCCAGCATCCCGCATCATCA  24
Template        1811  ........................  1788
```

>XM_036165754.1 PREDICTED: Mus musculus adenylate cyclase activating polypeptide 1 receptor 1 (Adcyap1r1), transcript variant X5, mRNA

```
product length = 199
Forward primer  1     TTCACTACTGCGTGGTGTCCAACT  24
Template        1380  ........................  1403

Reverse primer  1     ATATCCCAGCATCCCGCATCATCA  24
Template        1578  ........................  1555
```

>XM_030255112.2 PREDICTED: Mus musculus adenylate cyclase activating polypeptide 1 receptor 1 (Adcyap1r1), transcript variant X4, mRNA

```
product length = 199
Forward primer  1     TTCACTACTGCGTGGTGTCCAACT  24
Template        1616  ........................  1639

Reverse primer  1     ATATCCCAGCATCCCGCATCATCA  24
Template        1814  ........................  1791
```

>XM_011241151.3 PREDICTED: Mus musculus adenylate cyclase activating polypeptide 1 receptor 1 (Adcyap1r1), transcript variant X3, mRNA

```
product length = 199
Forward primer  1     TTCACTACTGCGTGGTGTCCAACT  24
Template        1317  ........................  1340

Reverse primer  1     ATATCCCAGCATCCCGCATCATCA  24
Template        1515  ........................  1492
```

0

>XM_006505388.4 PREDICTED: Mus musculus adenylate cyclase activating polypeptide 1 receptor 1 (Adcyap1r1), transcript variant X2, mRNA

```
product length = 199
Forward primer  1     TTCACTACTGCGTGGTGTCCAACT  24
Template        1620  ........................  1643

Reverse primer  1     ATATCCCAGCATCCCGCATCATCA  24
Template        1818  ........................  1795
```

>XM_030255114.1 PREDICTED: Mus musculus adenylate cyclase activating polypeptide 1 receptor 1 (Adcyap1r1), transcript variant X9, mRNA

```
product length = 199
Forward primer  1     TTCACTACTGCGTGGTGTCCAACT  24
Template        1576  ........................  1599

Reverse primer  1     ATATCCCAGCATCCCGCATCATCA  24
Template        1774  ........................  1751
```

>XM_011241153.3 PREDICTED: Mus musculus adenylate cyclase activating polypeptide 1 receptor 1 (Adcyap1r1), transcript variant X8, mRNA

```
product length = 199
Forward primer  1     TTCACTACTGCGTGGTGTCCAACT  24
Template        1317  ........................  1340

Reverse primer  1     ATATCCCAGCATCCCGCATCATCA  24
Template        1515  ........................  1492
```

>XM_011241152.3 PREDICTED: Mus musculus adenylate cyclase activating polypeptide 1 receptor 1 (Adcyap1r1), transcript variant X6, mRNA

```
product length = 199
Forward primer  1     TTCACTACTGCGTGGTGTCCAACT  24
Template        1380  ........................  1403

Reverse primer  1     ATATCCCAGCATCCCGCATCATCA  24
Template        1578  ........................  1555
```

1

>XM_011241150.1 PREDICTED: Mus musculus adenylate cyclase activating polypeptide 1 receptor 1 (Adcyap1r1), transcript variant X1, mRNA

```
product length = 199
Forward primer  1     TTCACTACTGCGTGGTGTCCAACT  24
Template        1645  ........................  1668

Reverse primer  1     ATATCCCAGCATCCCGCATCATCA  24
Template        1843  ........................  1820
```

>NM_007407.4 Mus musculus adenylate cyclase activating polypeptide 1 receptor 1 (Adcyap1r1), transcript variant 1, mRNA

```
product length = 199
Forward primer  1     TTCACTACTGCGTGGTGTCCAACT  24
Template        1168  ........................  1191

Reverse primer  1     ATATCCCAGCATCCCGCATCATCA  24
Template        1366  ........................  1343
```

>NM_001025372.2 Mus musculus adenylate cyclase activating polypeptide 1 receptor 1 (Adcyap1r1), transcript variant 2, mRNA

```
product length = 199
Forward primer  1     TTCACTACTGCGTGGTGTCCAACT  24
Template        1168  ........................  1191

Reverse primer  1     ATATCCCAGCATCCCGCATCATCA  24
Template        1366  ........................  1343
```

***Adcyap1r1*** PrimerCheck (SpliceCenter der Genomics and Bioinformatics Group, LMP, CCR, NCI, http://projects.insilico.us/SpliceCenter/PrimerCheck.jsp )

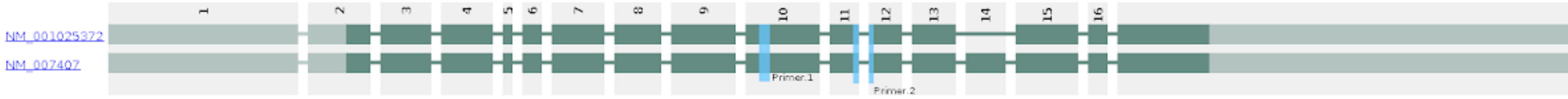

Adcyap1r1

*Adcyap1r1* Amplicon Sequence (https://www.ncbi.nlm.nih.gov/nucleotide/)

**TTCACTACTGCGTGGTGTCCAACT**ACTTCTG...AGCGTCC**TGATGATGC**CATTGCTCTTCTTGCTTTCCCCCTTAGAGGCTGGCTTT...GGTATGGACCCCCTCAGG
GAACCCAGGTTCATTCTGCCTACGCGAGGAGGAGGAAGTGGTGTTGAACCATACCCCAGCA**GGGATGCT**ATTTCTGAGAACTGCTTCCTGCAGACCGGGCAGAG
TTGCAGGGAAGAGTAGTTAGCATCAGAAAACTACCACGTTCT...AGGTTTTGGCTGGATGGCATATGGGGACTGAGGCTTTCTTGGCATGACCACCATTTACTATA
TGGAGAGCTACATGGGGAGGG**GGGATAT**

#### Structures

| Structure Name | Image | ΔG (kcal.mole$^{-1}$... | $T_M$ (°C) | ΔH (kcal.mole$^{-1}$) | ΔS (cal.K$^{-1}$mole$^{-1}$) | Output |
|---|---|---|---|---|---|---|
| 1 | | -23.85 | 43.8 | -401.2 | -1265.64 | Ct Det |
| 2 | | -23.47 | 41.9 | -437 | -1386.99 | Ct Det |
| 3 | | -22.69 | 47.7 | -321 | -1000.55 | Ct Det |
| 4 | | -22.15 | 43.1 | -386.9 | -1223.37 | Ct Det |
| 5 | | -21.48 | 42.3 | -390.9 | -1239.06 | Ct Det |

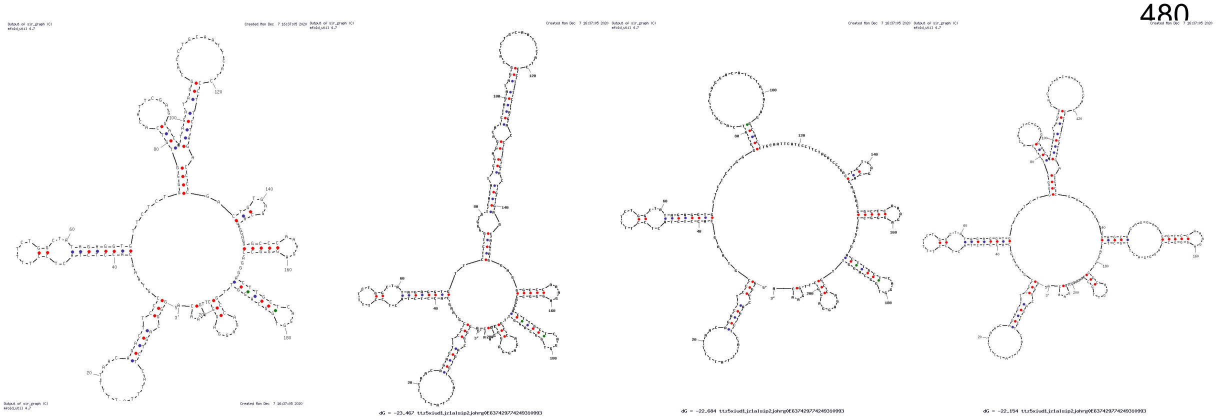

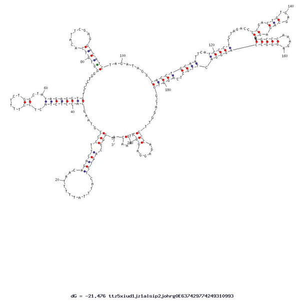

**Oxtr** PrimerBLAST (National Center for Biotechnology Information, Bethesda MD, USA, https://www.ncbi.nlm.nih.gov/tools/primer-blast )

**Primer pair 1**

| | Sequence (5'->3') | Length | Tm | GC% | Self complementarity | Self 3' complementarity |
|---|---|---|---|---|---|---|
| Forward primer | CGCACAGTGAAGATGACCTT | 20 | 58.20 | 50.00 | 5.00 | 5.00 |
| Reverse primer | ATGGCAATGATGAAGGCAGA | 20 | 57.55 | 45.00 | 4.00 | 0.00 |

**Products on target templates**

>XM_036165923.1 PREDICTED: Mus musculus oxytocin receptor (Oxtr), transcript variant X2, mRNA

```
product length = 131
Forward primer  1     CGCACAGTGAAGATGACCTT  20
Template        1343  ....................  1362

Reverse primer  1     ATGGCAATGATGAAGGCAGA  20
Template        1473  ....................  1454
```

>XM_006505723.3 PREDICTED: Mus musculus oxytocin receptor (Oxtr), transcript variant X1, mRNA

```
product length = 131
Forward primer  1     CGCACAGTGAAGATGACCTT  20
Template        1515  ....................  1534

Reverse primer  1     ATGGCAATGATGAAGGCAGA  20
Template        1645  ....................  1626
```

>NM_001081147.2 Mus musculus oxytocin receptor (Oxtr), mRNA

```
product length = 131
Forward primer  1    CGCACAGTGAAGATGACCTT  20
Template        839  ....................  858

Reverse primer  1    ATGGCAATGATGAAGGCAGA  20
Template        969  ....................  950
```

**Oxtr** PrimerCheck (SpliceCenter der Genomics and Bioinformatics Group, LMP, CCR, NCI, http://projects.insilico.us/SpliceCenter/PrimerCheck.jsp )

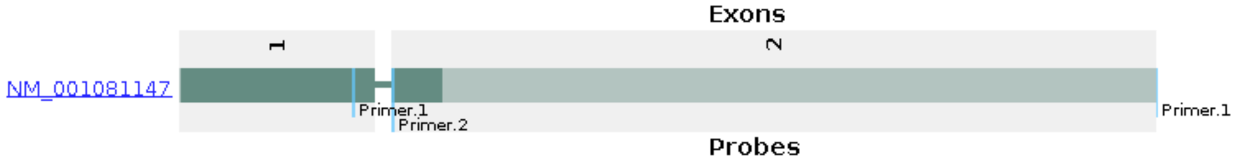

Oxtr

**Oxtr** Amplicon Sequence (https://www.ncbi.nlm.nih.gov/nucleotide/)

**CGCACAGTGAAGATGACCTT**CATCATT......CACAGCT**TCTGCCTTCATCATTGCCAT**

#### Structures

| Structure Name | Image | ΔG (kcal.mole$^{-1}$... | $T_M$ (°C) | ΔH (kcal.mole$^{-1}$) | ΔS (cal.K$^{-1}$mole$^{-1}$) | Output |
|---|---|---|---|---|---|---|
| 1 | | -33.95 | 55.4 | -366.9 | -1116.73 | Ct Det |
| 2 | | -32.85 | 53.7 | -374.4 | -1145.55 | Ct Det |
| 3 | | -32.28 | 51.4 | -396.5 | -1221.62 | Ct Det |
| 4 | | -31.97 | 52.5 | -378 | -1160.6 | Ct Det |
| 5 | | -31.6 | 52.1 | -379.2 | -1165.84 | Ct Det |
| 6 | | -30.95 | 52.6 | -365.8 | -1123.08 | Ct Det |
| 7 | | -30.48 | 50.4 | -388.5 | -1200.79 | Ct Det |
| 8 | | -30.21 | 53.9 | -341.8 | -1045.07 | Ct Det |

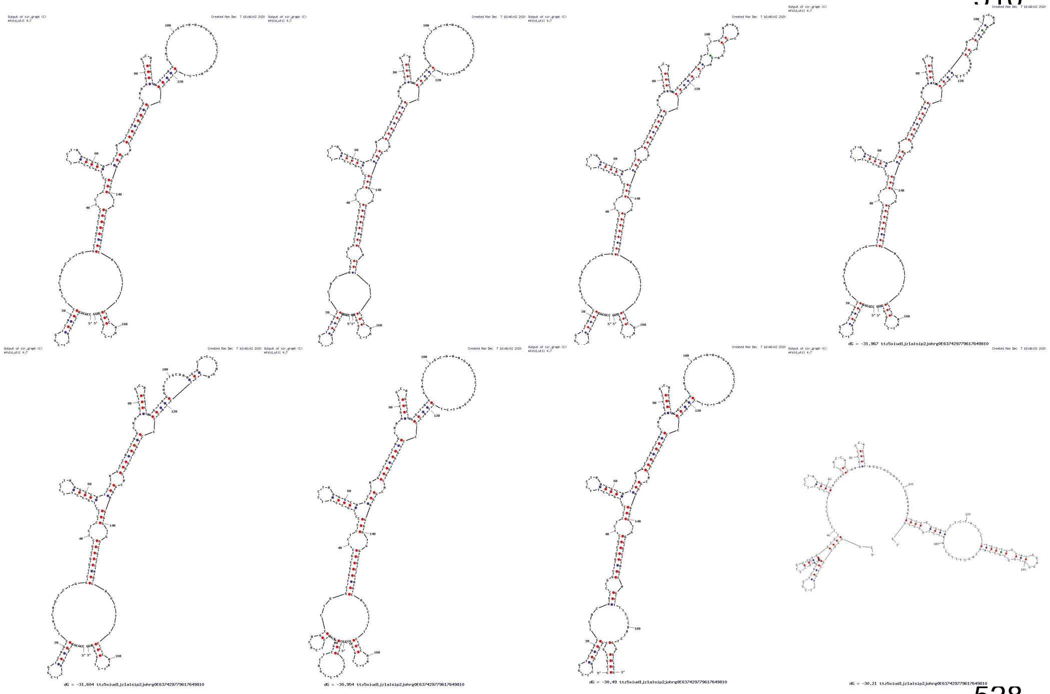

**ActB** PrimerBLAST (National Center for Biotechnology Information, Bethesda MD, USA, https://www.ncbi.nlm.nih.gov/tools/primer-blast )

**Primer pair 1**

| | Sequence (5'->3') | Length | Tm | GC% | Self complementarity | Self 3' complementarity |
|---|---|---|---|---|---|---|
| **Forward primer** | GATTACTGCTCTGGCTCCTAG | 21 | 57.34 | 52.38 | 4.00 | 4.00 |
| **Reverse primer** | GACTCATCGTACTCCTGCTTG | 21 | 58.20 | 52.38 | 4.00 | 0.00 |

**Products on target templates**

>NM_007393.5 Mus musculus actin, beta (Actb), mRNA

```
product length = 147
Forward primer  1     GATTACTGCTCTGGCTCCTAG  21
Template        1057  .....................  1077

Reverse primer  1     GACTCATCGTACTCCTGCTTG  21
Template        1203  .....................  1183
```

>NM_177093.3 Mus musculus leucine rich repeat containing 58 (Lrrc58), mRNA

```
product length = 146
Forward primer  1     GATTACTGCTCTGGCTCCTAG  21
Template        3191  .....................  3211

Reverse primer  1     GACTCATCGTACTCCTGCTTG  21
Template        3336  ........C............  3316
```

**ActB** PrimerCheck (SpliceCenter der Genomics and Bioinformatics Group, LMP, CCR, NCI, http://projects.insilico.us/SpliceCenter/PrimerCheck.jsp )

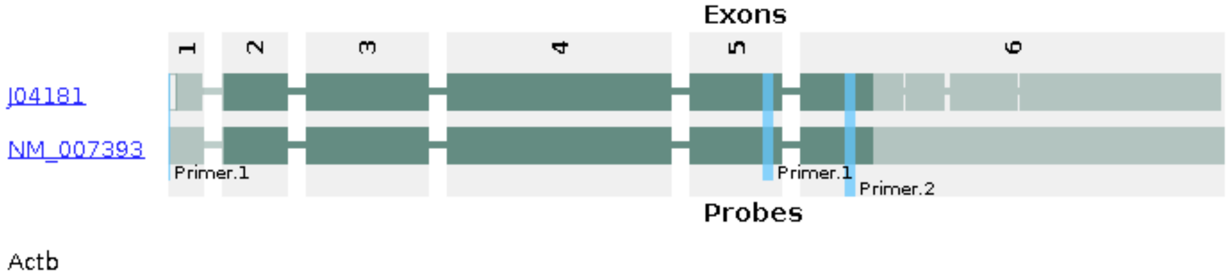

**ActB** Amplicon Sequence (https://www.ncbi.nlm.nih.gov/nucleotide/) :

**<u>GATTACTGCTCTGGCTCCTAG</u>**CACCATGAAGATCAAGGTAAGCTAAGCATCCTTAGCTTGGTGAGGGTGGGCCCTGTGGTTGTCAGAGCAACCTTCTAGGTTTA AGGGGAATCCCAGCACCCAGAGAGCTCACCATTCACCATCTTGTCTTGCTTTCTTCAGATCATTGCTCCTCCTGAGCGCAAGTACTCTGTGTGGATCGGTGGCTCC ATCCTGGCCTCACTGTCCACCTTCCAGCAGATGTGGATCAG**<u>CAAGCAGGAGTACGATGAGTC</u>**

#### Structures

| Structure Name | Image | ΔG (kcal.mole$^{-1}$... | $T_M$ (°C) | ΔH (kcal.mole$^{-1}$) | ΔS (cal.K$^{-1}$mole$^{-1}$) | Output |
|---|---|---|---|---|---|---|
| 1 | 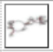 | -30.23 | 48.9 | -407.8 | -1266.38 | Ct Det |
| 2 | 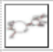 | -29.51 | 47.4 | -422.4 | -1317.76 | Ct Det |
| 3 | 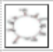 | -29.19 | 50.9 | -365.6 | -1128.34 | Ct Det |
| 4 | 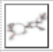 | -29.13 | 47.4 | -416.1 | -1297.9 | Ct Det |
| 5 | 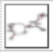 | -29.04 | 46.3 | -435.1 | -1361.94 | Ct Det |
| 6 | 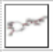 | -28.67 | 46.6 | -424.9 | -1328.96 | Ct Det |
| 7 | 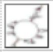 | -28.65 | 48.5 | -392.2 | -1219.34 | Ct Det |
| 8 | 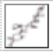 | -28.56 | 44.7 | -461.7 | -1452.76 | Ct Det |

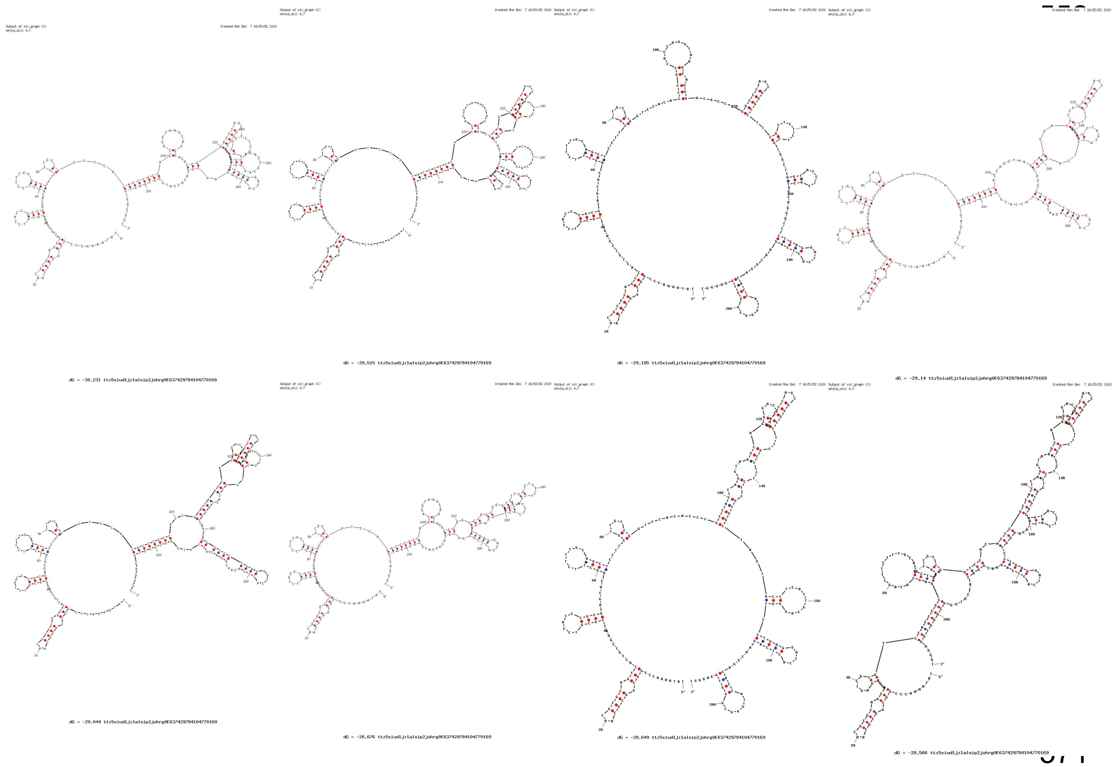

**Supplementary Data 2. Amplification plots and Melting curve analysis (RT-qPCR).**

**qPCR programs (used in qPCR runs with 55C and 60C anneal temperatures)**

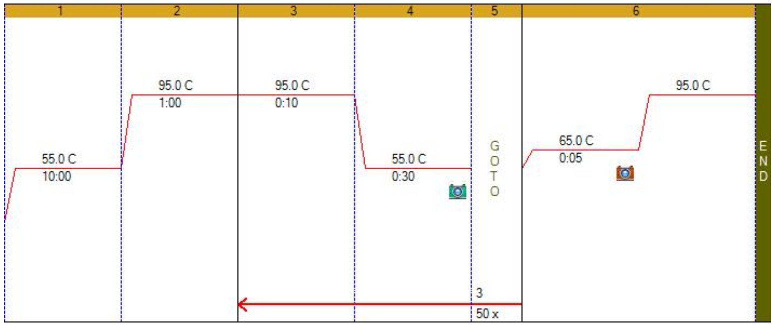

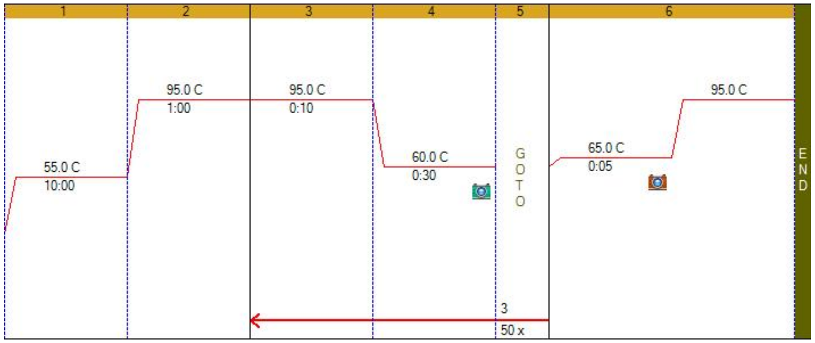

**Avp1ar (efficiency qPCR – standard curve)**

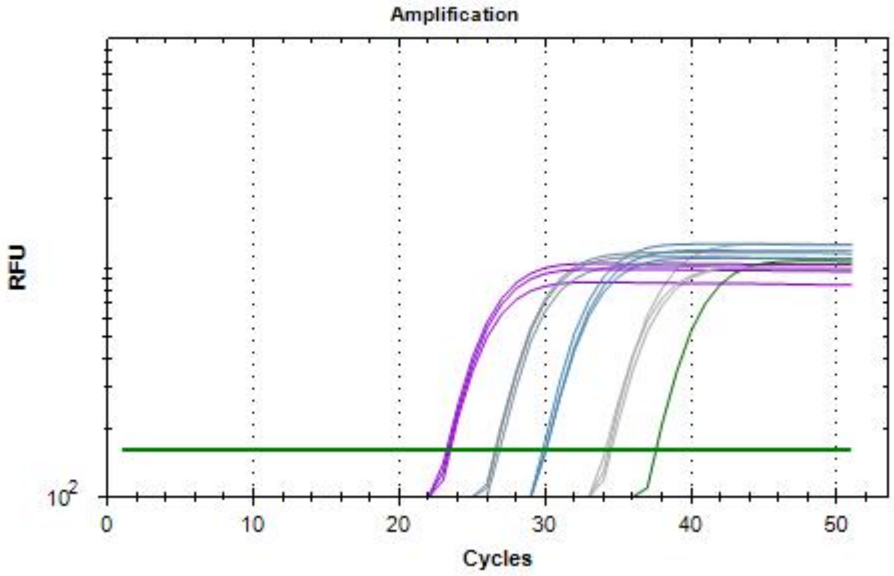

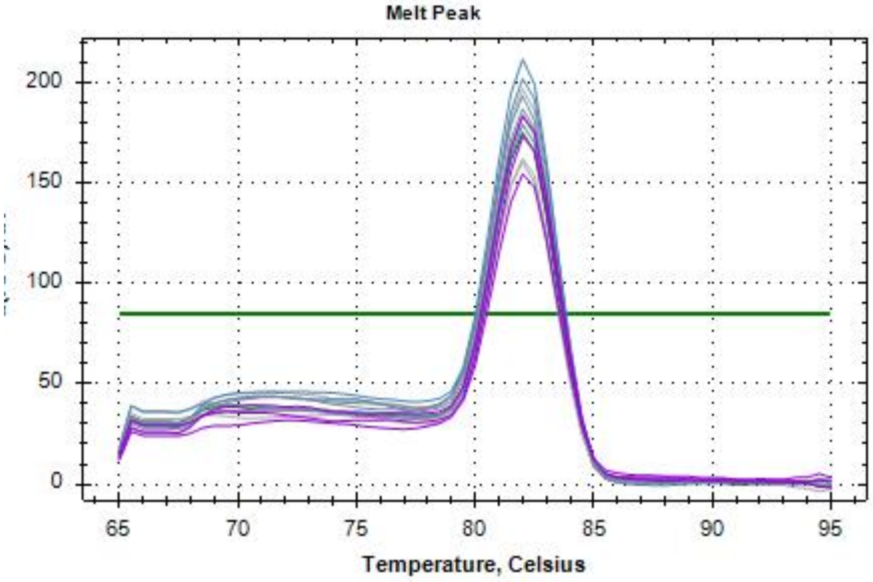

**Oxt (efficiency qPCR – standard curve)**

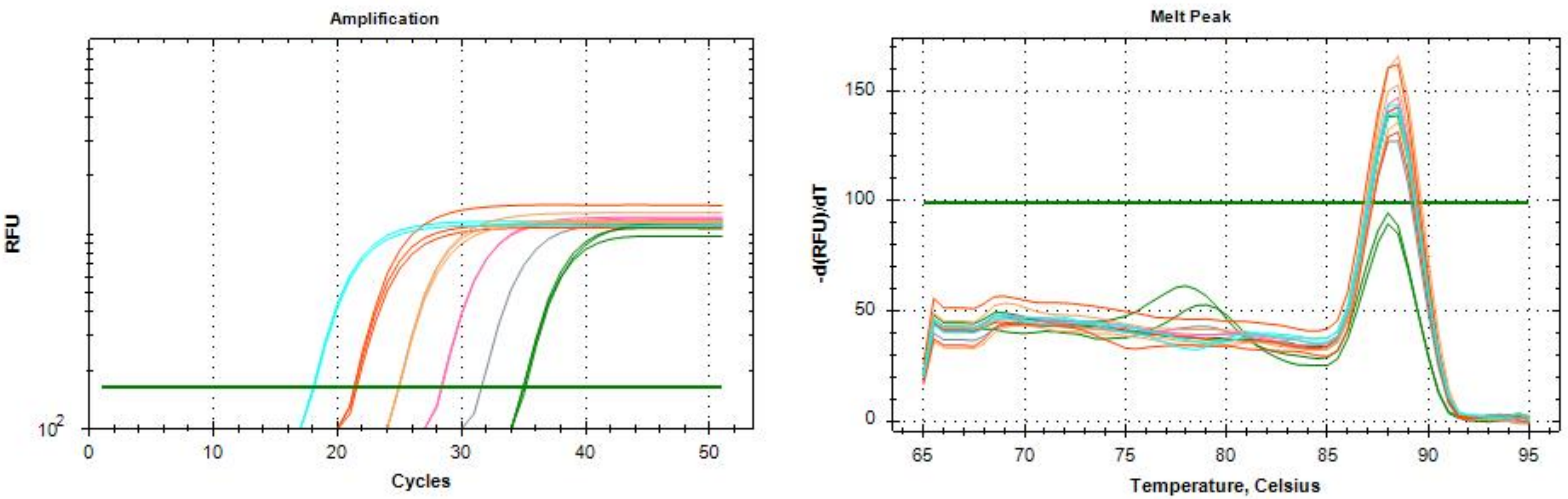

**AVP (efficiency qPCR – standard curve)**

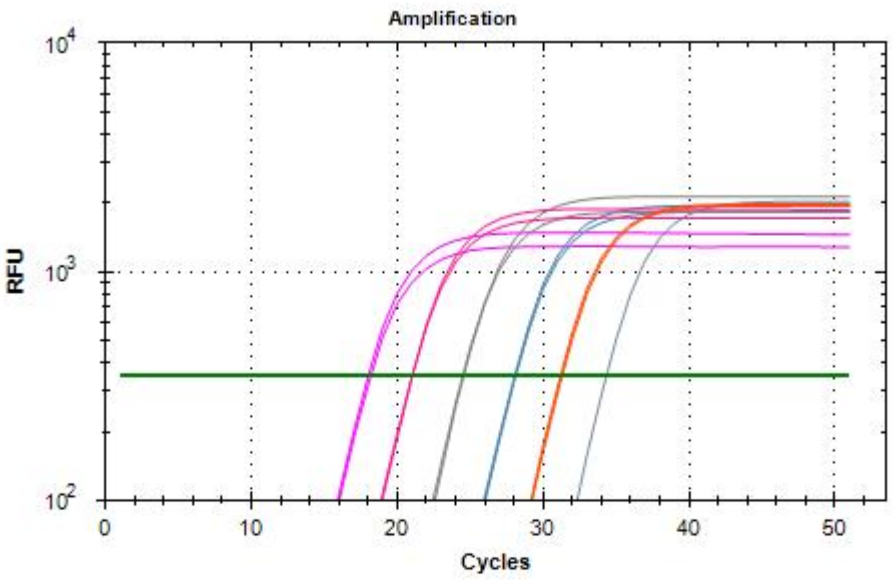
Amplification
RFU
$10^4$
$10^3$
$10^2$
0
10
20
30
40
50
Cycles

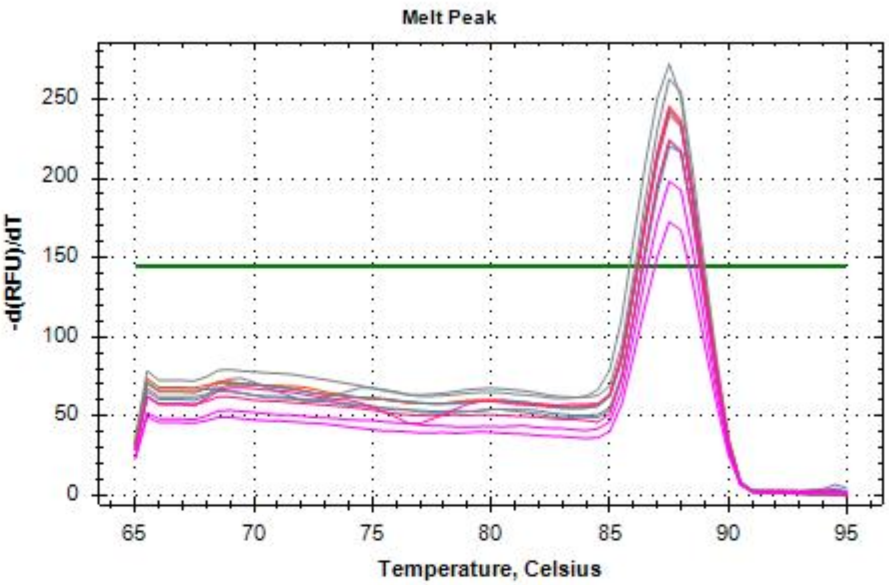
Melt Peak
-d(RFU)/dT
250
200
150
100
50
0
65
70
75
80
85
90
95
Temperature, Celsius

**PACAP (efficiency qPCR – standard curve)**

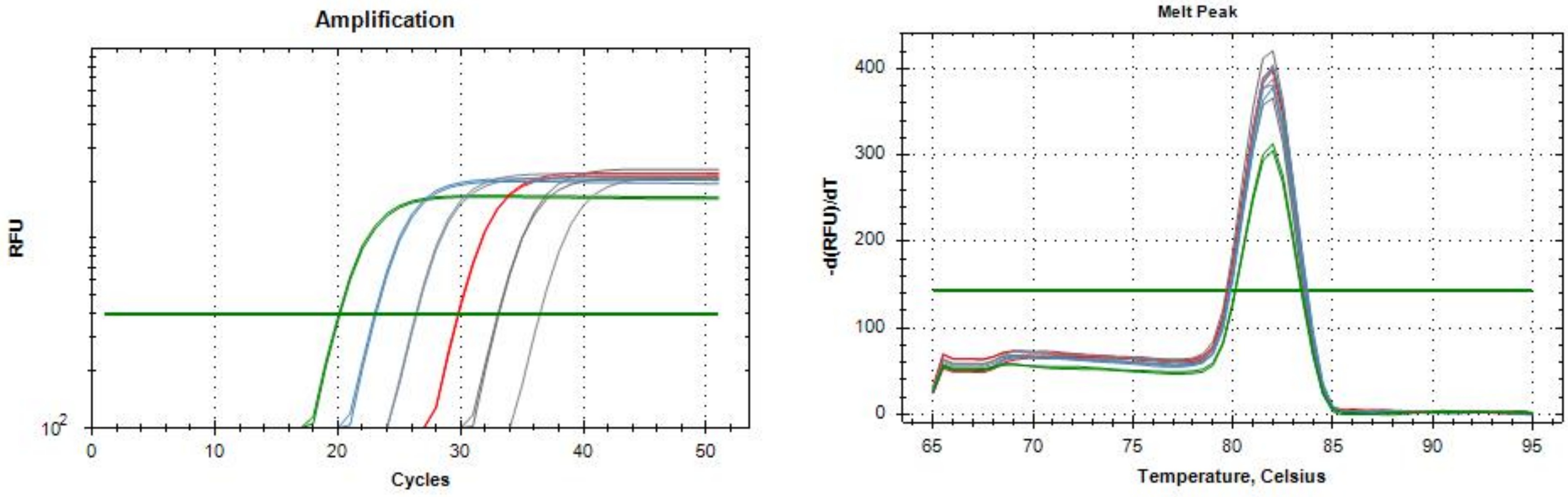

*Adcyap1r1* (efficiency qPCR – standard curve)

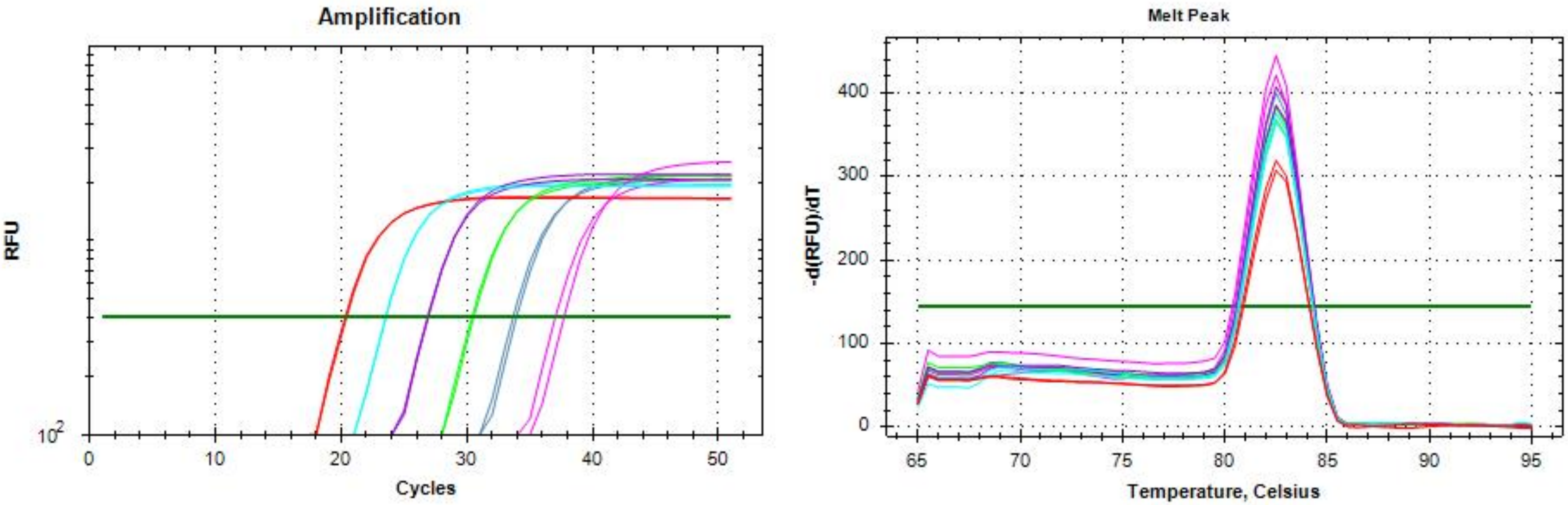

#### ActB (efficiency qPCR – standard curve)

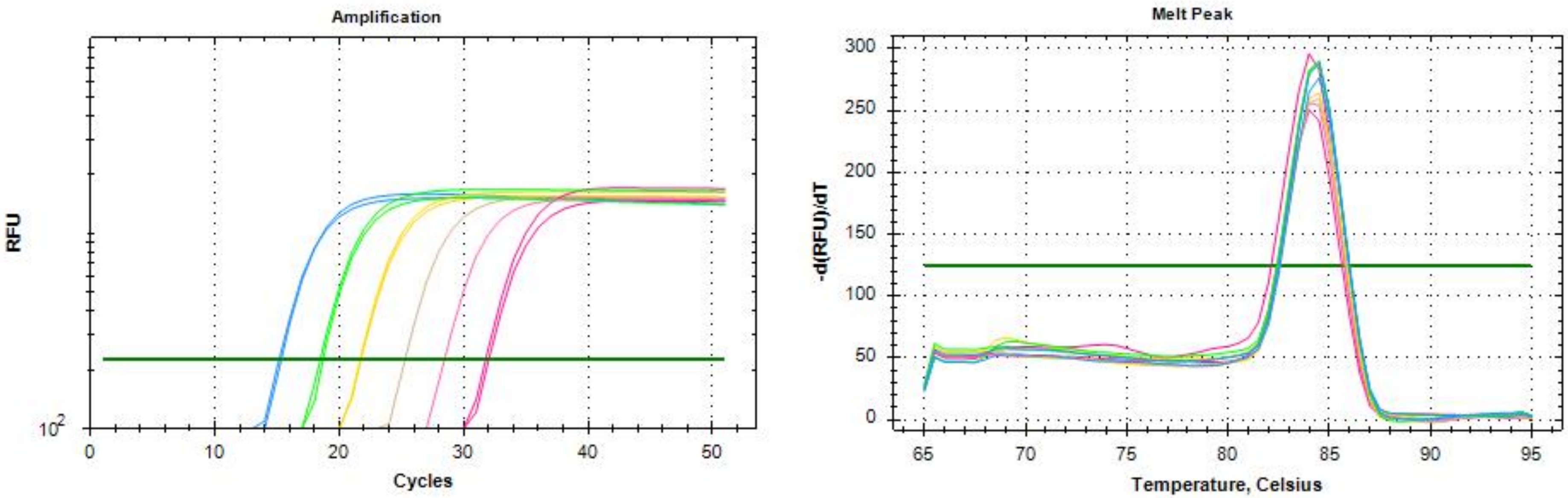

**Oxtr Hydrolysis Probe (efficiency qPCR – standard curve)**

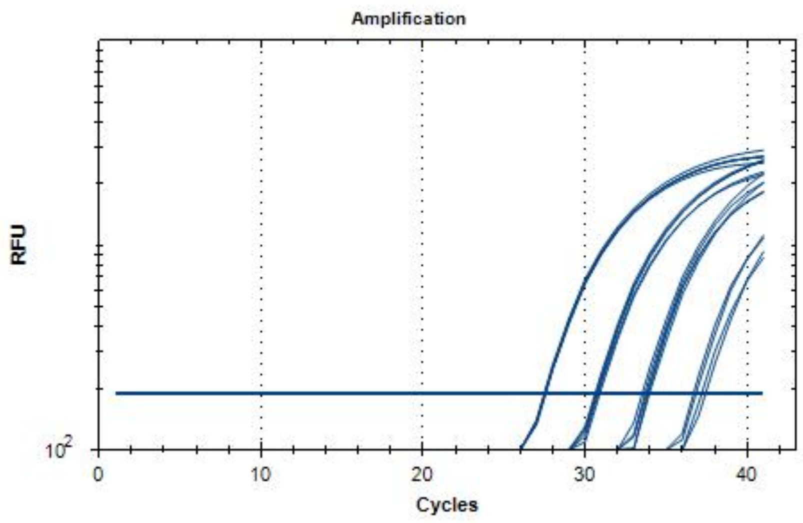

**ActB Hydrolysis Probe (efficiency qPCR – standard curve)**

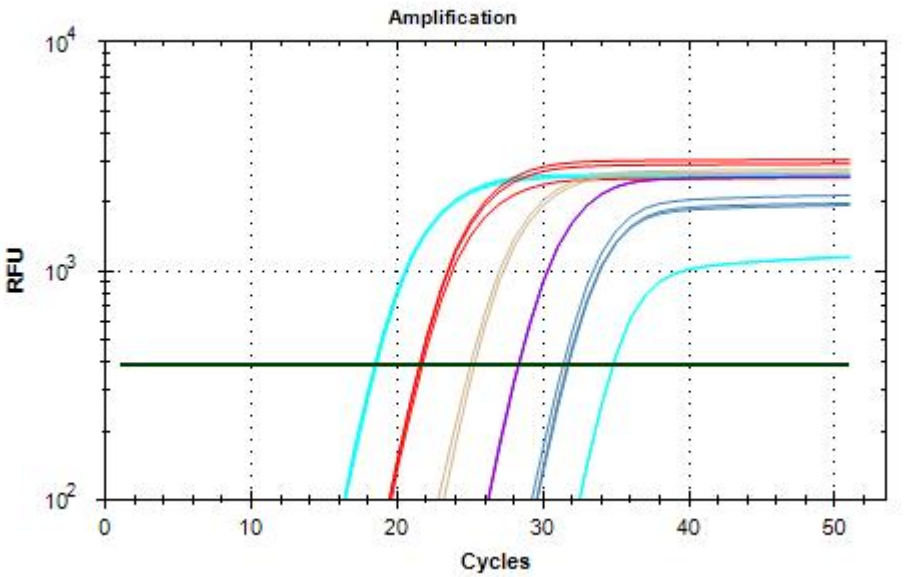

**Supplementary Data 3. Evaluation of qPCR Primer Efficiency**

**<u>ActB hydrolysis probe</u>** primer efficiency

| Gene | RNA equivalent (pg) | Cq Triplet | Cq 95%CI | Cq Mean | Cq SD | RNA dilution |
|---|---|---|---|---|---|---|
| ActB Probe<br>ActB Probe<br>ActB Probe | 100000<br>100000<br>100000 | 18.29<br>18.23<br>18.37 | 18.12/18.47 | 18.3 | 0.07 | 1:10 |
| ActB Probe<br>ActB Probe<br>ActB Probe | 10000<br>10000<br>10000 | 21.34<br>21.31<br>21.5 | 21.13/21.64 | 21.38 | 0.102 | $1{:}10^2$ |
| ActB Probe<br>ActB Probe<br>ActB Probe | 1000<br>1000<br>1000 | 24.82<br>25.1<br>24.5 | 24.06/25.55 | 24.81 | 0.298 | $1{:}10^3$ |
| ActB Probe<br>ActB Probe<br>ActB Probe | 100<br>100<br>100 | 28.11<br>28.15 | 27.88/28.38 | 31 | .028 | $1{:}10^4$ |
| ActB Probe<br>ActB Probe<br>ActB Probe | 10<br>10<br>10 | 31.52<br>31.44<br>31.18 | 30.94/31.82 | 31.38 | 0.182 | $1{:}10^5$ |
| ActB Probe<br>ActB Probe<br>ActB Probe | 1<br>1<br>1 | 35.34<br>34.58<br>34.57 | 33.73/35.93 | 34.83 | 0.44 | $1{:}10^6$ |
| ActB Probe | NTC | ND | | | | |

NTC, no template control; ND, not detected

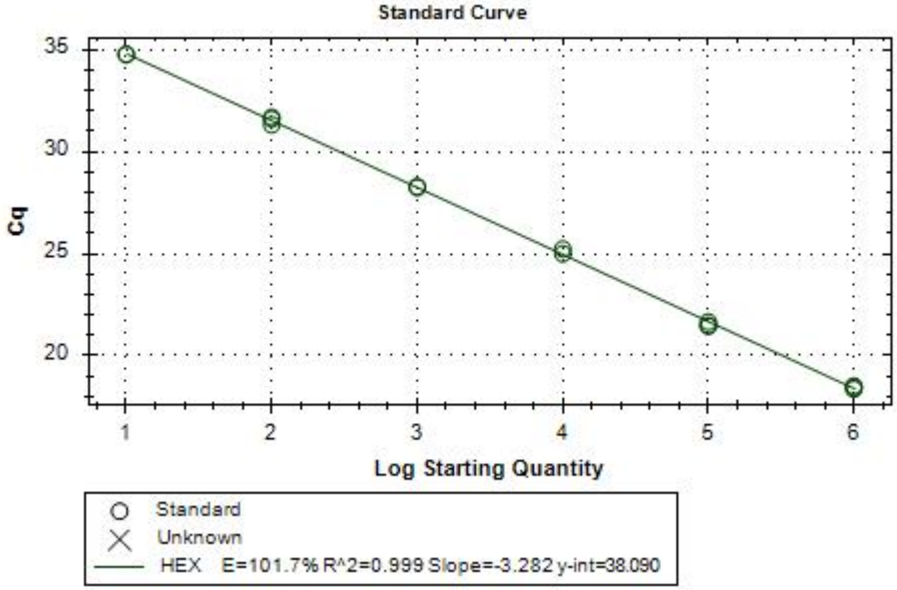

**LDR (**dilution range**):** 1:10-1:$10^6$
**LOD (**dilution**):** ≤1:$10^6$

SD = standard deviation; NTC = no-template control; LDR = linear dynamic range; LOD = limit of detection; R^2 = coefficient of determination; CI = confidence interval

**Avp1ar** primer efficiency

| Gene | RNA equivalent (pg) | Cq Triplet | Cq 95%CI | Cq Mean | Cq SD | RNA dilution |
|---|---|---|---|---|---|---|
| Avp1ar<br>Avp1ar<br>Avp1ar | 100000<br>100000<br>100000 | 23.38<br>23.27<br>23.16 | 23.00/23.54 | 23.27 | 0.112 | 1:10 |
| Avp1ar<br>Avp1ar<br>Avp1ar | 10000<br>10000<br>10000 | 26.83<br>26.46<br>26.53 | 26.12/27.09 | 26.6 | 0.199 | $1:10^2$ |
| Avp1ar<br>Avp1ar<br>Avp1ar | 1000<br>1000<br>1000 | 30<br>30.01<br>29.77 | 29.59/30.26 | 29.93 | 0.137 | $1:10^3$ |
| Avp1ar<br>Avp1ar<br>Avp1ar | 100<br>100<br>100 | 34.08<br>34.41<br>34.25 | 33.84/34.66 | 34.25 | 0.169 | $1:10^4$ |
| Avp1ar<br>Avp1ar<br>Avp1ar | 10<br>10<br>10 | NA<br>37.49<br>38.1 | 33.92/41.67 | 37.79 | 0.432 | $1:10^5$ |
| Avp1ar<br>Avp1ar<br>Avp1ar | 1<br>1<br>1 | 39.18<br>NA<br>46.48 | -3.55/89.21 | 42.83 | 5.161 | $1:10^6$ |
| Avp1ar | NTC | ND | | | | |

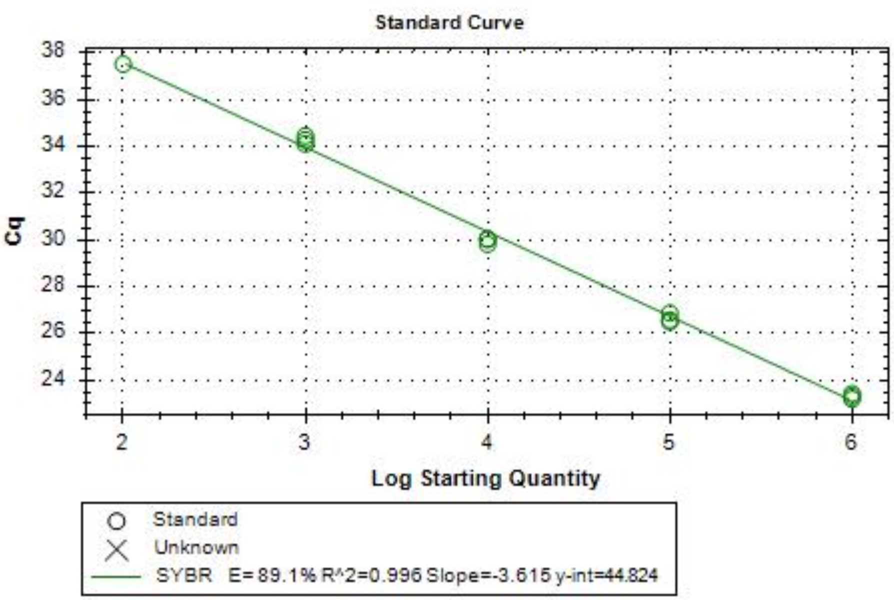

**LDR** (dilution range)**:** 1:10-1:$10^5$
**LOD** (dilution)**:** ≤1:$10^6$

SD = standard deviation; NTC = no-template control; LDR = linear dynamic range; LOD = limit of detection; R^2 = coefficient of determination; CI = confidence interval

**Oxt** primer efficiency

| Gene | RNA equivalent (pg) | Cq Triplet | Cq 95%CI | Cq Mean | Cq SD | RNA dilution |
|---|---|---|---|---|---|---|
| Oxt<br>Oxt | 100000<br>100000 | 18.09<br>18.02 | 17.61/18.50 | 18.06 | 0.049 | 1:10 |
| Oxt<br>Oxt<br>Oxt | 10000<br>10000<br>10000 | 21.47<br>21.32<br>21.28 | 21.11/21.61 | 21.36 | 0.100 | $1:10^2$ |
| Oxt<br>Oxt<br>Oxt | 1000<br>1000<br>1000 | 24.95<br>24.93<br>24.89 | 24.85/25.00 | 24.92 | 0.031 | $1:10^3$ |
| Oxt<br>Oxt | 100<br>100 | 28.29<br>28.35 | 27.94/28.70 | 28.32 | 0.042 | $1:10^4$ |
| Oxt | 10 | 31.53 | NA | 31.53 | NA | $1:10^5$ |
| Oxt<br>Oxt<br>Oxt | 1<br>1<br>1 | 34.99<br>34.97<br>35.12 | 34.82/35.23 | 35.03 | 0.081 | $1:10^6$ |
| Oxt | NTC | ND | | | | |

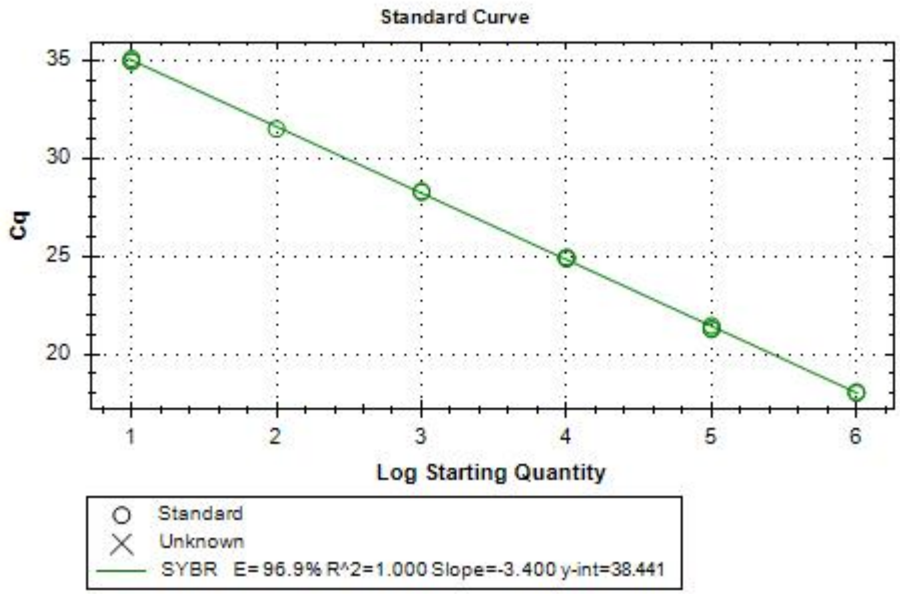

**LDR (**dilution range**):** 1:10-1:$10^6$
**LOD (**dilution**):** ≤1:$10^6$

SD = standard deviation; NTC = no-template control; LDR = linear dynamic range; LOD = limit of detection; R^2 = coefficient of determination; CI = confidence interval

**AVP** primer efficiency

| Gene | RNA equivalent (pg) | Cq Triplet | Cq 95%CI | Cq Mean | Cq SD | RNA dilution |
|---|---|---|---|---|---|---|
| AVP<br>AVP | 100000<br>100000 | 18.18<br>18.02 | 17.08/19.12 | 18.1 | 0.111 | 1:10 |
| AVP<br>AVP | 10000<br>10000 | 21.03<br>21.05 | 20.91/21.17 | 21.04 | 0.013 | $1:10^2$ |
| AVP<br>AVP | 1000<br>1000 | 24.44<br>24.5 | 24.09/24.85 | 24.47 | 0.047 | $1:10^3$ |
| AVP<br>AVP | 100<br>100 | 28.12<br>28.05 | 27.64/28.53 | 28.09 | 0.044 | $1:10^4$ |
| AVP<br>AVP | 10<br>10 | 31.17<br>31.27 | 30.58/31.86 | 31.22 | 0.07 | $1:10^5$ |
| AVP<br>AVP | 1<br>1 | 35.43<br>34.3 | 27.69/42.04 | 34.87 | 0.803 | $1:10^6$ |
| AVP | NTC | ND | | | | |

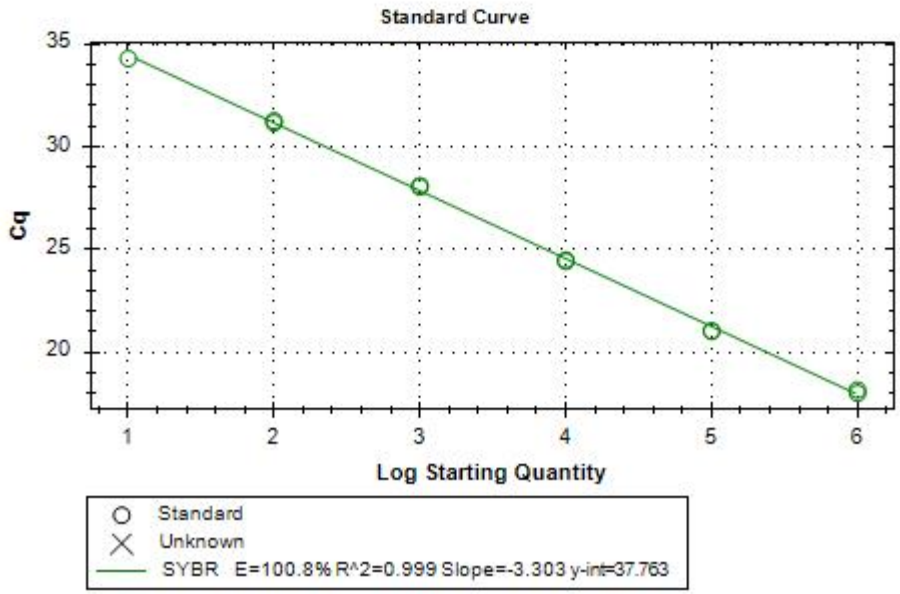

**LDR (**dilution range**):** 1:10-1:$10^6$
**LOD (**dilution**):** ≤1:$10^6$

SD = standard deviation; NTC = no-template control; LDR = linear dynamic range; LOD = limit of detection; R^2 = coefficient of determination; CI = confidence interval

Adcyap1 primer efficiency

| **Gene** | **RNA equivalent (pg)** | **Cq Replicate** | **Cq 95%CI** | **Cq Mean** | **Cq SD** | **RNA dilution** |
|---|---|---|---|---|---|---|
| Adcyap1<br>Adcyap1 | 100000<br>100000 | 20.04<br>20.12 | 19.57/20.59 | 20.08 | 0.058 | 1:10 |
| Adcyap1<br>Adcyap1 | 10000<br>10000 | 22.95<br>23.07 | 22.25/23.77 | 23.01 | 0.084 | $1:10^{2}$ |
| Adcyap1<br>Adcyap1 | 1000<br>1000 | 26.27<br>26.34 | 25.86/26.75 | 26.31 | 0.048 | $1:10^{3}$ |
| Adcyap1<br>Adcyap1 | 100<br>100 | 29.78<br>29.71 | 29.30/30.19 | 29.75 | 0.051 | $1:10^{4}$ |
| Adcyap1<br>Adcyap1 | 10<br>10 | 33.01<br>33.07 | 32.66/33.42 | 33.04 | 0.043 | $1:10^{5}$ |
| Adcyap1<br>Adcyap1 | 1<br>1 | 38.34<br>36.38 | 24.91/49.81 | 37.36 | 1.386 | $1:10^{6}$ |
| Adcyap1 | NTC | 38.48 | | | | |

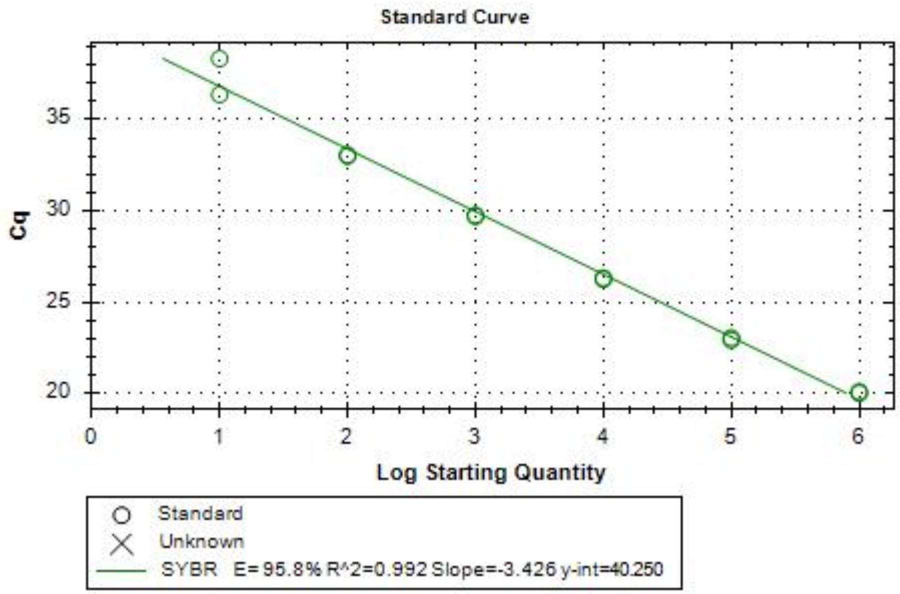

**LDR (dilution range):** 1:10-1:$10^5$
**LOD (dilution):** ≤1:$10^6$

SD = standard deviation; NTC = no-template control; LDR = linear dynamic range; LOD = limit of detection; R^2 = coefficient of determination; CI = confidence interval

*Adcyap1r1* primer efficiency

| **Gene** | **RNA equivalent (pg)** | **Cq Replicate** | **Cq 95%CI** | **Cq Mean** | **Cq SD** | **RNA dilution** |
|---|---|---|---|---|---|---|
| Adcyapr1r<br>Adcyapr1r | 100000<br>100000 | 20.27<br>20.37 | 19.68/20.96 | 20.32 | 0.071 | 1:10 |
| Adcyapr1r<br>Adcyapr1r | 10000<br>10000 | 23.47<br>23.49 | 23.35/23.61 | 23.48 | 0.011 | 1:10$^2$ |
| Adcyapr1r<br>Adcyapr1r | 1000<br>1000 | 26.83<br>26.9 | 26.42/27.31 | 26.86 | 0.049 | 1:10$^3$ |
| Adcyapr1r<br>Adcyapr1r | 100<br>100 | 30.34<br>30.48 | 29.52/31.30 | 30.41 | 0.097 | 1:10$^4$ |
| Adcyapr1r<br>Adcyapr1r | 10<br>10 | 33.61<br>33.91 | 31.85/35.67 | 33.76 | 0.214 | 1:10$^5$ |
| Adcyapr1r<br>Adcyapr1r | 1<br>1 | 37.02<br>37.66 | 33.27/41.41 | 37.34 | 0.453 | 1:10$^6$ |
| Adcyapr1r | NTC | 41.23 | | | | |

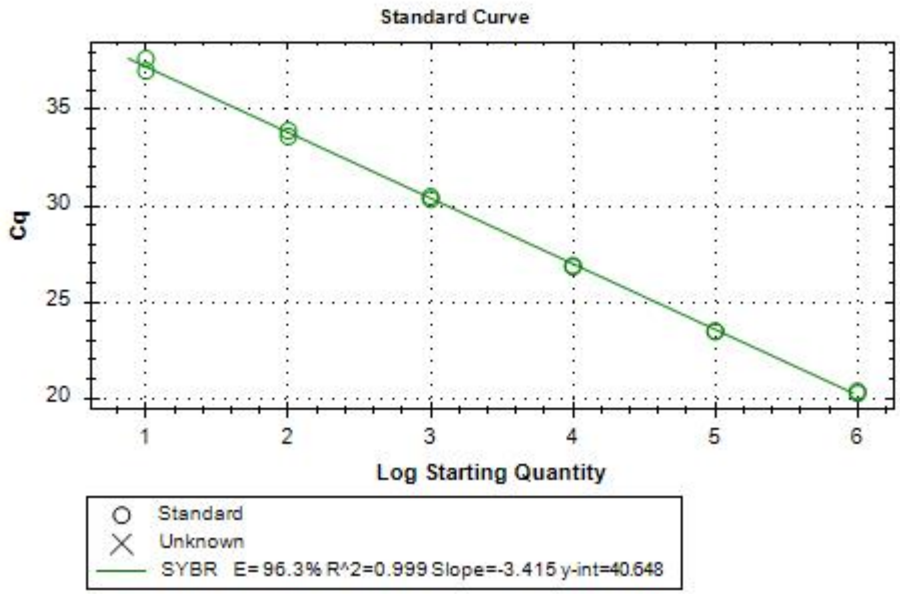

**LDR (**dilution range**):** 1:10-1:$10^6$
**LOD (**dilution**):** ≤1:$10^6$

SD = standard deviation; NTC = no-template control; LDR = linear dynamic range; LOD = limit of detection; R^2 = coefficient of determination; CI = confidence interval

Act B primer efficiency

| Gene | RNA equivalent (pg) | Cq Replicate | Cq 95%CI | Cq Mean | Cq SD | RNA dilution |
|---|---|---|---|---|---|---|
| ActB<br>ActB | 100000<br>100000 | 15.22<br>15.36 | 14.40/16.18 | 15.29 | 0.097 | 1:10 |
| ActB<br>ActB | 10000<br>10000 | 18.5<br>18.72 | 17.21/20.01 | 18.61 | 0.151 | $1:10^{2}$ |
| ActB<br>ActB | 1000<br>1000 | 21.8<br>21.73 | 21.32/22.21 | 21.77 | 0.05 | $1:10^{3}$ |
| ActB<br>ActB | 100<br>100 | 25.33<br>NA | NA | 25.33 | NA | $1:10^{4}$ |
| ActB<br>ActB | 10<br>10 | 28.5<br>24.88 | 3.692/49.69 | 26.69 | 2.56 | $1:10^{5}$ |
| ActB<br>ActB | 1<br>1 | 32.1<br>31.84 | 30.32/33.62 | 31.97 | 0.184 | $1:10^{6}$ |
| ActB | NTC | 39.77 | | | | |

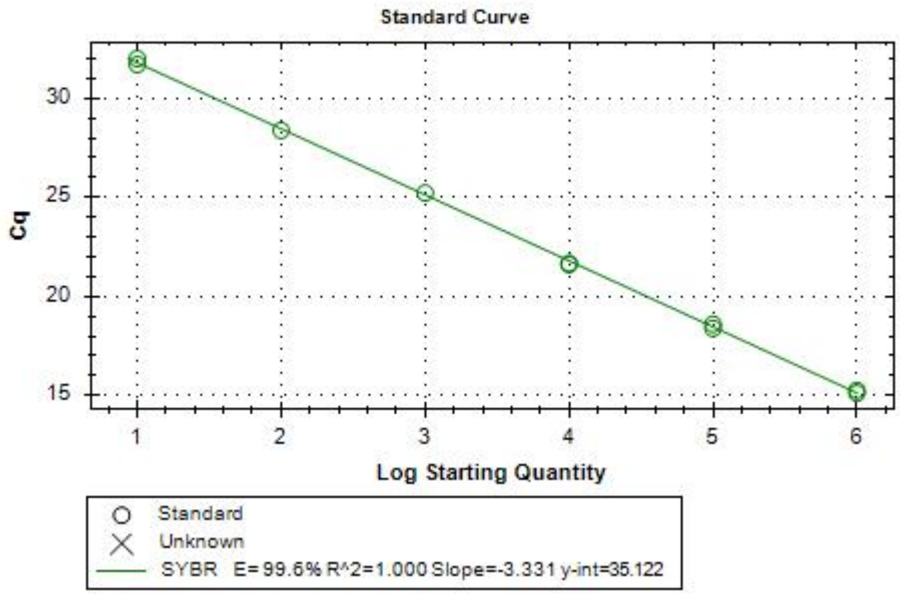

**LDR (**dilution range**):** 1:10-1:$10^4$
**LOD (**dilution**):** ≤1:$10^6$

SD = standard deviation; NTC = no-template control; LDR = linear dynamic range; LOD = limit of detection; R^2 = coefficient of determination; CI = confidence interval

Oxtr hydrolysis probe primer efficiency

| Gene | RNA equivalent (pg) | Cq Replicate | Cq 95%CI | Cq Mean | Cq SD | RNA dilution |
|---|---|---|---|---|---|---|
| Oxtr | 100000 | 27.1 | 27.08/27.14 | 27.11 | 0.024 | 1:10 |
| Oxtr | 100000 | 27.1 | | | | |
| Oxtr | 100000 | 27.12 | | | | |
| Oxtr | 100000 | 27.14 | | | | |
| Oxtr | 100000 | 27.08 | | | | |
| Oxtr | 10000 | 30.34 | 30.17/30.42 | 30.29 | 0.102 | 1:10$^{2}$ |
| Oxtr | 10000 | 30.44 | | | | |
| Oxtr | 10000 | 30.25 | | | | |
| Oxtr | 10000 | 30.25 | | | | |
| Oxtr | 10000 | 30.18 | | | | |
| Oxtr | 1000 | 33.35 | 33.15/33.54 | 33.35 | 0.160 | 1:10$^{3}$ |
| Oxtr | 1000 | 33.47 | | | | |
| Oxtr | 1000 | 33.12 | | | | |
| Oxtr | 1000 | 33.52 | | | | |
| Oxtr | 1000 | 33.27 | | | | |
| Oxtr | 100 | 37.56 | 36.16/37.45 | 36.8 | 0.519 | 1:10$^{4}$ |
| Oxtr | 100 | 36.77 | | | | |
| Oxtr | 100 | 37.02 | | | | |
| Oxtr | 100 | 36.27 | | | | |
| Oxtr | 100 | 36.39 | | | | |
| Oxtr | 10 | 38.5 | 37.15/39.75 | 38.45 | 1.047 | 1:10$^{5}$ |
| Oxtr | 10 | 38.99 | | | | |
| Oxtr | 10 | 39.09 | | | | |
| Oxtr | 10 | 36.63 | | | | |
| Oxtr | 10 | 39.06 | | | | |

| | | | | | | |
|---|---|---|---|---|---|---|
| Oxtr<br>Oxtr<br>Oxtr<br>Oxtr<br>Oxtr | 1<br>1<br>1<br>1<br>1 | NA<br>38.25<br>NA<br>NA<br>38.2 | 37.91/38.54 | 38.23 | 0.035 | 1:$10^6$ |
| Oxtr | NTC | 39.04 | | | | |

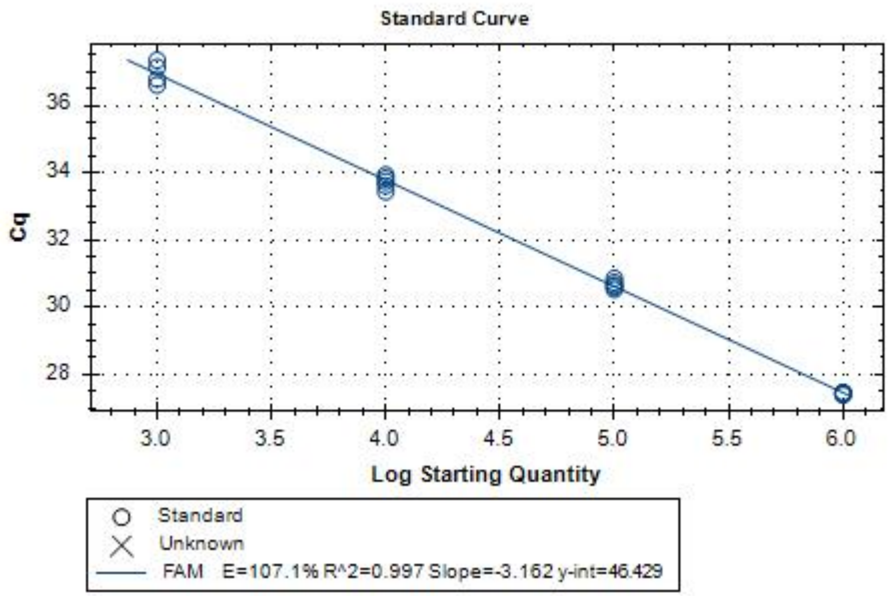

**LDR (**dilution range**):** 1:10-1:$10^5$
**LOD** (dilution)**:** ≤1:$10^6$

SD = standard deviation; NTC = no-template control; LDR = linear dynamic range; LOD = limit of detection; R^2 = coefficient of determination; CI = confidence interval
